# Dynamic metabolic exchanges between complementary bacterial types provide collaborative stress resistance

**DOI:** 10.1101/2021.06.24.449802

**Authors:** Kapil Amarnath, Avaneesh V. Narla, Sammy Pontrelli, Jiajia Dong, Brian R. Taylor, Tolga Caglar, Julia Schwartzman, Uwe Sauer, Otto X. Cordero, Terence Hwa

## Abstract

Metabolic cross-feeding plays vital roles in promoting ecological diversity. While some microbes depend on exchanges of essential nutrients for growth, forces driving the extensive cross-feeding needed to support the coexistence of free-living microbes are poorly understood. Here we characterize bacterial physiology under self-acidification, and establish that extensive excretion of key metabolites following acidification provides a collaborative, inter-species mechanism of stress resistance. This collaboration occurs not only between species isolated from the same community, but also between unrelated species with complementary (glycolytic vs. gluconeogenic) modes of metabolism. Cultures of such communities cycle through different phases in growth-dilution experiments, comprising of exponential growth, growth arrest upon acidification, collaborative stress relief, and growth recovery, with each phase involving distinct physiological states of individual species. Our findings challenge the static view of ecosystems commonly portrayed in ecological models, and offer an alternative dynamical view based on growth advantages of different species in different phases.

---

Metabolic cross-feeding underlies many positive interactions between microbes^1–3^. Many well-studied examples of cross-feeding involve species that are *dependent* on each other for essential metabolic functions, including synthetic complementary auxotrophy^4–13^ and designated cross-feeding between symbionts^14,15^. The driving force for metabolic cooperation between such inter-dependent bacteria is clear, since they lack the ability to generate essential metabolites themselves and must obtain them from other species in order to grow.

Many bacteria in nature, however, are prototrophic, or “free living” – that is, they can grow on simple substrates without the help of others^16,17^. Free-living bacteria can be forced to excrete large amounts of metabolites (e.g., at mM/OD level needed to support substantial biomass accumulation by other organisms) via genetic manipulation, the design and attainment of which is an important goal of synthetic biology^18–20^. However, naturally occurring free bacteria are generally not known to excrete such large amounts of their endogenous metabolites during normal growth, except for a few well-studied cases. Such cases include overflow metabolism during aerobic fermentation^21–23^, the excretion of nitrate/nitrite during anaerobic denitrification^24,25^, and complex cascades of fermentation product removal in anaerobic digesters^26,27^. Yet, recent studies indicate that substantial exchanges of diverse metabolites support many species of naturally occurring free-living bacteria in synthetic bacterial communities growing aerobically on simple substrates^28–30^. This suggests that there are additional important driving forces underlying the seemingly altruistic cross-feeding between free-living bacteria that we currently know little about.

In this study, we reveal physiological driving forces for substantial metabolic excretion by naturally occurring, free-living bacteria. We show that excretion of mM/OD level of valuable metabolites such as pyruvate naturally occurs under stress, and that the excreted metabolites are required for other species in the community to grow and relieve stress, ultimately rescuing the excreting species from death and restoring the community. This collaborative inter-species stress relief mechanism can occur between species taken from vastly different environments, indicating that it is not a result of selection in specific environments. Instead, this interaction is attributed to a fundamental complementarity between free-living bacteria with opposing modes of metabolism, with both modes needed to overcome stress.

The stress under study here arises from the accumulation of weak acid, e.g., acetate, which is commonly encountered in many environments, from the gut to bioreactors^31–39^. Weak acids are excreted during anaerobic growth^40^, but also aerobically under iron limitation^41^, and even in favorable growth conditions^42,43^. The excreted acids become toxic when the environment becomes acidified, i.e., when pH drops to the level of the acids’ dissociation constants, ∼5 for weak acids such as acetate ^44,45^. Under these conditions, the ionic form of the acid builds to high concentrations in the cytoplasm, reducing the pools of key metabolic intermediates due to a osmotic constraint^46^. While the presence of “acid eaters” specializing on acids in non-stress conditions can theoretically prevent acidification, we find that such acid eaters are themselves growth-inhibited when the pH drops. We reveal that during acid stress, an additional layer of metabolic exchange occurs transiently between the growth-inhibited acid excreters and acid eaters - ‘acid-induced cross-feeding’ - wherein large amounts of additional fermentation products such as pyruvate and lactate are converted from the external nutrients and released into the medium by the non-growing acid excreters to restore the growth of the acid eaters, ultimately allowing the latter to consume all of the acids and thereby detoxifying the environment for both types of species. We find this acid-induced metabolic exchange to occur readily between different pairs of glycolytically-oriented, sugar-consuming acid excreters and gluconeogenically oriented acid eaters, reflecting an underlying collaborative mechanism between bacterial species with complementary modes of metabolism. Through quantitative, systematic investigation, we will first describe acid-induced cross-feeding in a case of rapid acetate accumulation during the aerobic growth of a co-culture of marine bacteria. We will then show that the same process of stress relief occurs in co-cultures comprising of soil and enteric bacteria, and for different mechanisms of acetate accumulation.

## Results

### Simple acetate cross-feeding between two co-isolated bacteria in strong buffer

This study started with the characterization of *Vibrio splendidus* sp. 1A01 and *Neptunomonas phycotrophica* sp. 3B05, two species co-isolated from a chitin enrichment culture of coastal ocean water^47^, to study simple cross-feeding between two natural, free living strains of bacteria. When cultured alone on chitin, 1A01 grew but 3B05 did not (**Extended Data Fig. 1a**). To investigate possible reasons for the presence of 3B05 and many other non-chitin-degrading bacteria in the enrichment culture^47^, we grew these two strains together using N-acetyl-glucosamine (GlcNAc), the monomer of chitin, as the sole carbon and nitrogen source, in defined, minimal medium strongly buffered at pH = 8, the nominal pH of sea water^48^; see **Materials and Methods**. After inoculating 1A01 and 3B05 at equal ratio, the co-culture was left to grow for 24 hours, then diluted 40-fold into fresh medium. Such 24-h growth-dilution cycles were repeated for several days (**Fig. 1a**). Before each dilution, the abundance of each species was monitored using 16S PCR^49^ (**Extended Data Fig. 1b**). The two species were found to coexist stably, settling after a few cycles to a ratio 3B05:1A01≈1:3 by 16S abundance (**Fig. 1b**, black diamonds).

**Figure 1.**
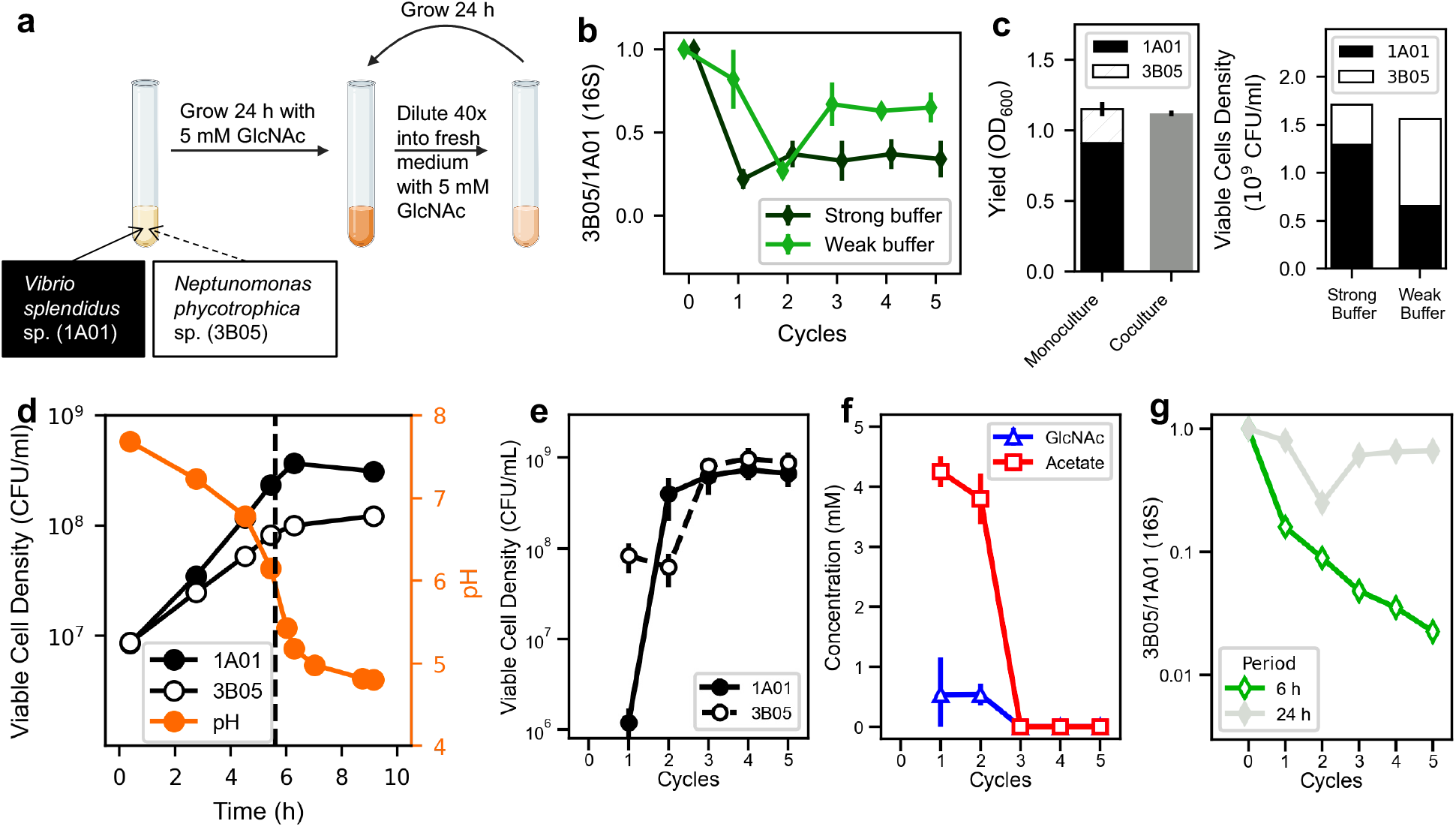
Coexistence of 1A01 and 3B05 in growth-dilution cycles. **(a)** *Vibrio splendidus* sp. 1A01 and *Neptunomonas phycotrophica* sp. 3B05 were cocultured in growth-dilution cycles with 5 mM GlcNAc as the sole carbon and nitrogen source, with a 24-h cycle and 40x dilution after each cycle. **(b)** Ratio of 3B05 to 1A01 based on 16S reads at the end of each cycle, starting from 1:1 mixture of exponentially growing cells each at OD_600_=0.01 (for the remainder of the manuscript we abbreviate OD_600_ as OD). Results from experiments done in strong buffer (40 mM HEPES) are shown in black, and those from experiments done in weak buffer (2 mM NaHCO_3_) are shown as green symbols. The error bars indicate the standard error from 3 biological replicates. **(c)** Left panel: Solid left bar indicates the monoculture yield of 1A01 (in OD_600_) on 5 mM GlcNAc. Open left bar indicates the monoculture yield of 3B05 on 7.4 mM acetate; the latter is the amount of acetate excreted by 1A01 after growth on 5 mM GlcNAc (**Extended Data Fig. 1d**). Grey right bar indicates the measured yield of the 1A01-3B05 co-culture on 5 mM GlcNAc. Data in the left panel were derived from exponentially growing co-culture in the strong buffer. The error bars indicate the standard error from 3 biological replicates, which was larger than the measurement variability of ±0.002 OD unit on the basis of repeated measurements of the same sample of culture. Right panel: composition of the co-culture based on viable cell count after five 24-h growth-dilution cycle in the strong 40 mM HEPES buffer (left bars) and in the weak 2 mM bicarbonate buffer (right bar). **(d)** Viable cell density of 1A01 (filled circles) and 3B05 (open circles) in a co-culture in the weak 2 mM bicarbonate buffer inoculated at 1:1 initial ratio. The vertical dashed line indicates the time where biomass growth ceased (defined in **Extended Data Fig. 3a**). **(e)** Densities of viable 1A01 cells (filled circles) and 3B05 cells (open circles), and **(f)** concentrations of GlcNAc (blue triangles) and acetate (red squares) in the medium, measured at the end of each 24-h cycle for the co-culture grown in weak buffer. The error bars indicate the standard error from 3 biological replicates. **(g)** Ratio of 3B05 to 1A01 based on 16S reads, collected at the end of each cycle of a 6-h growth-dilution experiments in the weak buffer (open green diamonds). These values were independently confirmed by plating. Results for the 24-h cycle (filled grey diamonds) are reproduced from panel b for comparison.

To determine the mechanistic basis of the coexistence of these two species on GlcNAc, we quantified the growth and uptake/excretion characteristics of each species in monoculture. Only 1A01 grew in monoculture with GlcNAc as the sole carbon and nitrogen source (**Extended Data Fig. 1c**). We analyzed the culture medium using HPLC columns (readily detecting > 10 μM of common carbohydrates and amino acids; see **Materials and Methods**) and found substantial accumulation of acetate and ammonium (**Extended Data Fig. 1d**). A closer examination of the excretion data (summarized in **Extended Data Fig. 1e**) suggests that the acetate liberated in the conversion of GlcNAc to glucosamine^50^ was directly released into the medium, in addition to the acetate released due to overflow metabolism during rapid growth on glucose^42^.

The substantial excretion of acetate and ammonium by 1A01 in monoculture suggested that 3B05 might be growing on these carbon and nitrogen sources in the coculture. As a first test, we grew 1A01 and 3B05 as monocultures on acetate and ammonium and found that only 3B05 grew (open black squares, **Extended Data Fig. 1f**). The results suggest that a simple commensalism between 1A01 and 3B05 underlies the coexistence found in **Fig. 1b**. Indeed, the yields of both species in the co-culture can be quantitatively explained (left panel, **Fig. 1c)** by the single-strain characteristics summarized in **Extended Data Table 1** that were measured during exponential, steady-state growth.

### Acetate cross-feeding insufficient to support coexistence in weak buffer

One important feature of the coculture described above is the high buffer capacity (40 mM HEPES) used, which fixed the medium pH despite acetate accumulation and enabled us to focus solely on nutrient consumption and cross-feeding. A scenario of broad ecological relevance is one in which the medium is acidified by the excreted acids, as many natural environments including the ocean are weakly buffered^51–53^, and acidification (i.e., pH drop) can easily occur when the excreted acid reaches the order of the buffer capacity in the environment. Thus, in the ocean which is buffered by ∼2 mM bicarbonate (primarily from equilibration with atmospheric CO_2_ ^53^), acidification would occur when the excreted acetate reaches ∼2 mM. As bacterial growth is generally inhibited at reduced pH, especially in the presence of weak acids such as acetate^54–56^, and the presence of acid-eaters such as 3B05 would alleviate acidification, the relationship between 1A01 and 3B05 changes from a commensal one for the co-culture in a strong buffer to a syntrophic one in a weak buffer; see illustration in **Extended Data Fig. 2a**. Assuming that 3B05 is less affected by reduced pH than 1A01 in a weakly buffered co-culture, the growth of 3B05 on acetate would limit the acetate buildup and hence the medium pH, resulting in a canonical syntrophy scenario in which the two species grow exponentially at the pH where the growth rate of the two species matches; see **Extended Data Fig. 2b**.

However, when we grew 1A01 and 3B05 together in 2 mM bicarbonate (the buffer in the ocean, primarily from equilibration with atmospheric CO_2_ ^53^), the co-culture stopped growing after ∼6 hours (black squares, **Extended Data Fig. 3a**), reaching a final OD which is less than half of that reached in the strong buffer (horizontal dotted line). This is consistent with our analysis of the medium, which found GlcNAc dropping to only about half of the starting concentration at the time of growth arrest (blue triangles, **Extended Data Fig. 3b**). We also found accumulation of acetate in the medium (red squares), exceeding the buffer capacity (2 mM) at around 6-h where pH started dropping rapidly (orange circles, **Extended Data Fig. 3a**). Interestingly, GlcNAc concentration continued to decrease and acetate continued to increase after OD stopped increasing after 6 hours, suggesting residue metabolic activity in the non-growing co-culture.

To see why 3B05 was unable to prevent acetate build-up as depicted in the classic syntrophy scenario (**Extended Data Fig. 2a, 2b**), we characterized the densities of viable 1A01 and 3B05 cells using plating (**Extended Data Fig. 3c-e**). Our data shows that both species stopped growing as the pH started plummeting (**Fig. 1d**). We tested for the steady-state growth of these species individually at various fixed pH and found 3B05 to be more sensitive to reduced pH than 1A01 (**Extended Data Fig. 2c**), contrary to the scenario of **Extended Data Fig. 2b** canonically assumed^57^. Thus, given that 1A01 grows faster on GlcNAc than the rate 3B05 grows on acetate (**Extended Data Fig. 1c, 1f**, respectively), acetate accumulation in the medium and the resulting pH drop and growth arrest of both species become inevitable. The dynamics of the co-culture observed here are quantitatively captured by a simple metabolic model (**Extended Data Fig. 2d, 2e**)^58^, using single-strain characteristics obtained from the two monocultures without *ad-hoc* parameter fitting; see **Supplementary Note 1**.

Because 3B05 grew less than 1A01 during the 6-h period prior to the pH drop (**Fig. 1d**), we expected it to be depleted from the co-culture if the growth-dilution experiment of **Fig. 1a** was repeated in the weak buffer. Contrary to our expectation, however, coexistence remained, as measured by 16S ratio (green diamonds, **Fig. 1b**) and by cell count (**Fig. 1e**), and the co-culture settled to a stable state composition favoring 3B05 more than that of the strong buffer case (right panel, **Fig. 1c**). Moreover, measurements of GlcNAc and acetate concentration in the medium at the end of each cycle showed complete consumption of GlcNAc with no acetate accumulation once the co-culture stabilized after a few cycles (**Fig. 1f**). To look for possible syntrophic interaction that might have escaped our analysis, we repeated the growth-dilution experiment with 6-h cycles to maintain the co-culture in exponential growth, mimicking a rapidly diluting chemostat (since the co-culture grew exponentially the first 6 hours, see **Fig. 1d**). 3B05 is seen to deplete rapidly as expected (**Fig. 1g**, open diamonds). Thus, the coexistence observed in the 24-h growth-dilution experiment resulted from some syntrophic effect that occurred outside of the exponential growth phase.

Additionally, because 1A01 experienced rapid cell death following the depletion of GlcNAc (**Extended Data Fig. 3f, 3g**), we examined the possibility that the coexistence in the 24-h cycles arose from preferential death of 1A01^59–62^. However, adding death to our metabolic model could not account for the coexistence observed, because death-mediated coexistence would have a very skewed species ratio and would also leave many nutrients unconsumed (**Supplementary Note 2**), both of which are ruled out by our data (**Fig. 1e, 1f**).

### Growth and metabolite dynamics in the stable cycle

To find the mechanisms enabling both coexistence and full consumption of carbon, we analyzed the dynamics of the co-culture in the “stable cycle”, several cycles after the initial inoculation when the levels of the two species and the carbon concentrations stabilized (**Fig. 1e, 1f**). We measured the viability of 1A01 and 3B05 (**Fig. 2a**) and concentrations of GlcNAc and acetate (**Fig. 2b**) at various time during the stable cycle. The dynamics observed were strikingly different from that in the first 24-h: First, acetate in the medium was high only briefly in the middle of the stable cycle (red squares, **Fig. 2b**), with all of it consumed shortly after 1A01 stopped growing. Given the prolonged exposure to high acetate in the first 24-h and the rapid death of 1A01 in high acetate (**Extended Data Fig. 3b, 3f, 3g**), the maintenance of 1A01 viability in the stable cycle can be attributed to the rapid disappearance of acetate. Next, the growth of 3B05 (open circles, **Fig. 2a**) surged when acetate was being depleted (shaded region), even though the co-culture including 3B05 stopped growing when acetate accumulated to similar levels (∼2.5 mM) in the first cycle (**Extended Data Fig. 3b, 3f**). Moreover, at the beginning of the stable cycle, 1A01 did not grow despite the absence of acetate and the availability of GlcNAc in the fresh medium; yet 3B05 managed to grow in that same condition. These puzzles are addressed below by analyzing monocultures in conditions mimicking various phases of the stable cycle. The results will reveal how acetate is removed and species coexistence is maintained in the stable cycle.

**Figure 2.**
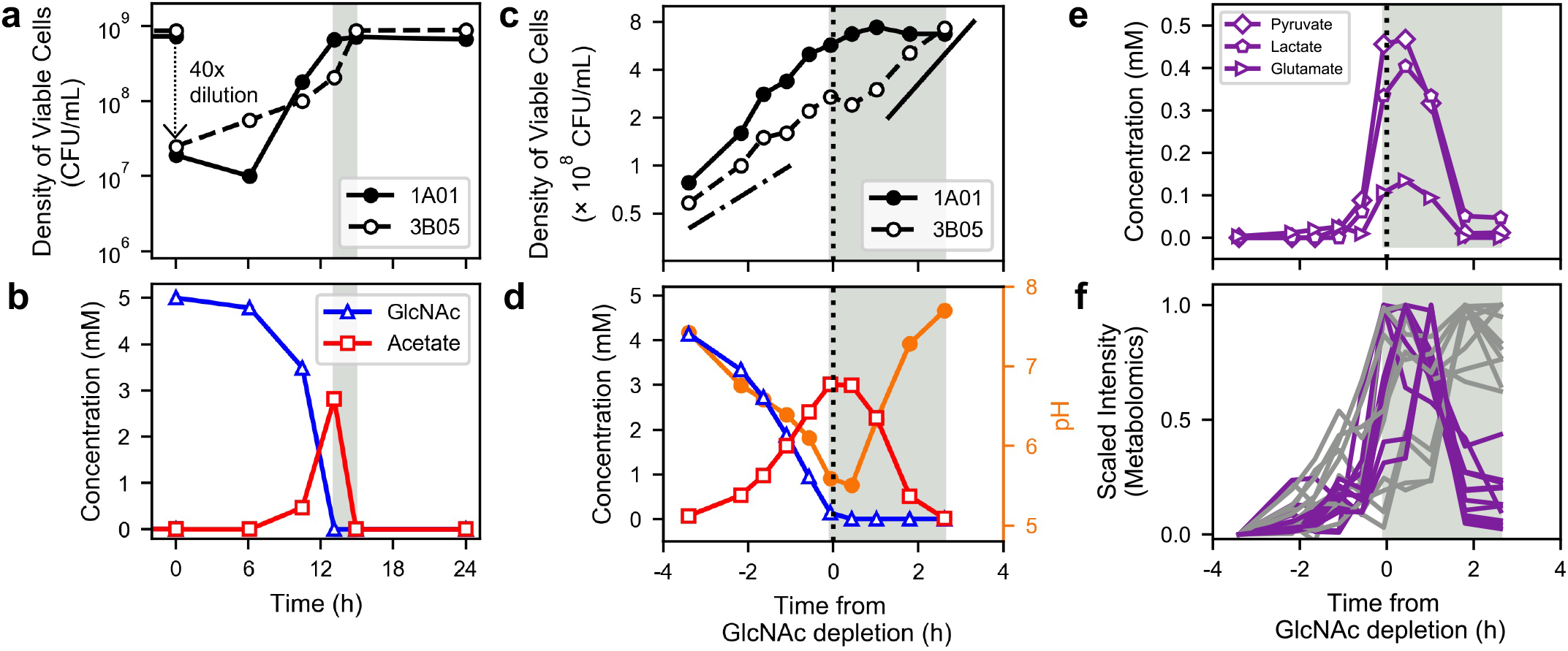
Cross-feeding in the stable cycle of the coculture in weak buffer. Measurements of various quantities of the coculture throughout the fifth 24-h growth-dilution cycle. **(a)** The viable counts of 1A01 and 3B05 cells. The value shown was the average of three measurements on the same sample from a single co-culture; the error between these measurements was less than the size of the data marker. **(b)** The concentrations of GlcNAc and Acetate in the medium from the same co-culture measured in (a). The measurement variability for the determination of sugar, organic acid, or amino acids concentrations by HPLC was ∼2% on the basis of repeated measurements of the same spent media sample. The light grey regions in (a) and (b) indicate the period when 3B05 continued to grow after GlcNAc depletion. In **(c)-(f)**, the duration around the ‘acetate peak’ was densely sampled using a protocol that mimicked the stable cycle; see Methods. The data from all four panels were measured on the same coculture. The pH measurement was accurate to ±0.02 pH unit on the basis of repeated measurements of pH standards. The dotted line at time “0” indicates the time of GlcNAc depletion, around 12 h into the cycle (panel b). Grey shaded regions are the same as those in panels (a) and (b). Same symbols are used in (c) and (d) as in (a) and (b). In (c) the dashed-dotted line indicates exponential growth of 3B05 at rate ∼0.35/hr before the acetate peak; the solid line indicates a growth rate ∼0.55/hr after the acetate peak. The filled orange circles in (d) indicate the culture pH (right vertical axis). **(e)** Concentrations of pyruvate, lactate, and glutamate in the medium as measured by HPLC. **(f)** Scaled intensities of metabolites in the medium as measured by untargeted metabolomics; see Methods. Metabolites consumed (defined as those with the scaled intensity of the last time point < 0.5) are plotted in purple. Other detected metabolites are plotted in grey. Identities of the metabolites are shown in **Extended Data Table 2**.

### Growth of 3B05 after the arrest of 1A01 was aided by metabolites excreted by 1A01 during acid stress

To determine the cause of the surge in 3B05 towards the end of co-culture growth in the stable cycle, we measured cell viability and analyzed the spent medium at many time points during the period when acetate peaked (**Fig. 2c-f**). The dense sampling revealed that the growth of both species dropped as acetate accumulated, driving pH below 6 (red squares and orange circles, **Fig. 2d**). This growth inhibition is referred to here as “acetate stress” or more generally as “acid stress”. Unlike the first 24-h (**Extended Data Fig. 3a, 3b**), the growth arrest in the stable cycle coincided approximately with the complete exhaustion of GlcNAc.

Analysis of the medium by HPLC revealed that, in addition to acetate, several other metabolites, namely pyruvate, lactate, and glutamate, accumulated to high concentrations starting from 30-60 min before GlcNAc was exhausted (**Fig. 2e**). The ensuing disappearance of these metabolites (grey shaded region) coincided with the depletion of acetate and the recovery of the pH (**Fig. 2d**), and the surge of 3B05 growth (open circles, **Fig. 2c**), while the density of 1A01 remained constant during this period (filled circles, **Fig. 2c**). During its surge, 3B05 grew at a rate substantially larger than on acetate alone (compare solid and dash-dotted lines in **Fig. 2c**). This faster growth rate is consistent with the growth rate of 3B05 on a mixture of acetate, lactate, pyruvate, and glutamate at normal pH (**Extended Data Fig. 1f**, triangles), suggesting that the surge of 3B05 was aided by the internal metabolites in the medium. The consumption of these internal metabolites in addition to acetate would also account for the higher composition of 3B05 reached in the stable cycle in weak buffer compared to that in strong buffer (right panel, **Fig. 1c**).

To understand how 3B05 managed to grow during the surge period when the pH was low, while it did not grow below a pH of 5.7 in the monoculture (open circles, **Extended Data Fig. 2c**), we grew the 3B05 monoculture in medium acidified by acetate, with and without the supplement of lactate, pyruvate, and glutamate, the metabolites which accumulated significantly during the surge (**Fig. 2e**). 3B05 only grew with the supplement (filled squares, **Fig. 3a**), which resulted in the uptake of both acetate and the supplements (filled squares, **Fig. 3b**), accompanied by pH recovery (filled circles, **Fig. 3c**). Thus, these supplements relieved the cause of growth inhibition when 3B05 was with acetate alone.

**Figure 3.**
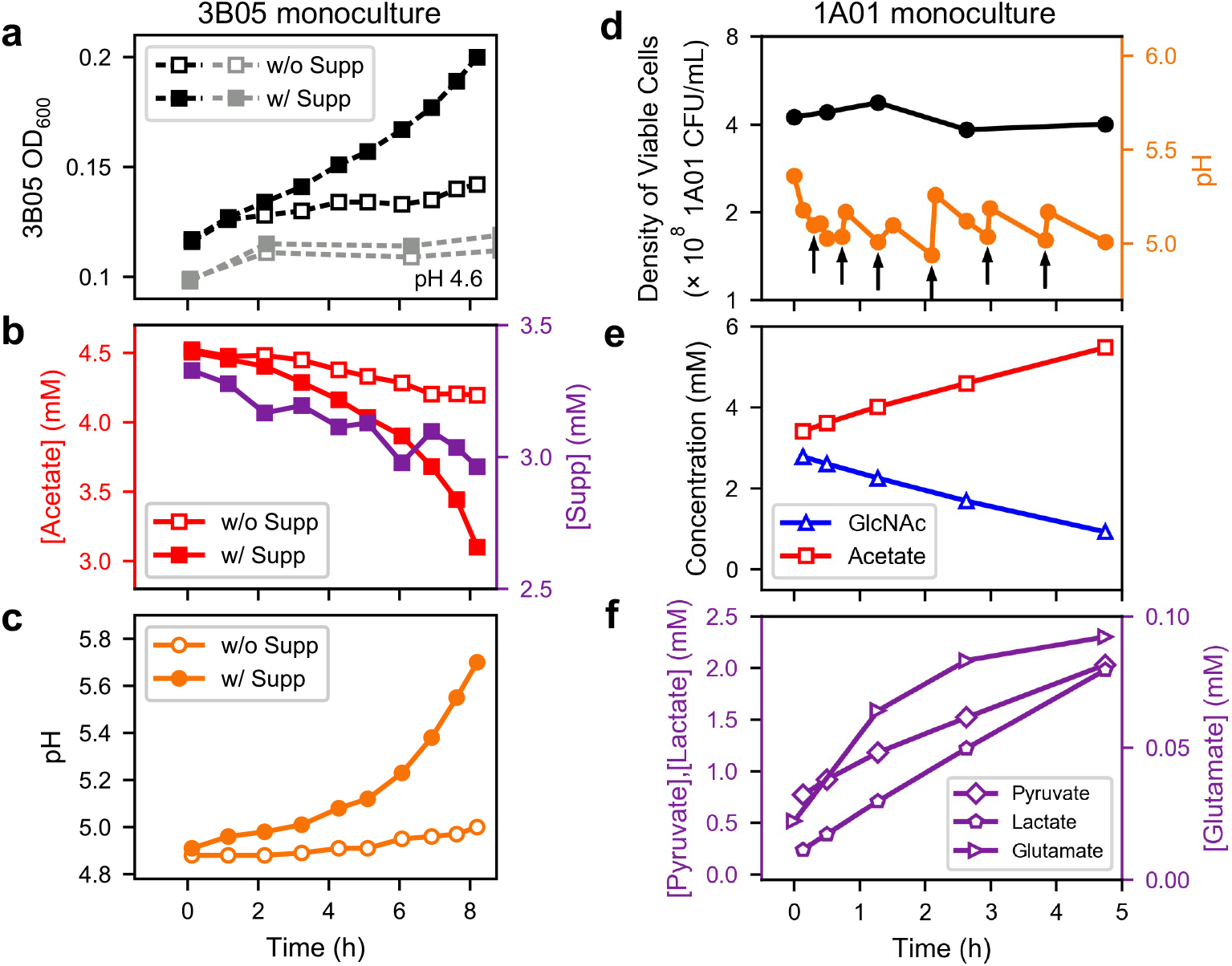
Key physiological and metabolic characteristics of 1A01 and 3B05 monocultures under acetate stress. **(a)-(c)** 3B05 was precultured alone in strongly buffered acetate medium. Then the culture was washed and transferred to a weakly buffered medium (2 mM NaHCO_3_) with 4.5 mM sodium acetate (with pH = 4.9), supplemented with (filled symbols) or without (open symbols) the addition of pyruvate, lactate, and glutamate (referred to collectively as a Supplement, or “Supp”). **(a)** OD, with the grey symbols showing results from the same experiment starting at a lower pH (4.6, by the addition of HCl). The measurement variability for the determination of OD was ±0.002 on the basis of repeated measurements of the same culture sample. **(b)** Acetate (left axis) and the sum of pyruvate, lactate, and glutamate concentrations ([Supp], right axis) in the medium of the two cultures described in panel (a), and **(c)** pH of the medium. The measurement variability for the determination of acetate, pyruvate, lactate, and glutamate concentrations by HPLC was ∼2% on the basis of repeated measurements of the same spent media sample. The pH measurement was accurate to ±0.02 pH unit on the basis of repeated measurements of pH standards. **(d)-(f)** Exponentially growing 1A01 monoculture was initiated at OD = 0.2 and grew in steady-state in GlcNAc medium with the weak buffer (2 mM NaHCO_3_) until acetate excretion dropped the pH to ∼5 where OD reached 0.45, corresponding to viable cell density of ∼4×10^8^ CFU/mL. The pH was then maintained in a narrow pH range by manually titrating with 0.1 M NaHCO_3_. **(d)** Viable cell count (black circles) and pH (orange circles); the arrows indicate times at which NaHCO_3_ was added. **(e)** Concentrations of GlcNAc (blue triangles) and acetate (red squares) in the medium. **(f)** Concentrations of Pyruvate, Lactate, and Glutamate in the medium.

To clarify where these metabolites came from, we maintained an acetate-inhibited 1A01 monoculture in GlcNAc at pH between 5 and 5.5 (orange circles, **Fig. 3d**), to capture the conditions during the acetate peak in the stable cycle of the co-culture where pH dropped below 5.5 (**Fig. 2d**). In this high acetate, low pH condition, 1A01 did not grow (black circles, **Fig. 3d**) but GlcNAc was gradually depleted while acetate, lactate, pyruvate, and glutamate accumulated in the medium (**Fig. 3e, 3f**), in contrast to the accumulation of just acetate under normal pH (**Extended Data Fig. 1c**). The additional metabolites were not mainly from dead/lysed cells, because 1A01 viability did not drop while these metabolites were accumulating (**Fig. 3d**), and more importantly, the amount of carbon released (1.2 mM pyruvate, 1.8 mM lactate, 2mM acetate, totaling ∼13mM of C-atoms in 5 hours) was comparable to that contained in the GlcNAc consumed (≲ 2 mM) during this period (**Fig. 3e,f**). Thus, these metabolites were actively converted from GlcNAc by the growth-arrested 1A01 cells under acetate stress. (The amount of glutamate was negligible compared to pyruvate and lactate and not included here and below.) Untargeted metabolomic analysis^63,64^ of the spent medium of 1A01 monoculture during self-acidification showed the increase of numerous other metabolite features in addition to those already mentioned (**Extended Data Fig. 4a-j**). To see whether the corresponding metabolites may also be cross-fed in the co-culture, we analyzed the spent media collected during the acetate peak using untargeted metabolomics. Many metabolite features were found (purple curves, **Fig. 2f**) with similar dynamics as those exhibited by lactate, pyruvate and glutamate in **Fig. 2e**. Altogether, these data suggest that, in addition to acetate, diverse metabolites were excreted by 1A01 and cross-fed to 3B05, although at a quantitative level, pyruvate, lactate and acetate were the dominant ones.

### Approach to the stable cycle and its independence on the initial inoculation

To understand why the cross-feeding of pyruvate/lactate was able to rescue the co-culture after several growth-dilution cycles but not during the first 24 hours (**Fig. 1d-f**), we developed a mathematical model of acid-induced cross-feeding under growth-dilution dynamics (**Extended Data Fig. 5a, Supplementary Note 3**). Quantitative account of the observed dynamical features by the model required not only the use of strain characteristics obtained in monocultures as described above, but also the lag of 1A01 and the growth of 3B05 at the beginning of the stable cycle (**Fig. 2a**). Additional experiments were performed to show that this lag resulted from a metabolic memory imposed by the acetate stress these cells experienced during the previous cycle; see **Extended Data Fig. 6**. The resulting full model has most parameters fixed by our data, with minimal tuning only for kinetic processes inaccessible experimentally; see **Supplementary Note 3** for a full description. The model was able to capture the stable cycle dynamics quantitatively, including the timing and magnitude of the major metabolites around the acetate peak and the densities of the two species (**Extended Data Fig. 5b**, compare to **Fig. 2a, 2b, 2e**; see **Supplementary Note 4** for details). The model also captured the approach of the co-culture to the stable cycle, quantitatively reproducing the strain abundances and GlcNAc/acetate concentrations at the end of each cycle (**Extended Data Fig. 5c**, compare to **Fig. 1e, 1f**).

Moreover, the model can be used to depict the details of how the co-culture organizes itself dynamically through each growth-dilution cycle to the stable cycle, e.g., for different initial ratio of the two species (**Extended Data Fig. 5d, 5e**, detailed in **Supplementary Note 4**). At 3:1 initial ratio (of 3B05 to 1A01), the stable cycle is predicted to be reached within a single cycle as verified (**Extended Data Fig. 5f**,), with the same stable-cycle characteristics as the case with 1:1 initial ratio (compare **Extended Data Fig. 5g** with **Fig. 2b, 2e**). These results establish that features of the stable cycle are properties of the community, independent of the initial condition and transient dynamics.

### Physiological basis for acid-induced cross-feeding between 1A01 and 3B05

To understand the metabolic basis of the positive interaction between 1A01 and 3B05, we turn to the basic physiological problem faced by bacterial cells under acetate stress ^54,56^. As explained in **Fig. 4**, the presence of a few mM of acetate at low pH (∼5) leads inevitably to the accumulation of high acetate concentration in the cytoplasm with corresponding decrease of endogenous metabolites. This remodeling of the metabolome has several important consequences on bacterial physiology: Based on results from a recent metabolomic study of *E. coli*^56^, we hypothesize that for bacteria growing on glycolytic substrates (such as 1A01 on GlcNAc), respiration becomes limited under acetate stress due to the depletion TCA intermediates, and that cells increase glycolytic flux for energy biogenesis. The lack of free coA (shifted mostly to acetyl coA by mass action due to the high internal acetate concentration) then would force this glycolytic flux be excreted as pyruvate. This scenario, depicted on the left side of **Fig. 4** for 1A01, is supported by the severe depletion of internal glutamate and aspartate, two amino acids reversibly connected to TCA intermediates, under acetate stress (open striped bars, **Extended Data Fig. 4k**). This model rationalizes the continual consumption of GlcNAc, along with a nearly equal-molar excretion of pyruvate and lactate, for growth-arrested 1A01 cells under acetate stress (**Fig. 3d-f, Extended Data Fig. 4b**). Co-excretion of lactate likely resulted from the additional need to release a portion of the NADH generated from glycolysis; see **Fig. 4**.

**Figure 4.**
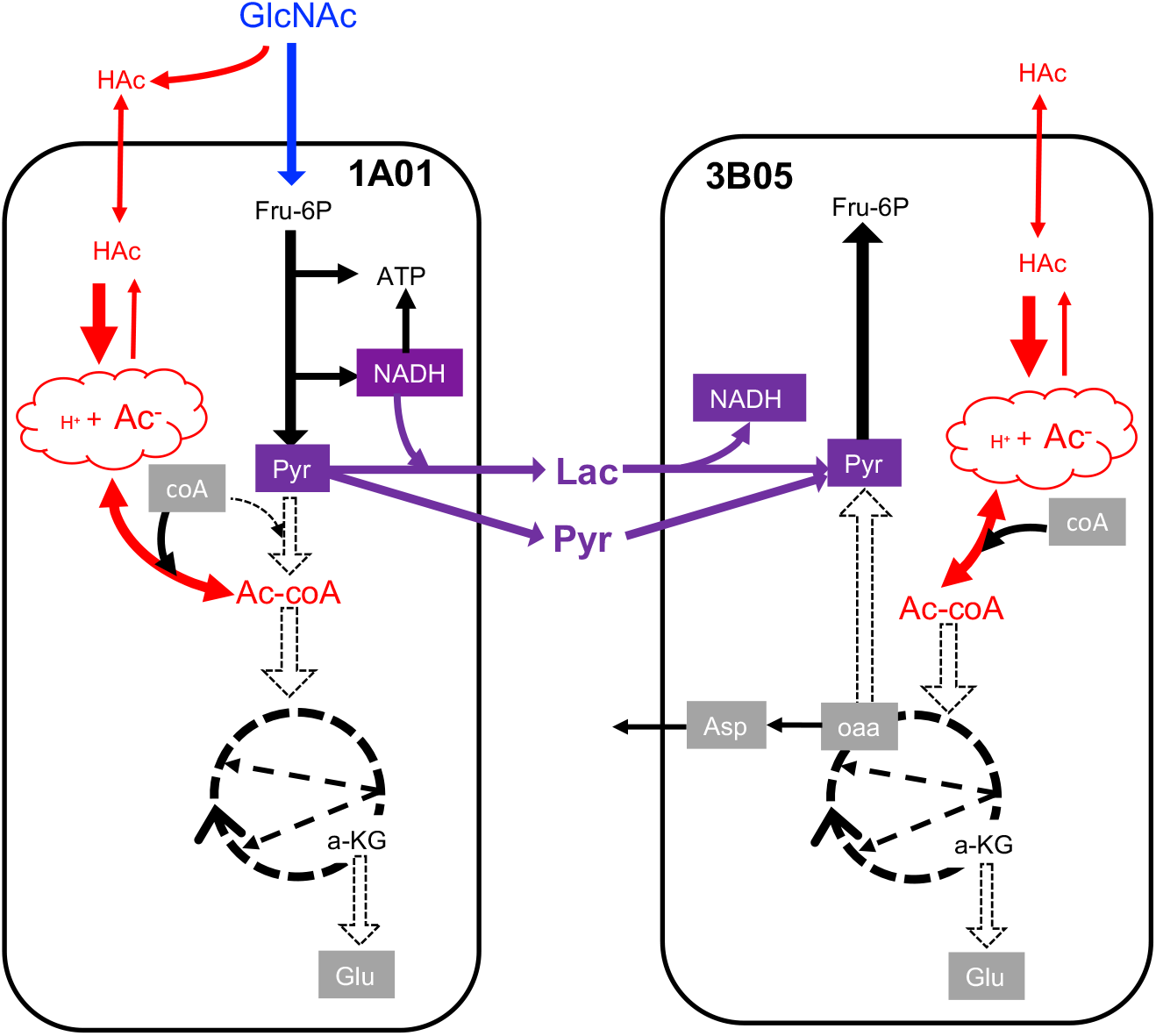
Metabolic model of acid-induced cross-feeding between 1A01 and 3B05. Schematic sketch indicating the key carbon fluxes in 1A01 and 3B05 during acetic acid stress. Metabolites in grey boxes are depleted, metabolites in red are related to acetate, and metabolites in purple (boxes or otherwise) are cross-fed from 1A01 to 3B05. Dashed arrows indicate reactions with negligible flux. The abbreviations are as follows – N-acetyl-glucosamine (GlcNAc), acetic acid (Hac), fructose-6-phosphate (Fru-6P), pyruvate (Pyr), lactate (Lac), acetyl-coA (Ac-coA), α-ketoglutarate (a-KG), oxaloacetate (oaa), aspartate (Asp), and glutamate (Glu). Acetic acid (HAc) is in equilibrium with the anion species, Acetate (Ac^-^), with the ratio of the two concentrations governed by the pH, i.e., 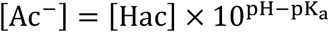 where pK_a_ ≈ 4.75. Because HAc is a small, neutral molecule, it is permeable through the cell membrane. The ratio of the intracellular and extracellular acetate concentrations is given by^54–56^

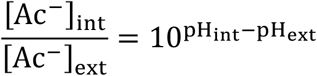

assuming the pK_a_ does not change significantly in the cytoplasm. If the medium pH drops to ∼5, and assuming internal pH is maintained at ∼7, then 3 mM of acetate in the medium would result in ∼300 mM in cells, of the order of the sum of the concentrations of all endogenous metabolites^78^. Based on detailed quantitative studies in *E. coli*, this obligatory flooding of the cytoplasm by acetate has an important physiological consequence: Osmotic balance forces bacteria to adapt by reducing the pools of many endogenous metabolites, particularly TCA intermediates and related amino acids such as glutamate and aspartate, to keep the total metabolite concentration (including acetate) roughly constant^56^. For cells like 1A01 growing on glycolytic substrates, a drop in the coA pool leads to the accumulation of pyruvate and drop in carbon influx, which in turn leads to a drop in the anapleurotic flux and hence reduced growth. The lack of TCA intermediates would further limit the use of the high Ac-coA pool for respiration. Then, glycolytic flux must be increased, with concomitant increase in pyruvate excretion, to supply the cell’s energy demand; see left panel above. Acid-consuming bacteria such as 3B05 would generally be less affected by a low coA pool. Given a high Ac-coA pool, TCA intermediates can be readily generated via the glyoxylate shunt. However, biomass synthesis also requires gluconeogenesis to generate glycolytic intermediates including pyr, pep, and Fru-6P. We hypothesize a bottleneck in the conversion of malate/oaa to pyr/pep during acetate stress, such that 3B05 assimilates acetate into aspartate and glutamate which are excreted back into the medium; see right panel above. Thus, 1A01 and 3B05 form a complementary metabolic partnership under acetate stress: 1A01 cannot move carbon past pyruvate and thus cannot fill the TCA intermediates. (Since 1A01 does not grow on acetate, it is presumably incapable of supplying TCA intermediates from Ac-coA alone.) Because it takes in sugar but does not grow, it has an excess of energy and carbon in the form of lactate and pyruvate. On the flip side, 3B05 has difficulty supplying glycolytic intermediates. Lactate and pyruvate from 1A01 relieves the growth bottleneck of 3B05, allowing it to resume growth and thereby consume acetate, the source of stress. While we have emphasized metabolic interactions in this model, we note that gene regulation would likely also play important roles during the growth recovery process as described in recent studies^46,79^.

The glutamate and aspartate pools of the acetate-utilizing 3B05 were also strongly reduced under acetate stress (filled striped bars in **Extended Data Fig. 4k**). These data suggest reduced pools of TCA intermediates that glutamate and aspartate are reversibly connected to. Since high pools of TCA intermediates are needed to drive gluconeogenesis^65^, we expect 3B05 to have difficulty with gluconeogenesis under acetate stress. This expectation is corroborated by the prolonged excretion of glutamate and aspartate upon exposure to acetate stress (**Extended Data Fig. 4l**, compared to minutes for *E. coli* ^56^), which again indicates bottleneck in gluconeogenesis. This scenario, depicted on the right side of **Fig. 4**, rationalizes the synergy between 1A01 and 3B05 in a co-culture under acetate stress: 1A01 extensively converts GlcNAc into acetate, pyruvate, and lactate even when it is growth-arrested once pH drops and acetate gushes in, while 3B05 uses the pyruvate and lactate excreted by 1A01 to overcome its limitation in gluconeogenesis, allowing it to grow on acetate, eventually removing acetate as the source of stress for both species.

### Unrelated pairs of bacteria show similar metabolic complementarity under acid stress

As the above mechanism of collaborative resistance against acetate stress relies mostly just on the depletion of TCA intermediates upon the accumulation of acetate in the cytoplasm, we expect it to be applicable generally across co-cultures involving bacteria with complementary metabolic types, i.e., glycolytically oriented sugar consumers and gluconeogenically oriented acid consumers. To test the predicted generality, we selected *E. coli* along with three bacterial species from a previously studied soil bacterial consortium^29,66,67^ to test the generality of acid-induced cross-feeding (**Fig. 4**), established so far between 1A01 and 3B05. We performed growth-dilution experiments, pairing a species from the Enterobacteriaceae family, which prefers growing on sugars while excreting acetate, with a species from the Pseudomonadaceae family, which prefers growing on organic acids including acetate^66,68^. When *Citrobacter freundii* from Enterobacteriaceae was grown in GlcNAc alone with a weak phosphate-based buffer, the monoculture stopped growing at a low OD when pH dropped below 6 (vertical dashed line, **Extended Data Fig. 7a**). Analyzing the spent media in the *C. freundii* monoculture, we found the accumulation of acetate and pyruvate, with the amount of acetate increasing above ∼1.5 mM as pH dropped below 6 (**Extended Data Fig. 7b,c**). Adding *Pseudomonas fluorescens* to the culture extended the saturating OD by 3∼4 fold (solid black line, **Fig. 5a**), suggesting the possibility of acid-induced cross-feeding in the co-culture. To test the occurrence of the latter, we first confirmed that the co-culture maintained coexistence over several 24-h growth-dilution cycles (**Fig. 5b**). Using a pH-sensitive dye to continuously monitor the pH dynamics in co-cultures (**Extended Data Fig. 7d, e**), we found that the pH dropped below 6 for several hours in the middle of each growth-dilution cycle, before recovering to the starting pH (**Fig. 5c**). Analysis of the spent medium revealed the accumulation of pyruvate in addition to acetate, peaking during the trough of the pH dip (**Fig. 5d**). These observations are highly analogous to the dynamics exhibited by the 1A01-3B05 co-culture (**Fig. 2**), despite the very different characteristics of these species pairs.

**Figure 5.**
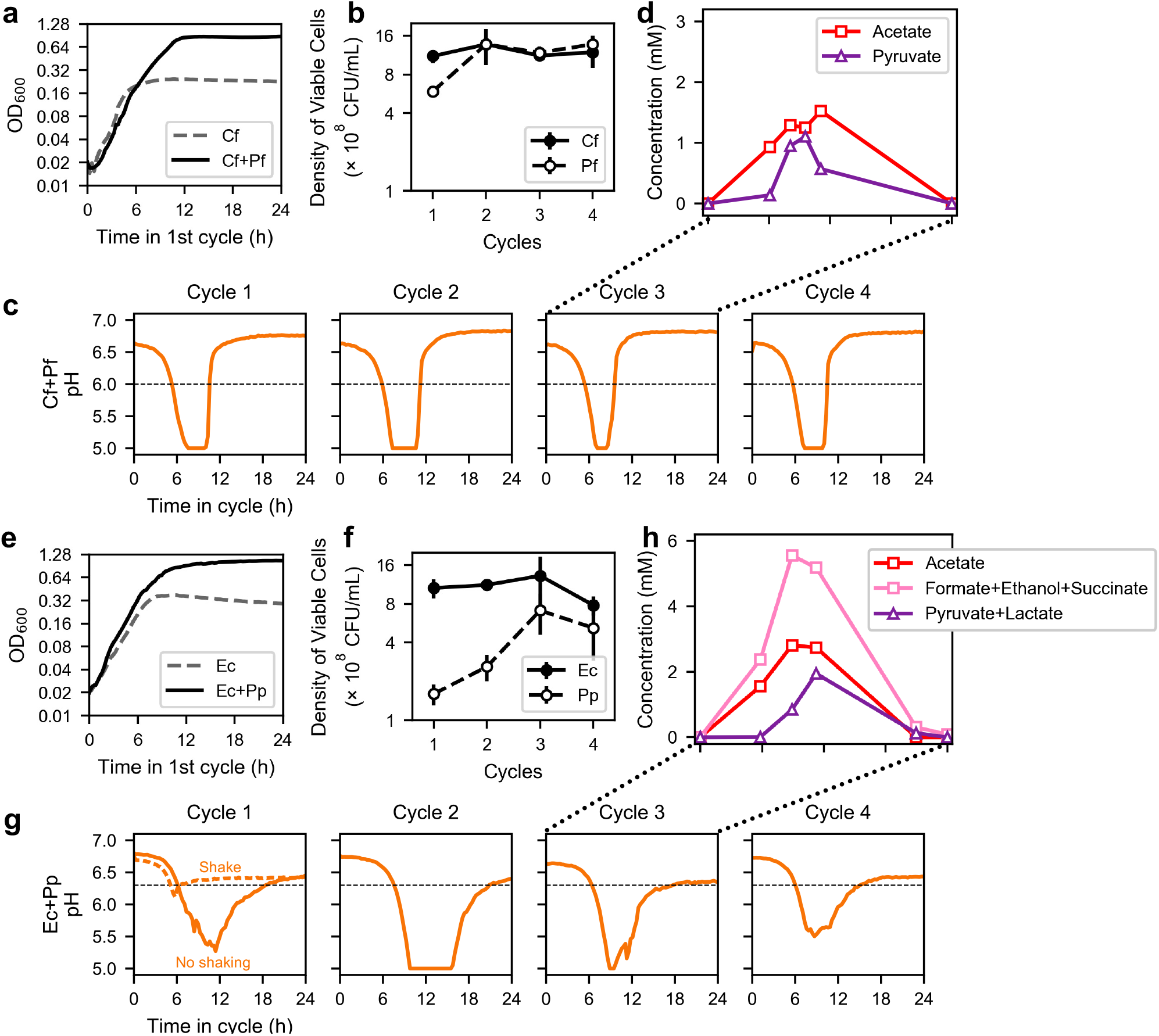
Acid-induced cross-feeding between ‘acid excreters’ and ‘acid eaters’. **(a)** Comparison of growth curves of *Citrobacter freundii* (Cf) alone with shaking in 10 mM GlcNAc as the sole carbon source in weak phosphate-based buffer (dashed grey line), and with *C. freundii* growing in the same medium and condition in co-culture with *Pseudomonas fluorescens* (Cf+Pf, solid black line). **(b)-(d)** Results from co-cultures of *C. freundii* and *P. fluorescens* grown with shaking in 10 mM GlcNAc as the sole carbon source and low buffer, in 24-h growth-dilution cycles with 100x dilution. The co-culture was started from a 1:1 mixture of exponentially growing cells of each species, each at OD=0.01. **(b)** The viable counts of *C. freundii* (Cf) and *P. fluorescens* (Pf) cells at the end of each cycle. The error bars indicated the standard error from 6 biological replicates. **(c)** pH dynamics throughout the first four cycles measured as described in **Extended Data Fig. 7d,e**. The horizontal dashed black line indicates the pH below which acid-induced excretion was observed under identical growth conditions for a monoculture of *C. freundii* (**Extended Data Fig. 7a-c**). **(d)** Metabolites measured in the medium during Cycle 3. Purple triangles indicate the concentration of pyruvate. **(e**) Comparison of the growth curve of *E. coli* (Ec) growing alone with 10 mM glucose as the sole carbon source without shaking in medium strength buffer (dashed grey line), and with *E. coli* growing in the same medium and condition in coculture with *Pseudomonas putida* (Ec+Pp, black line). **(f)-(h)** Results from co-cultures of *E. coli* and *P. putida* described in panel (e), grown in 24h growth-dilution cycles with 100x dilution. The co-culture was started from a 1:1 mixture of exponentially growing cells each at OD=0.01. **(f)** Viable counts of *E. coli* (Ec) and *P. putida* (Pp) cells at the end of each cycle. The error bars indicated the standard error from 6 biological replicates. **(g)** pH of the co-culture throughout the first four cycles are shown as the solid orange line. The dashed orange line in Cycle 1 indicates the pH of the coculture when shaken. The horizontal dashed black line indicates the pH below which acid-induced excretion was observed under identical growth conditions for a monoculture of *E. coli* (**Extended Data Fig. 7f-h**). **(h)** Metabolites measured in the spent medium of Cycle 3. The sum of the concentrations of succinate, formate, and ethanol, major fermentation products excreted by *E. coli* during anaerobic growth, is shown in pink. Sum of the concentrations of pyruvate and lactate, whose excretions were induced by acetate (**Extended Data Fig. 7f-h**), is shown as purple triangles.

GlcNAc is a unique sugar with an extra acetyl group that leads to high acetate excretion under aerobic growth (**Extended Data Fig. 1e**). To see whether acid-induced cross-feeding between pairs of growing species established here may be applicable to other means of acetate accumulation, we also examined the effect of self-acidification through poor aeration by culturing with exposure to air but not shaking, as was done in a number of recent studies^29,66,67^; see **Methods**. Here, we chose *E. coli* as the sugar-consuming acid excreter and *Pseudomonas putida* as the acid eater. Growing *E. coli* alone in glucose minimal medium with a weak phosphate-based buffer, we again found the monoculture to stop growing as pH dropped (**Extended Data Fig. 7f**). The medium accumulated acetate, succinate, formate, and ethanol (**Extended Data Fig. 7g**), the canonical fermentation products excreted by *E. coli* during anaerobic growth soon after growth started, with lactate and pyruvate accumulating as growth slowed down (vertical dashed line, **Extended Data Fig. 7h**).

Addition of *P. putida* again substantially extended the growth of the co-culture, suggesting cross-feeding (**Fig. 5e**). The two species coexisted in 24-h growth-dilution cycles (**Fig. 5f**), and pH dynamics again revealed the repeated dip for several hours in the middle of each cycle (**Fig. 5g**). Measurement of the co-culture media during the third cycle (**Fig. 5h**) showed the buildup and depletion of acetate (red squares), the remaining anaerobic excretants (pink squares), and the acid-induced excretion (purple triangles). To confirm the role of oxygen deprivation in the observed phenomena, we repeated the experiment with the co-culture shaken throughout the cycle: the sharp dip in pH disappeared in this case (dashed orange line, **Fig. 5g**, Cycle 1).

## Discussion

Weak acids are excreted by fast-growing sugar-eating bacteria in many environments^31–39^. While these weak acids serve as natural growth substrate for a variety of acid-eating bacteria, typically the acid eaters grow more slowly than the sugar eaters and this mismatch of growth rates inevitably leads to a rapid buildup of the excreted acids as we found in a co-culture of the marine bacteria 1A01 and 3B05. This acid buildup would crash the pH once the buffer capacity of the medium is exceeded, putting the community of bacteria under acid stress. Because the acid eaters are not capable of growing on acid at low pH (**Fig. 1d, 3a**), a puzzle is presented on how the co-culture is able to remove the acid and restore growth under repeated growth-dilution cycles (**Fig. 1e,f**). Using detailed, quantitative analysis, we revealed a hidden layer of metabolic collaboration that occurs in a co-culture of 1A01 and 3B05 during acid stress. As depicted in the metabolic model of **Fig. 4**, a positive interaction is realized whereby acid-induced excretion of pyruvate and lactate by 1A01 helped 3B05 to grow on the excreted acid, hence detoxifying the environment for both species. Despite the complexity of the metabolic interactions, a simple dynamical model with constrained parameters was sufficient to capture the bulk of the observed dynamics (**Extended Data Fig. 5**).

Rather than being an interaction specific to marine isolates, acid-induced cross-feeding appears to be a collaborative mechanism that occurs generally between complementary cell types – glycolytically oriented, sugar-consuming acid excreters and gluconeogenically oriented acid eaters (**Fig. 6a**). Recent work suggests that intrinsic limitations on the directionality of carbon metabolism force species to pick whether to excel at glycolytic or gluconeogenic metabolism^68^. Thus acid-induced cross-feeding is a positive interaction that arises not specifically for this purpose; rather it occurs as a by-product of the natural division of copiotrophic, heterotrophic bacteria into glycolytically and gluconeogenically oriented modes of metabolism. Here we showed acid-induced cross-feeding between *Pseudomonas* species from soil isolates and Enterobacteriaceae such as *C. freundii* and *E. coli* in addition to *Vibrio* sp. 1A01 and *Neptumonas* sp. 3B05. Recent work suggests that *Staphlococcus aureus* and *Pseudomonas aeruginosa* likely have the same metabolic complementarity, and acid-induced cross-feeding may also promote the coexistence of these species during infection^69^.

**Figure 6.**
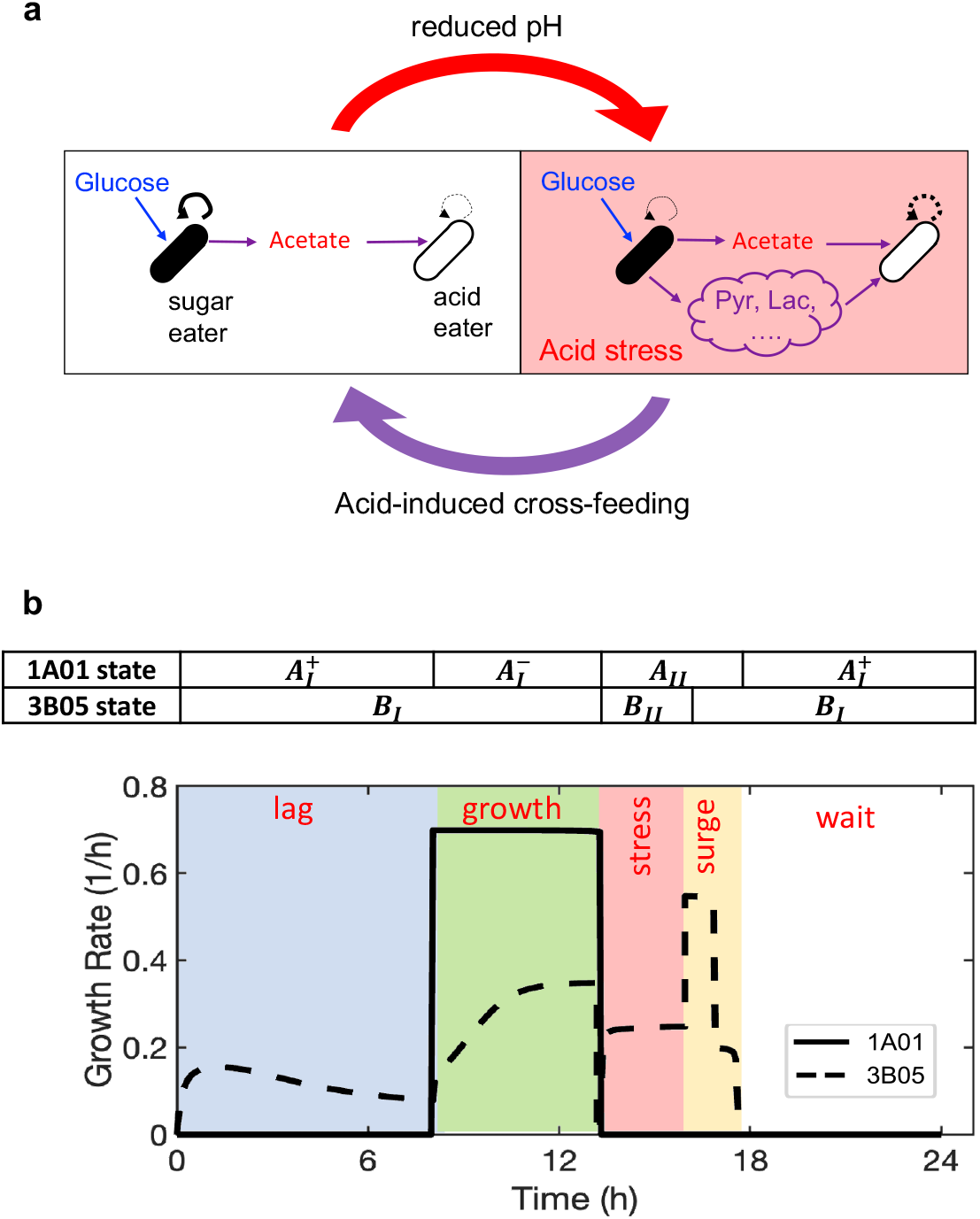
Different phases of community dynamics. **(a)** Cartoon of acid-induced cross-feeding. During growth, weak acids are excreted by sugar eaters for a variety of reasons. As long as the excreted flux exceeds that of the consumption by ‘acid eaters’ (left panel), the concentration of the excreted acid will accumulate in the medium, eventually reducing the medium pH when the accumulate acid exceeds the buffer capacity of the medium. The reduced pH results in an acid stress that inhibits the growth of *both* species. During the stress, acid-induced cross-feeding enables the acid eater to remove the accumulated acid for both types of species (right panel). **(b)** Instantaneous growth rate of 1A01 (solid line) and 3B05 (dashed line) through a stable cycle according to the model (**Extended Data Fig. 5a, Supplementary Note 3**). The abrupt changes of growth rates are defined by a number of phases of the co-culture, indicated by the colored bands. The latter arose due to a combination of the physiological states each species goes through during the cycle, as indicated by the table above the plot and defined in **Extended Data Fig. 5a**.

Our data provide a physiological basis for the general idea of microbial diversity promoted by extensive cross-feeding among free-living bacteria. For each pair of co-culture we investigated, mM level of valuable metabolites were excreted in addition to acetate, allowing substantial growth by acid eaters on the excreted substrates. Sustained excretion at this level was not a minor leakage by stressed cells. In the case of 1A01, excretion was sustained by non-growing cells which actively took up GlcNAc from the medium and converted them into pyruvate and lactate (**Fig. 3d-f**). We suggest two benefits for such extensive excretion by 1A01: An immediate benefit is that, due to limited respiration in acetate stressed cells^56^, glycolysis is an effective way for 1A01 to generate energy for its maintenance even when it is inhibited from growing (**Fig. 4**). Another is that 1A01 would die rapidly over the course of a day (**Extended Data Fig. 3f, 3g**) if it is left under acetate stress without rescue by 3B05, and the latter occurs only in the presence of large amounts of pyruvate and lactate sustained by 1A01. In all of the cases studied here, stress was a pre-requisite before excretion other than acetate took place. This picture – in which stressed cells extensively excreted metabolites while not growing – challenges the current theoretical picture in which cells are continuously sharing metabolites during steady-state exponential growth (as attained in a chemostat), even in growth-dilution scenarios^30,70^. In support of this multi-stage cross-feeding picture, coexistence of 1A01 and 3B05 which occurred in 24-h growth-dilution cycles featuring extended growth arrest failed to establish in the same system under 6-h cycles that provided steady, stress-free growth (**Fig. 1g**).

Our work suggests that niches for different species are created *out of* steady-state growth as gradients in stresses emerge^71–73^. Using our model which quantitatively captured co-culture dynamics (**Extended Data Fig. 5**), we plotted in **Fig. 6b** the growth rate of 1A01 and 3B05 throughout the duration of the stable cycle: The plot shows different species dominating in different phases of the cycle, a key ingredient for the maintenance of both species in the system, reminiscent of early theoretical studies of simple growth-dilution cycles^74,75^. Here, different phases of the community (indicated by bands of different colors in **Fig. 6b**) resulted from interactions of different physiological states of each species, a theme that should be even more relevant to more complex communities. Thus, species abundances and nutrient levels observed at the end of growth-dilution cycles in microbial ecology studies likely depend on the dynamics of the community throughout the cycle as established in simplified systems here, with metabolic and possibly other modes of interactions giving rise to species dominance within different time windows of a cycle, in stark contrast to steady state models that have guided microbial ecology research for many decades^76,77^.

## Acknowledgments

We are grateful to helpful discussions with numerous colleagues during the course of this work, including Josh Goldford, Jeff Gore, Seppe Kuehn, Pankaj Mehta, Arvind Murugan, Shaul Pollak, Wenying Shou, and members of the Hwa lab, particularly Hiro Okano and Jack Reddan for technical assistance. We would like to thank Josh Goldford, Alvaro Sanchez, and Sylvie Estrela for sharing growth protocols. This work is supported by the Simons Foundation through the Principles of Microbial Ecosystems (PriME) collaboration (Grant no. 542387 to TH, 542395 to OXC, 608247 to SP) and the NIH (Grant 5R01GM095903 to TH). KA acknowledges a Simons PriME fellowship (Grant no. 647858). JD acknowledges the hospitality of UCSD during her sabbatical visit, and the support of the NSF through grants DMR-1703231 and MCB-2029480.

## Data and material availability

Raw mass spectral data is deposited to massIVE and accessible with the accession code MSV000087136 (ftp://MSV000087136@massive.ucsd.edu, password Reviewer123), and will be made publicly accessible upon publication.

## Extended Data Table and Figures

**Extended Data Table 1.**
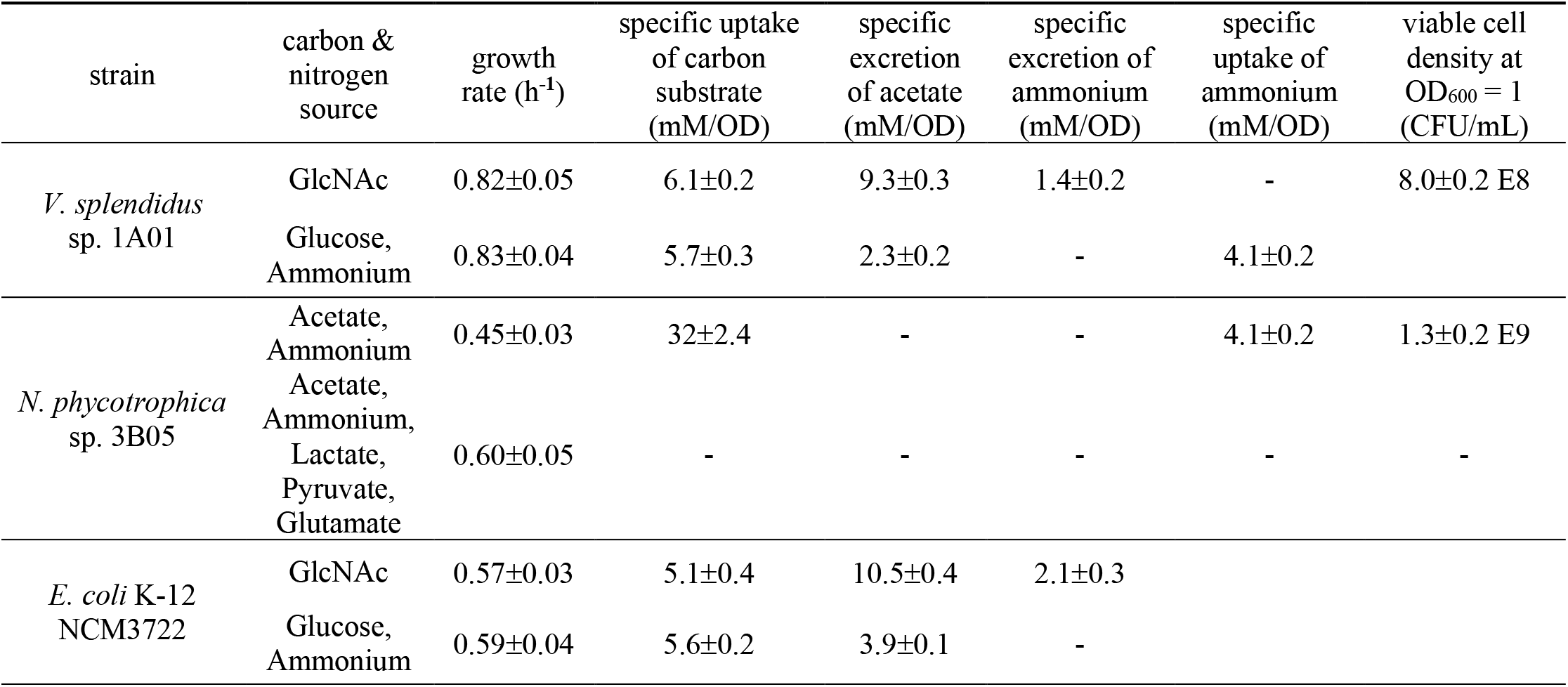
Monoculture steady-state physiological parameters for *V. splendidus* sp. 1A01, *N. phycotrophica* sp. 3B05, and *E. coli* NCM3722. Note that the growth yield on a substrate is the inverse of the specific uptake of that substrate listed in the table, with biomass density given in unit of OD_600_; the conversion of the latter to cell density is given by the last column. 1A01 and 3B05 were grown at 27°C in 0.35 M NaCl in the strongly buffered HEPES medium described in Materials and Methods; all parameters were the same in the weakly buffered medium with 2mM sodium bicarbonate, except for 1A01’s GlcNAc growth rate, which was 0.72/h, and 3B05’s growth rate on acetate, which was 0.35/h. *E. coli* NCM3722 was grown at 37°C in MOPS medium with 0.4 M NaCl; the yields in 0.4 M NaCl were indistinguishable from those in 0.1 M NaCl suggesting that the high excretion of acetate of 1A01 during growth on GlcNAc is not due to its adaptation to growing in higher salinity environments. Standard error estimated from at least three biological replicates.

| strain | carbon & nitrogen source | growth rate (h<sup>-1</sup>) | specific uptake of carbon substrate (mM/OD) | specific excretion of acetate (mM/OD) | specific excretion of ammonium (mM/OD) | specific uptake of ammonium (mM/OD) | viable cell density at OD<sub>600</sub> = 1 (CFU/mL) |
| --- | --- | --- | --- | --- | --- | --- | --- |
| *V. splendidus* sp. 1A01 | GlcNAc | 0.82±0.05 | 6.1±0.2 | 9.3±0.3 | 1.4±0.2 | - | 8.0±0.2 E8 |
|  | Glucose, Ammonium | 0.83±0.04 | 5.7±0.3 | 2.3±0.2 | - | 4.1±0.2 |  |
| *N. phycotrophica* sp. 3B05 | Acetate, Ammonium | 0.45±0.03 | 32±2.4 | - | - | 4.1±0.2 | 1.3±0.2 E9 |
|  | Acetate, Ammonium, Lactate, Pyruvate, Glutamate | 0.60±0.05 | - | - | - | - | - |
| *E. coli* K-12 NCM3722 | GlcNAc | 0.57±0.03 | 5.1±0.4 | 10.5±0.4 | 2.1±0.3 |  |  |
|  | Glucose, Ammonium | 0.59±0.04 | 5.6±0.2 | 3.9±0.1 | - |  |  |

**Extended Data Figure 1.**
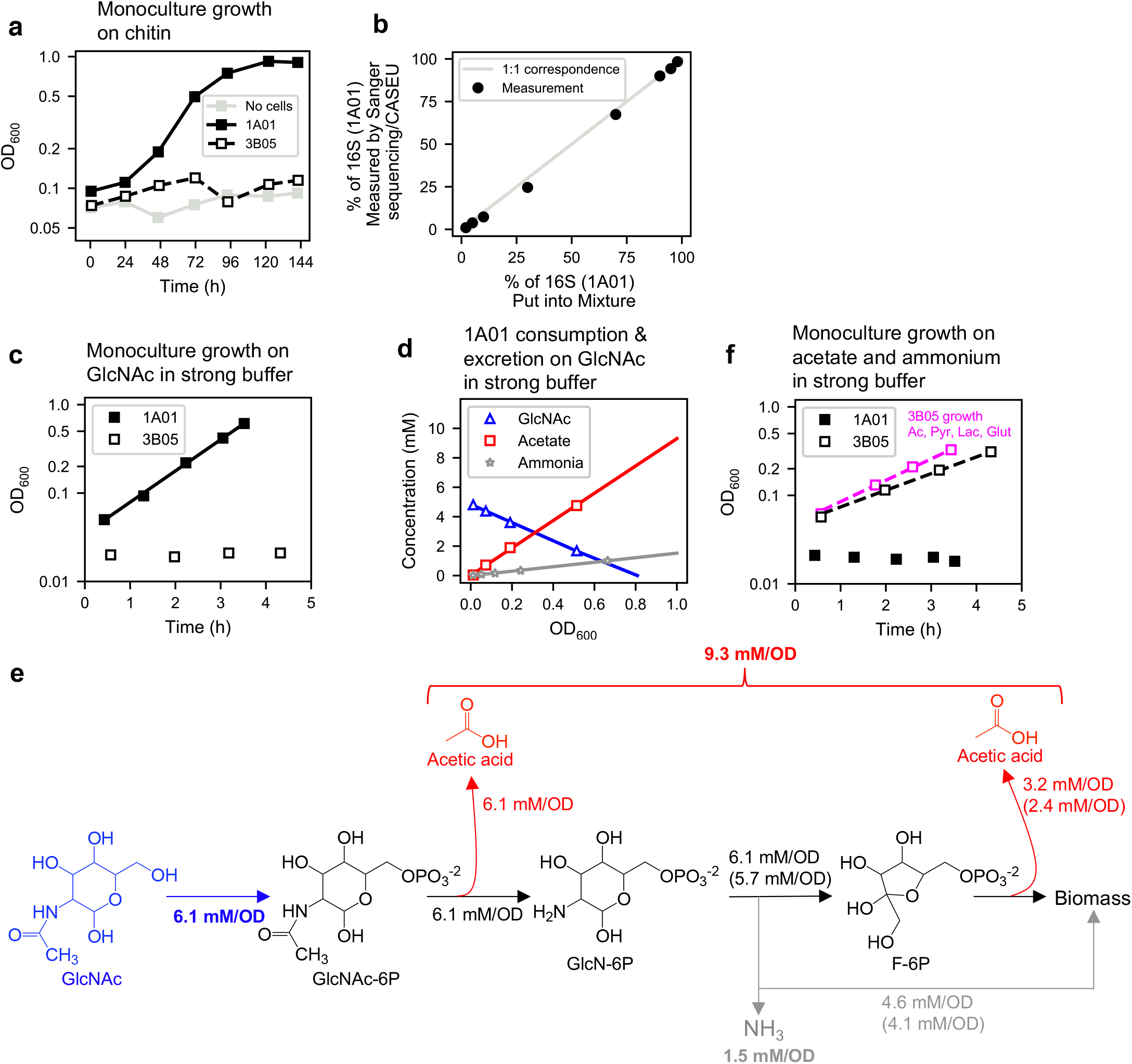
Single-strain characteristics of 1A01 and 3B05 growing in strong buffer. All results for batch culture growth described in this work were done in medium with 40 mM HEPES buffer and 0.35 M NaCl. The culture was maintained in water bath shaker at 27°C and 250 rpm. See details of growth medium and condition, including pre-culturing, in the Methods. **(a)** Growth on chitin. 1A01 and 3B05 were grown in monoculture with 0.2% w/v chitin chips as the sole carbon and nitrogen source. 1A01 was pre-cultured in GlcNAc HEPES minimal medium with ammonium and 3B05 was pre-cultured in sodium acetate HEPES with ammonium. OD_600_ measurements were taken from well-mixed culture after allowing the visible chitin chips to sink for 4 min. To see the effect of residual small chitin pieces in the culture, a ‘no cells’ control is also shown as light grey squares. **(b)** Accuracy of Sanger sequencing electropherograms in determining the composition of 1A01 and 3B05 16S amplicon. Purified 16S amplicon of 1A01 and 3B05 were mixed in defined proportions using their concentrations measured by Nanodrop (expected concentration, light grey line). The mixtures were submitted for Sanger sequencing and the resulting electropherograms were deconvolved using CASEU^49^ to determine the fraction of each type of 16S sequence. **(c)** Steady-state growth of 1A01 (filled black squares) and 3B05 (open black squares) on in minimal medium with 5 mM GlcNAc as the sole carbon and nitrogen source buffered by 40 mM HEPES. See Methods for details of the growth protocol including preculturing preceding time ‘0’. The solid line indicates an exponential fit to the 1A01 growth curve (0.81/h with R^2^=0.999). **(d)** Depletion of GlcNAc (blue triangles) and accumulation of acetate (red squares) and ammonium (grey stars) during steady-state growth of 1A01 on GlcNAc. The solid lines indicate linear fits, with the inverse biomass yield on GlcNAc being 6.1 mM/OD (R^2^ = 0.999), and acetate and ammonium excretion yields of 9.3 mM/OD (R^2^ = 0.998) and 1.5 mM/OD (R^2^ = 0.999), respectively. The variance computed from 3 biological replicates is indicated in **Extended Data Table 1. (e)** GlcNAc degradation pathway delineated previously for a *Vibrio* species^50^ and for *E. coli*^80^ depicting the excretion of acetic acid (red) and ammonium (grey). The yield numbers in bold are based on the measurements during exponential growth (panel c). The numbers in regular text are deduced based on stoichiometry and the measured values. The numbers in parenthesis are numbers obtained from similar measurements during exponential growth of 1A01 on glucose and ammonium (where 5.7 mM of glucose was consumed for 1 OD of cells). As shown in panel (d), we measured 9.3 mM/OD of acetate excreted during growth on GlcNAc. Based on stoichiometry, we expect 6.1 mM/OD of acetate from the conversion of GlcNAc-6-P to GlcN-6-P (left red arrow) and 3.2 mM/OD from acetate overflow^42^ (right red arrow). The latter is in line with what we measured for acetate excretion on glucose (2.4 mM/OD). Similarly based on stoichiometry, we expect 6.1 mM/OD of ammonium to be released from GlcN-P. We measured 1.5 mM/OD of ammonium excreted, implying 4.6 mM/OD of ammonium was assimilated into biomass, in line with the degree of ammonium assimilation we measured during growth on glucose (4.1 mM/OD). This analysis shows that growth yield on GlcNAc is similar to the yield on glucose and ammonium, so that the additional excretion of acetate and ammonium for the growth on GlcNAc compared to growth on glucose can be understood approximately as a result of the “extra” acetyl and amine groups accompanying the glucose in GlcNAc. In support of this interpretation, we found similar yield of acetate and ammonium due to excretion by *E. coli* growing on GlcNAc MOPS minimal medium with 0.4 mM NaCl (**Extended Data Table 1**). The excretion yield for glucose is also provided there for reference. **(f)** Steady-state growth of 1A01 (filled black squares) and 3B05 (open black squares) on HEPES-buffered minimal medium with 60 mM sodium acetate as the sole carbon source and 10 mM of ammonium chloride as the sole nitrogen source. The dashed black line indicates an exponential fit to the 3B05 growth curve, with growth rate 0.43/h (R^2^=0.999). The open black triangles indicate the growth curve of 3B05 on 20 mM acetate supplemented with 20 mM each of lactate, pyruvate, and glutamate. The dashed black line through the triangles indicates an exponential fit to the growth curve, with growth rate 0.58/h (R^2^ = 0.998). The best-fit parameters in panels c, d, and f are summarized in **Extended Data Table 1**.

**Extended Data Figure 2.**
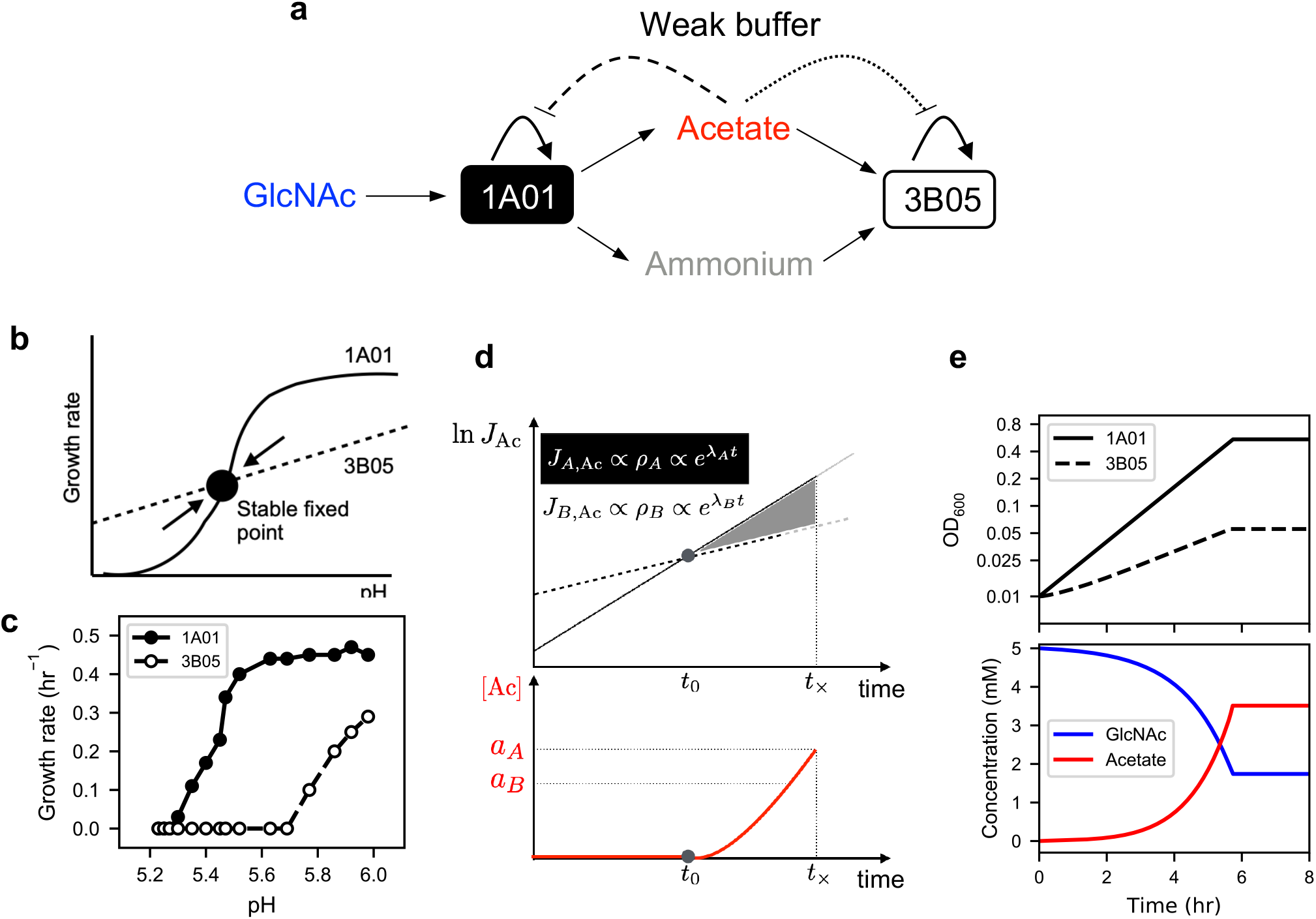
Acetate cross-feeding cannot support exponential growth of 1A01 and 3B05 co-culture in weak buffer. **(a)** Solid arrows indicate schematic of acetate cross-feeding based on single-strain characteristics derived from **Extended Data Fig. 1**. In the weak buffer, acetate excretion is expected to lead additional to a drop in pH and inhibition of cell growth, as indicated by the dashed and dotted lines. A canonical scenario of syntrophy is realized if the effect of growth inhibition on the acid excreter (1A01) is stronger than that on the acid eater (3B05); see panel b. However, if the growth of 3B05 is inhibited more strongly (panel c), then the acetate build-up is inevitable (panel d), leading to the growth arrest of both species before the consumption of the provided nutrients (panel e). **(b)** In the canonical scenario of syntrophy, growth of the strain generating toxicity should be more sensitive to toxicity (solid black line) than the strain relieving toxicity (dashed black line) as illustrated. The intersection of these two lines is the stable fixed point describing a stable, exponential growing coculture: above this pH, the faster-growing strain (1A01) would excrete more acetate than what the de-toxifier (3B05) could take up, resulting in a decrease in the pH. Below this pH, 3B05 would be able to take up acetate faster than 1A01 could produce it, resulting in an increase in the pH. This balance would lead to a stable exponential growth phase for the coculture. **(c)** Measured dependences of the growth rate of 1A01 (solid circles) and 3B05 (open circles) on the medium pH. Cells were grown in minimal medium buffered by 10 mM MES with different ratios of the acid and base form to obtain the desired pH; otherwise, the medium was identical to the HEPES-buffered medium. Glycerol was used as the sole carbon source as both strains grew on it and neither strain excreted acetate or other fermentation products which would have changed the medium pH during the course of experiment, making the results difficult to interpret. The data shows that 3B05 is more sensitive to pH than 1A01, hence precluding the scenario of a stable, exponentially-growing co-culture depicted in panel b. **(d)** Qualitative account of the co-culture dynamics of 1A01 and 3B05 in weak buffer. Because the acetate excretion (*J*_*A,*A*C*_) and uptake flux (*J*_*B,*A*C*_) is proportional to the density of 1A01 and 3B05, respectively (Eq. (1.4), **Supplementary Note 1**), each flux increases exponentially with the corresponding growth rate (*λ*_*A*_ and *λ*_*B*_), shown by the solid and dashed black lines. Since *λ*_*A*_ > *λ*_*B*_ at all pHs (panel c), acetate excretion by 1A01 (solid black line) will eventually exceed its consumption by 3B05 (dashed black line); this is marked by the circle at time *t*_0_. For *t* > *t*_0_, the acetate concentration would rise approximately exponentially in time (red line, bottom panel), until 3B05 stops growing when a “stopping concentration” α_*B*_ (corresponding to the pH where the growth of 3B05 stops, panel c) is reached. Soon after that (at time *t*_×_), acetate concentration reaches the value α_*A*_, corresponding to the pH where 1A01 ceases growing also. At this point, the entire co-culture ceases. **(e)** Results of calculation based on a quantitative model (detailed in **Supplementary Note 1**) with single-strain characteristics derived from **Extended Data Fig. 1** and acid response data shown in panel (b). The model outputs on the densities of 1A01 and 3B05 cells (solid and dashed lines) agree quantitatively with the observed viable cell densities shown in **Fig. 1d**, and the model output on the concentrations of GlcNAc (blue line) and acetate (red line) also agree quantitatively with those measured (**Extended Data Fig. 3b, 3f**) up to the time of growth arrest. In particular, the model correctly predicted growth arrest to occur about 6 hours after inoculation, with about half of the initial GlcNAc still remaining at that point. This simple model does not predict what occurs after the growth arrest, which is main topic of the current study.

**Extended Data Figure 3.**
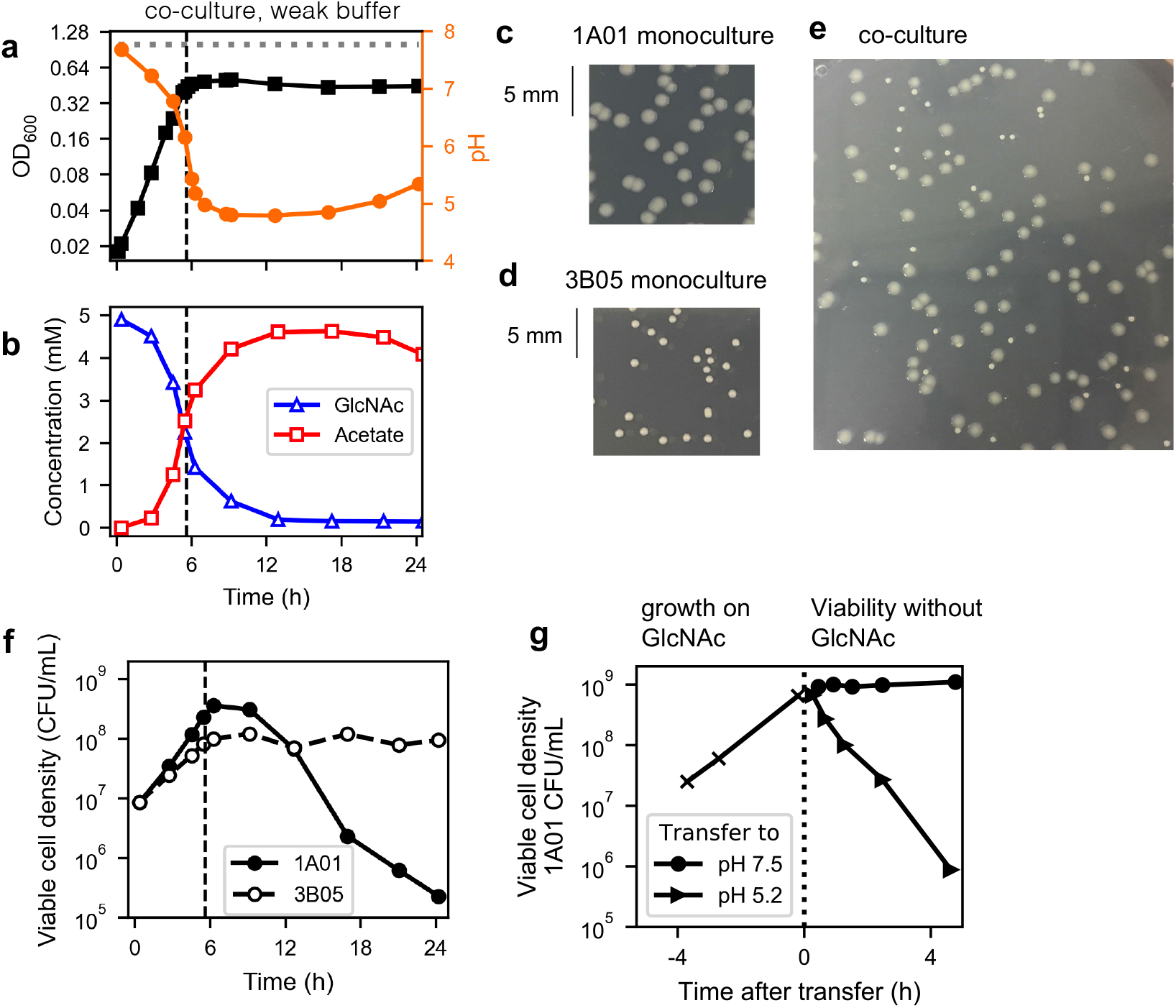
1A01-3B05 co-culture grown in the weak buffer. Measurement during a 24-hr growth period for the co-culture in 5 mM GlcNac with the weak 2 mM bicarbonate buffer. **(a)** shows the OD (black squares) and pH (orange circles), with the horizontal dotted line indicating the final OD reached by the same culture grown in strong buffer. **(b)** shows the GlcNAc (blue triangles) and acetate (red squares) concentrations in the medium. The vertical dashed line indicates the time where growth in OD ceased. **(c-e)** Colonies of 1A01 and 3B05 exhibit different morphologies. 1A01 monoculture **(c)**, 3B05 monoculture **(d)**, and 1A01/3B05 **(e)** coculture growing in HEPES buffered medium was plated on 1.5% agar with rich medium (marine broth). After incubation of the plate for ∼36 hr at 27°C, 1A01 and 3B05 colonies were visibly different. 1A01 colonies have a much larger area and an off-white color, while 3B05 colonies are punctate and white. **(f)** Viable cell counts for 1A01 (filled black circles) and 3B05 (open black circles) measured during the 24-h growth period for the co-culture grown in weak buffer. **(g)** The death of 1A01 in monoculture was characterized by measuring the viable cell density (colony forming units or CFU per mL of culture) before and after exponentially growing 1A01 monoculture is transferred to medium with different pH: Before the transfer, 1A01 grew in HEPES-buffered medium with GlcNAc as the sole carbon and nitrogen source. Viable cell count (black x’s) showed exponential growth consistent with growth rate obtained from OD measurement (**Extended Data Fig. 1c**). At time t=0 (black dotted line), these cells were washed and resuspended into GlcNAc-free medium with 2mM bicarbonate buffer, with the pH set to 7.5 (black circles) or pH 5.2 (black triangles). In the latter case, 3.6 mM of acetic acid was added to lower the pH. Rapid drop of viable 1A01 cells was observed for the ones transferred to the medium at low pH but not at normal pH. Thus, low pH with acetate and lack of GlcNAc were sufficient for the rapid death of 1A01.

**Extended Data Figure 4.**
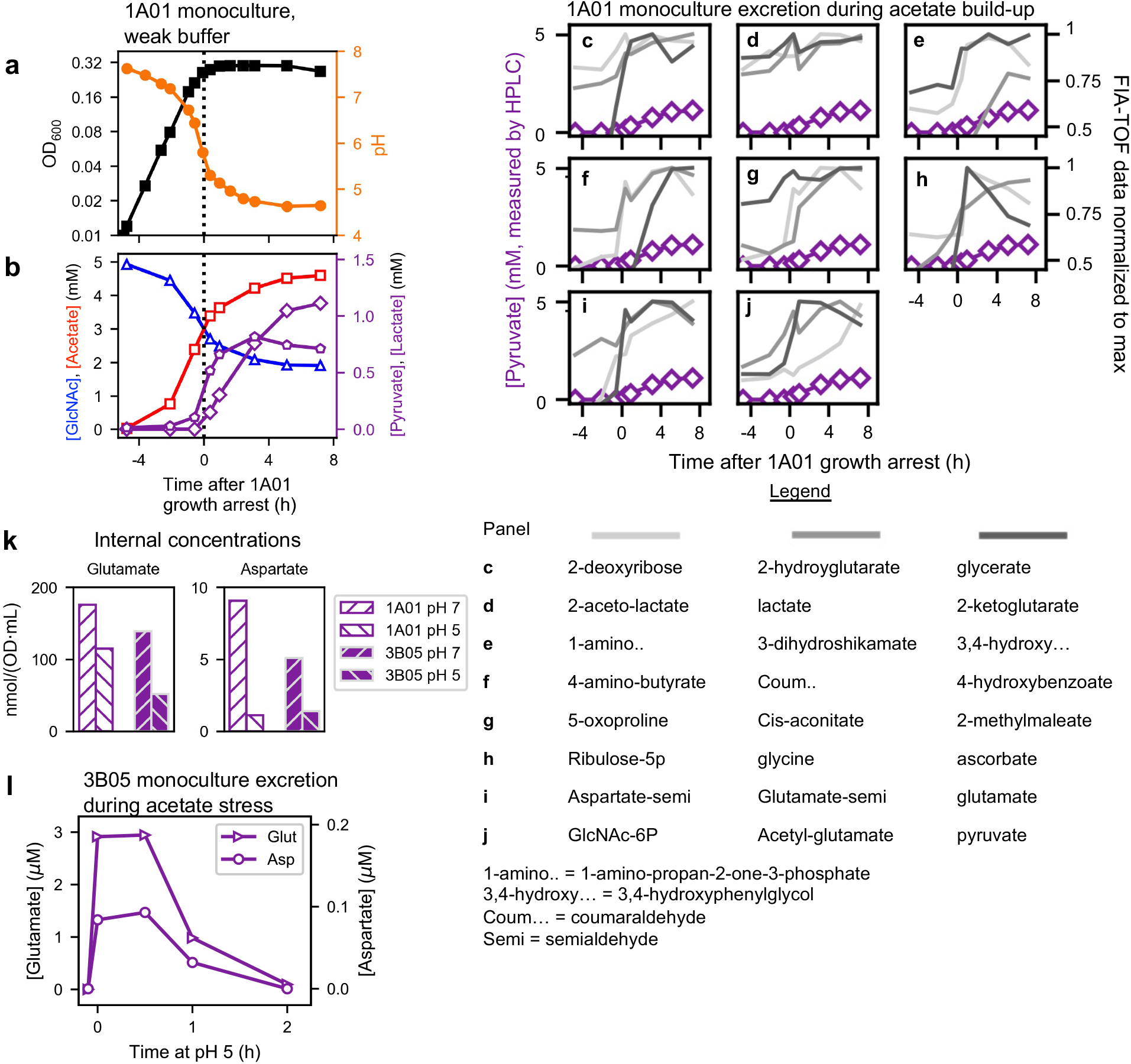
Excretion of metabolites in 1A01 monoculture during growth on GlcNAc in weak buffer. 1A01 was grown in 5 mM GlcNAc in the weak 2 mM bicarbonate buffered medium. **(a)** shows OD (squares) and pH (circles), and **(b)** shows the GlcNAc and acetate concentrations in the medium. A black dotted line indicates the point at which 1A01 stops growing. **(c-j)** FIA-QTOF-MS was applied to the same spent media samples measured in part b. The right y-axis indicates the measured intensity normalized by the max intensity during the time course for that metabolite. For reference, we additionally plotted the absolute concentration of pyruvate (purple diamonds) independently measured on the same samples using HPLC (left y-axis). **(k)** Measurement of internal concentrations of glutamate and aspartate in 1A01 and 3B05 in facile conditions (‘pH 7’) and during acid stress with lower pH (‘pH 5’). For 1A01, pH 7 cells were taken when the monoculture described in **(a)** reached OD 0.1, and pH 5 cells were taken from the same culture when the pH dropped to 5 due to the buildup of acetate. For 3B05, we took a preculture growing exponentially in acetate in strong buffer, washed the cells, and resuspended into either 10 mM acetate in the weak 2 mM bicarbonate buffered medium (pH 7) or 4.5 mM acetate in the weakly buffered medium with the pH set to 5 (pH 5). The data were taken 1 h after resuspension. **(l)** Measurement of metabolites in media before and after 3B05 was exposed to the acid stress described in **(k)**. Pyruvate and lactate concentrations were below detectable levels and are not shown.

**Extended Data Figure 5.**
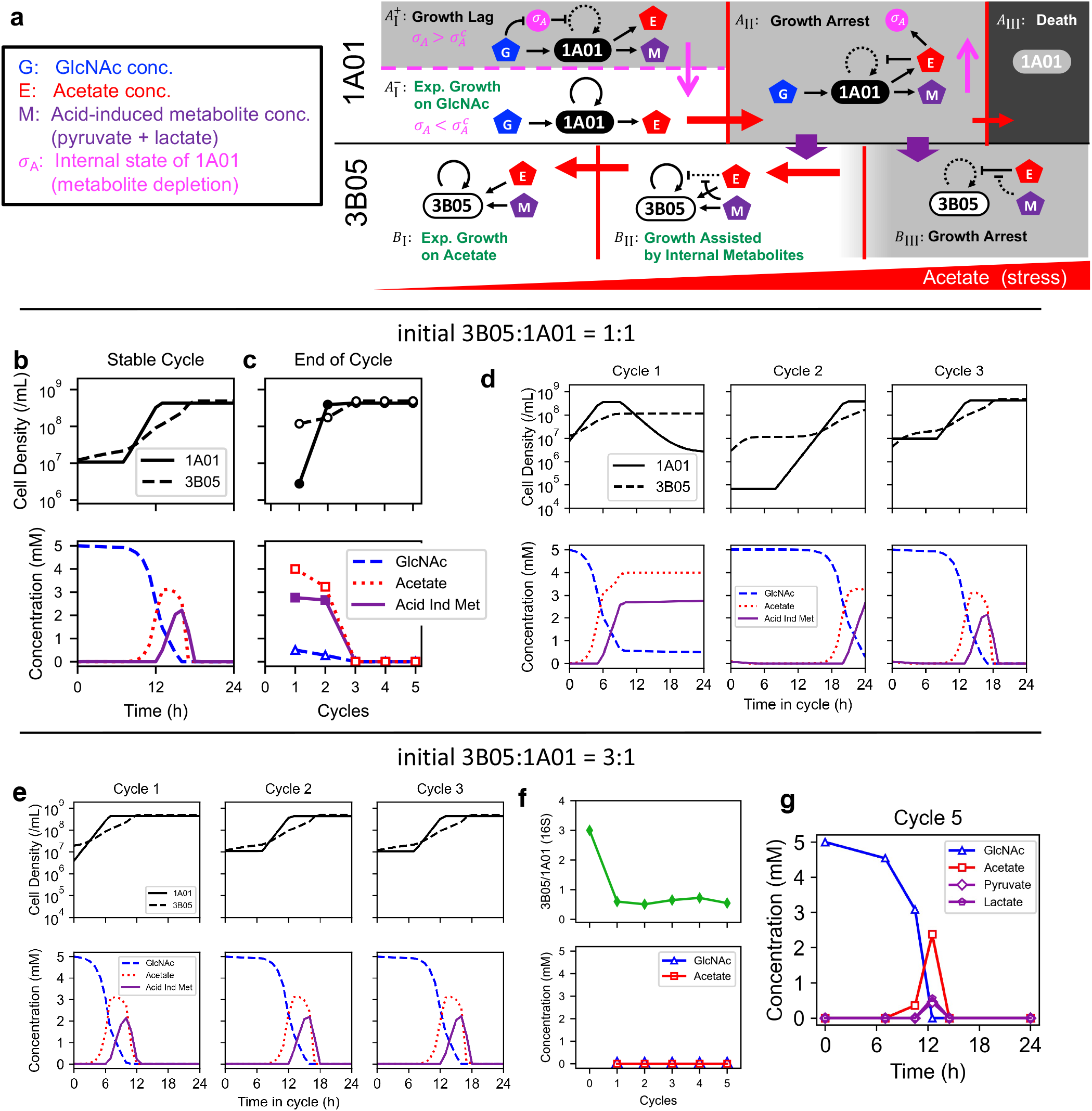
Model of acid-induced cross-feeding between 1A01 and 3B05 and the resulting population dynamics in 24-h growth-dilution cycles. **(a)** We describe the cross-feeding dynamics by a consumer-resource model outside of steady state growth. The model involves the densities of 1A01 and 3B05 cells and the concentrations of GlcNAc (G), acetate (E), and acetate-induced metabolites (M, to be interpreted as the sum of pyruvate and lactate concentrations in the medium). We also introduce an additional variable *σ*_*A*_ to describe the internal state of 1A01 due to the depletion of other metabolites such as aspartate and glutamate (**Extended Data Fig. 4i, Extended Data Fig. 6**). The key feature of our model is that the growth/death rate of the two species and the rates of uptake/excretion of the metabolites G, E, and M by the two species are dependent on the degree of acetate stress (E) and 1A01’s internal state (*σ*_*A*_). As illustrated in the schematic, we approximate these dependences by switching between several distinct forms of the rate functions depending on the values of *σ*_*A*_ and E. The rate functions corresponding to each of the four regimes for 1A01 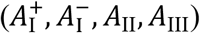, depend on both *σ*_*A*_ and E, while the three regimes for 3B05 (*B*_I_, *B*_II_, *B*_III_) depend on E. These regimes are separated by the vertical red lines and horizontal dashed magenta line; see **Supplementary Note 3** for a detailed description. In the schematic, black arrows with solid and dashed lines indicate effective and ineffective interactions in each regime. Red arrows indicate the change in the acetate concentration E, and the magenta arrows indicate the change of *σ*_*A*_. Thick purple arrows indicate the crucial cross-feeding of acetate-induced metabolites allowing 3B05 to grow during acetate stress (and hence reduce the acetate concentration in the medium). Panels **(b)** and **(c)** show numerical results of the model in (a) for the density of live 1A01 and 3B05 cells (top) and the concentration of GlcNAc, acetate, and acetate-induced metabolites (bottom) for the 24-h growth-dilution cycles with 1:1 initial species ratio. **(b)** Numerical results of the model during the stable cycle, which leads to two coordinated paths, 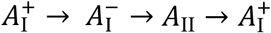 for 1A01 (solid line) and *B*_I_ → *B*_II_ → *B*_I_ for 3B05 (dashed line) over time. **(c)** Numerical results of the model at the end of each 24-hr cycle. Using the model of 1A01-3B05 cross-feeding dynamics described in **Supplementary Note 3**, we simulated the co-culture in 24h growth-dilution cycle starting with **(d)** 1:1 ratio and **(e)** 3:1 ratio of 3B05 to 1A01. The model predicts that the stable cycle is attained after two 24-h cycles for 1:1 starting ratio and after a single cycle for 3:1 starting ratio. The dynamics of the metabolites (GlcNAc, dashed line, acetate, dotted line, internal metabolites, solid line) during the stable cycle is predicted to be the same for different initial ratios. See **Supplementary Note 4** for a description of the approach to the stable cycle. The predictions for the 1:1 initial ratio captured the experimental findings shown in panels (b) and (c). The predictions for 3:1 initial ratio were tested experimentally in panels **(f)** and **(g)** with all other experimental conditions being identical to those described in the main text, i.e., for the 1A01-3B05 co-culture growing in minimal medium with 5 mM GlcNAc as the sole carbon and nitrogen source, buffered by 2 mM sodium bicarbonate, with 24h growth-dilution cycles at 40x dilution, and total initial OD = 0.02. **(f)** Ratio of 16S reads (3B05:1A01), and the concentrations of GlcNAc (blue triangles) and acetate (red squares) were measured at the end of each growth cycle. The data validated the predicted approach to the stable cycle after a single cycle. **(g)** The concentrations of GlcNAc, acetate, pyruvate (purple diamonds), and lactate (purple pentagons) on day 5 of a growth-dilution experiment that started with a 3:1 ratio of 3B05 to 1A01. The same acetate peak appeared transiently, at around t=12 h where GlcNAc was depleted, as seen in the stable cycle starting from 1:1 initial ratio (**Fig. 2b**). Thus, the same stable cycle is reached for both initial ratios of 3B05 to 1A01, despite very different transient dynamics.

**Extended Data Figure 6.**
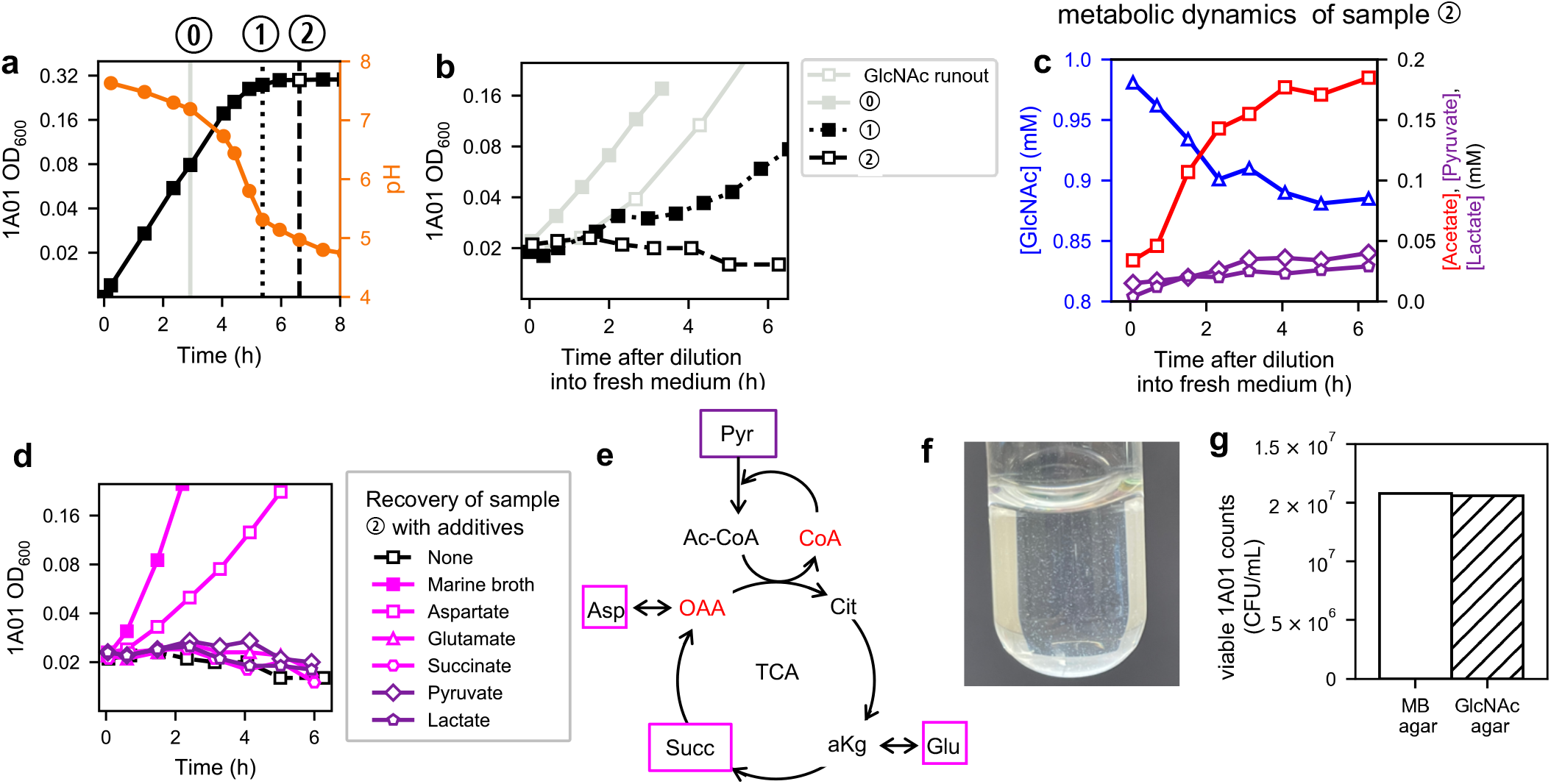
Delay in 1A01 growth recovery after experiencing acetate stress in the previous growth-dilution cycle. We hypothesize that the 6-hour lag by 1A01 after it was placed back into fresh medium at the start of the “stable cycle” (**Fig. 2a**) was due to the depletion of some of its excreted metabolites (taken up by 3B05) during the brief exposure to acetate stress in the previous cycle. To test this, we grew 1A01 monocultures in GlcNAc and weak buffer, and took the culture at various time during the self-acidification process to characterize the growth recovery dynamics upon resuspension into fresh GlcNAc medium at normal pH. **(a)** We grew 1A01 monoculture (black squares) in 5 mM GlcNAc with the weak 2 mM bicarbonate buffer, and collected samples of culture at time < 0.5 h and ∼1.5 h after growth arrest (dotted and dashed black lines, indicated by ➀ and ➁, respectively). These samples attempt to capture transient exposure to low pH (orange symbols), as well as at normal pH (solid grey line, indicated by 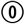). **(b)** The cells taken at different points in (a) were washed and resuspended in fresh GlcNAc medium with 2 mM bicarbonate, with pH ∼7.5. Growth recovery are shown for each case as indicated by the legend. We indeed observed a lag, whose duration increased with increased exposure to acid stress (duration and the value of pH) prior to resuspension. Open grey squares indicate a control of 1A01 cells that spent ∼12 h in strongly buffered medium following GlcNAc runout without pH dropping. **(c)** For cells taken at ➁ in (a), although the cell density did not recover during the first 6 hours (open black squares in panel b), the concentrations of GlcNAc in the medium (triangles, left axis) was being depleted over time, while Acetate, Pyruvate, and Lactate (red squares, purple diamonds, purple pentagons, respectively, right axis) accumulated in the medium. The excreted metabolites were likely what fueled the growth of 3B05 in the co-culture during the first 6-h of the stable cycle where 1A01 did not grow (**Fig. 2a**). **(d)** The lag by cells taken at ➁ was relieved if GlcNAc was supplemented by fresh marine broth (filled magenta squares), indicating that acetate-stressed 1A01 cells were not intrinsically limited from re-growth and suggested the lack of certain key metabolites. The identities of the excreted metabolites indicate that the resuspended 1A01 was still deprived of coA, which we hypothesize to result from an inability to recycle the pool of Ac-coA accumulated in the previous cycle back to coA (left panel, **Fig. 4**). The bottleneck limiting Ac-coA recycling is possibly a low pool of oxaloacetate (oaa) which catalyzes entry to TCA; see illustration in panel **(e)**. We tested this hypothesis by supplementing the resuspended cells with compounds closely related to TCA (marked in panel **(e)**): Supplementation of 5 mM Aspartate, which is linked to oaa by a single reversible transamination reaction, effectively shortened the lag (open magenta squares). In contrast, 5 mM glutamate and 5 mM succinate (triangle and circles, respectively), as well as 5 mM pyruvate and 5 mM lactate (purple markers) did not show any effect as expected by our hypothesis. **(f)-(g)** We finally describe a different aspect of the lag phase, a moderate drop of 1A01 colony count when plating the culture sampled during the first 6 hours after dilution in the stable cycle (**Fig. 2a**). This drop is due to cell aggregation, not loss of cell viability: **(f)** Picture of co-culture taken during the lag phase of the stable cycle of the 24-h growth-dilution experiment in weak buffer, 3 hours after diluting into fresh GlcNAc medium: Macroscopic aggregates are seen in the culture. Thus, aggregation of 1A01 cells may account for all or a part of the drop in 1A01 colony count observed during the lag phase. **(g)** To show that viability is not lost during the lag phase, 1A01 cells were plated onto GlcNAc minimal medium plate and marine broth plate immediately following dilution of the stable cycle. On either plate, individual cells have no chance to aggregate. If exposure to GlcNAc minimal medium was somehow toxic to 1A01, we expect to see a drop in colony count on the GlcNAc minimal medium plate compared to marine broth plate, since the latter culture grew rapidly in marine broth without lag (filled squares, **(d)**). However, we saw no difference in colony count between the two plate types. Thus, there is no death of 1A01 due to dilution into GlcNAc minimal medium.

**Extended Data Figure 7.**
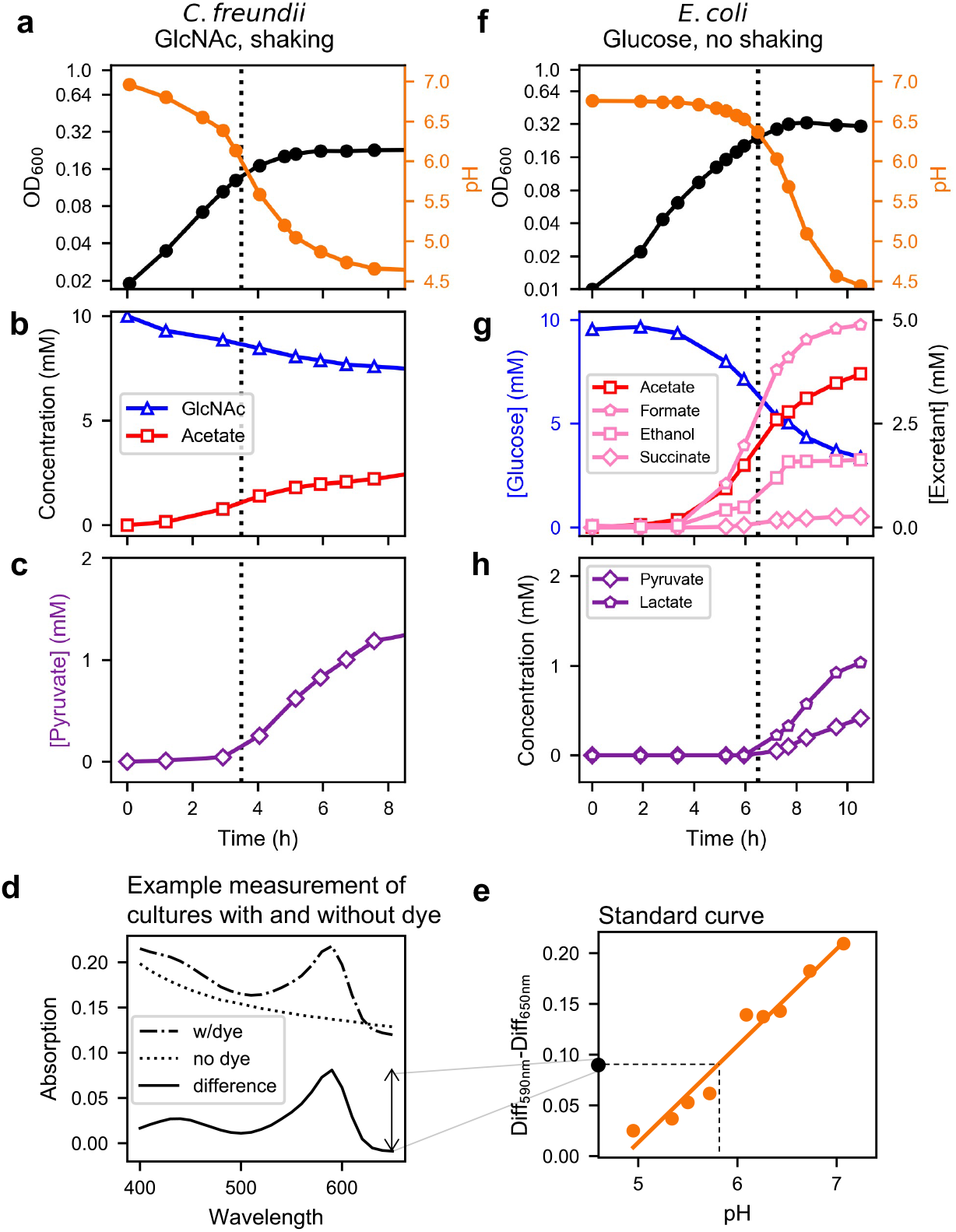
Excretion and self-acidification during monoculture growth of *Citrobacter freundii* and *Escherichia coli*. **(a)**-**(c)** Measurements during monoculture of *C. freundii* under the same conditions of the co-culture shown in **Fig. 5a-d** – 10 mM GlcNAc as the sole carbon and nitrogen sources in weak phosphate-based buffer and with shaking. **(a)** OD and pH, **(b)** consumption of GlcNAc and excretion of acetate, and **(c)** excretion of pyruvate. **(d-e)** To enable the continuous measurement of pH with a plate reader, we developed the following method. First, we initiated two cultures with identical inoculant. The only difference between the growth media in the two cultures was that one culture contained 0.0004% w/v bromocresol purple (‘with dye’) and the other did not (‘without dye’). We measured the absorption spectrum of the two cultures every 15 min as the cells grew. **(d)** To calculate the pH at a particular time point, we first subtracted the spectrum of the culture ‘with dye’ (dash dotted line) from the spectrum of the culture ‘w/o dye’ (dotted line) to remove the scattering due to the cells and get the ‘difference’ (solid line). Next, we subtracted the difference value at 650 nm from that at 590 nm to calculate the height of the peak for the basic form of the dye. **(e)** This height was then converted to a pH using a standard curve measured using growth media adjusted to different, known pH values (slope = 0.10, y-intercept = -0.46, R^2^ = 0.96). **(f)**-**(h)** Measurements during monoculture of *E. coli* under the same conditions of the co-culture shown in **Fig. 5e-h** – 10 mM Glucose as the sole carbon source and without shaking. **(f)** OD and pH, **(g)** consumption of glucose and excretion of acetate, succinate, formate, and ethanol, and **(h)** excretion of pyruvate and lactate.

