## Supplemental Information for "Dynamic metabolic exchanges between complementary bacterial types provide collaborative stress resistance"

### Supplementary Information

#### Table of Contents:


| | |
|---|---|
| **Supplementary Methods** | 2-11 |
| **Supplementary Note 1** | 12-13 |
| **Supplementary Note 2** | 14-15 |
| **Supplementary Note 3** | 16-23 |
| **Supplementary Note 4** | 24-26 |
| **Supplementary Table 1** | 27 |
| **Supplementary Table 2** | 27 |
| **Supplementary References** | 28 |

**Supplementary Methods**

### 1. Strains

*Vibrio splendidus* sp. 1A01 and *Neptunomonas phycotrophica* sp. 3B05 were natural isolates obtained by Datta et al[1]. In that work, ocean water collected near Woods Hole, MA, was mixed with chitin beads in the lab. 1A01 and 3B05 reached greater than 1% abundance on the surface of the beads at some point over the course of 6 days.

The additional strains used to test for acid-induced cross-feeding were *E. coli* NCM3722, *Citrobacter freundii* (ATCC# 8090), *Pseudomonas fluorescens* (ATCC# 13525), and *Pseudomonas putida* (ATCC# 12633).

### 2. Growth media

*2.1 Preparation of marine broth and LB agar plates.*

Marine broth medium was prepared by mixing 37.4 g of dried solid (Difco Marine Broth 2216) with $ddH_2O$ to 1-L. This solution was boiled for 1 min and allowed to cool before it was vacuum filtered through a 0.22 µm filter. The solution was stored at room temperature. Marine broth (1.5%) agar plates were prepared by mixing together 2x marine broth medium (74.8 g/L) and 2x (30 g/L) autoclaved agar on a stir/hot plate. The temperature of this solution was maintained above 50 °C to prevent any agar solidification. Fifteen mL of the 1x marine broth/1.5% agar solution was added to a petri dish (Fisherbrand, 100 mm x 15 mm). Following solidification of the agar, the plates were stored in stacks face down in sealed bags at 4 °C. LB plates were prepared the same way, except 2x LB Broth (Miller, 50 g/L) medium was combined with 2x agar.

*2.2 Preparation of 'strongly buffered' HEPES minimal growth medium.*

We prepared a growth medium inspired by that used in the Marine Biological Laboratory's Microbial Diversity Summer Course and MOPS medium used for growth of enteric bacteria such as *E. coli*[2]. The benefits of this medium are 1) it is stable and supports steady-state growth of copiotrophic, heterotrophic marine bacteria to high densities, 2) it is easily made/purchased, and 3) it is clear and thus amenable to OD measurement of biomass.

We prepared the growth medium with HEPES as the buffer ('strongly buffered medium') as follows. (i) Prepare 1 L of a 10x concentrate of by mixing the following: HEPES sodium salt, freshly prepared, 1.0 M, adjusted to pH 8.2 using 5 M HCl (400 mL); Tricine, freshly prepared, 1.0 M, adjusted to pH 7.4 with 5 M NaOH (40 mL); 1.0 M $Na_2SO_4$ (10 mL); trace metals (50 mL), a solution containing 7.6 mM $FeSO_4 \cdot 7H_2O$, 0.48 mM $H_3BO_3$, 0.8 mM $CoCl_2 \cdot 6H_2O$, 12 µM $CuSO_4$, 0.5 mM $MnCl_2 \cdot 4H_2O$, 0.5 mM $ZnSO_4 \cdot 7H_2O$, 0.15 mM $Na_2MoO_4 \cdot 2H_2O$, 0.1 mM $NiCl_2 \cdot 6H_2O$, 23 µM $SeO_2$; and adding $ddH_2O$ to the mixture to 1 L. Filter sterilize this 10x concentrate by vacuum filtration through a 0.2 µm filter and store at -20 °C. (ii) Prepare 1 L of 4x concentrate of a simplified seawater (SW) mixture: 1.37 M NaCl, 59 mM $MgCl_2 \cdot 6H_2O$, 4 mM $CaCl_2 \cdot 2H_2O$, and 27 mM KCl. Filter sterilize and store at room temperature. (iii) Prepare a carbon source (i.e., 1 M sodium acetate), 1 M $NH_4Cl$ as the nitrogen source, and 0.5 M $Na_2HPO_4$ as the phosphorus source. Filter sterilize and store at room temperature (or -20 °C in the case of 0.4 M GlcNAc). (iv) To prepare the final growth medium, add the following to make 40 mL of 5 mM glucose medium, for example: 1) 25.32 mL of autoclaved $ddH_2O$, 10 mL of 4x seawater, 0.2 mL of 1 M glucose, 0.4 mL of 1 M $NH_4Cl$, 0.08 mL of 0.5 M $Na_2HPO_4$, 4 mL of 10x C-N-P-SW-concentrate. Vortex. This medium is stable at room temperature for at least a week. For medium with GlcNAc as the sole carbon source, no ammonium is provided unless otherwise indicated.

*2.3 Preparation of bicarbonate ('weakly buffered') minimal growth medium*
We did not include tricine in the growth medium with sodium bicarbonate as the buffer since it affected buffering. Tricine was included in the HEPES buffered medium because iron would crash out upon storage at 4°C due to the pH 8 of the medium. The bicarbonate buffered medium was stable (pH 7.5, iron remained solubilized) for at least 24 h at room temperature. To prepare 40 mL of GlcNAc growth medium with 2 mM bicarbonate carbonate as the buffer and no additional nitrogen source, we added the following: 28.38 mL of autoclaved dd$H_2O$, 10 mL of 4x seawater, 0.5 mL of 0.4 M GlcNAc, 0.08 mL of 0.5 M $Na_2HPO_4$, 0.04 mL of 1 M $Na_2SO_4$, 0.2 mL of the trace metals mixture described above, and 0.8 mL of a 0.2 µm-filtered freshly prepared solution of 0.1 M $NaHCO_3$; then vortexed to mix. Note: we used this medium only after ~30 min to allow acid-base and bicarbonate equilibration with the atmosphere.

*2.4 Preparation of phosphate buffered medium*
The base minimal medium for the soil strains was a 1x M9 medium[3] (we used 10 mM $NH_4Cl$ instead of 18.7 mM) with 1x micronutrients and a carbon source. The 1000x micronutrient solution contained 20 mM $FeSO_4$, 500 mM $MgCl_2$, 1 mM $MnCl_2 \cdot 4H_2O$, 1 mM $CoCl_2 \cdot 6H_2O$, 1 mM $ZnSO_4 \cdot 7H_2O$, 1 mM $H_{24}Mo_7N_6O_{24} \cdot 4H_2O$, 1 mM $NiSO_4 \cdot 6H_2O$, 1 mM $CuSO_4 \cdot 5H_2O$, 1 mM $SeO_2$, 1 mM $H_3BO_4$, and 50 mM $CaCl_2$ dissolved in a 0.1 N HCl solution[4]. For simplicity in the remainder of the methods, we call this base minimal medium 'M9'.

For the *C. freundii-P. fluorescens* co-culture, we used M9 medium with the phosphate buffer component diluted 32x to give a buffer concentration similar to 2 mM bicarbonate. For the *E. coli-P. putida* co-culture we used M9 medium with the phosphate buffer component diluted 4x.

**3. Growth of monocultures**
*3.1 Batch monoculture growth*
All cultures (except for those grown in the plate reader, see below) were grown in a water bath shaker at 27°C with shaking at 250 rpm. We used this temperature for all growth experiments because the growth rate of 1A01 was highest at this temperature. We used this shaking frequency and the culture volumes specified below to ensure that oxygen availability was not limiting for $OD_{600} < 1.5$ (except for the *E. coli-P. putida* cocultures and *E. coli* monocultures, see sections 3.7 and 4.4). $OD_{600}$ was measured using a Thermo Scientific GENESYS 30 Spectrophotometer.

Each growth experiment involved three steps: 1) a seed culture, 2) a preculture, and 3) an experimental culture. The seed culture was started by inoculating 2 mL of marine broth medium in a 16 mm x 125 mm test tube (borosilicate glass, Fisherbrand, Cat. No. 14-961-30) from a single colony on a marine broth/agar plate. Once the seed culture saturated (which took ~7 hr for 1A01 and ~12 hr for 3B05), the cells were washed and resuspended in 1x seawater to an $OD_{600}$ of ~1 before being diluted into the experimental medium (3 mL in a 16 mm tube) for growth overnight, such that, by the following day, the preculture a) doubled ≥10 times and b) remained growing exponentially. While the preculture was still in exponential growth, we diluted the preculture into fresh experimental medium (8 mL in a 25 mm x 150 mm tube, prewarmed to 27 ºC) to an $OD_{600}$ of ~0.01. After another two doublings in the experimental culture, we took samples for various measurements, e.g., for the growth curve, spent media, etc. See Sec. 5 of **Supp Methods** for details on sample collection. Whenever cells were washed with or transferred to another medium, it should be assumed that the medium was prewarmed to 27 ºC unless otherwise indicated. Also, all wash steps were for 2 min x 7.5k rpm unless otherwise indicated.

*3.2 Growth on chitin*
For the measurement of growth on chitin (**Extended Data Fig. 1a**), we prepared 1A01 and 3B05 precultures in 10 mM GlcNAc (-N) and 60 mM acetate/10 mM $NH_4Cl$ HEPES minimal medium, respectively. We washed and resuspended the cells in C-N- HEPES minimal medium before adding to 8 mL of HEPES minimal medium with 0.2% w/v chitin flakes and 10 mM $NH_4Cl$ (Millipore Sigma, C7170) to an $OD_{600}$ of 0.05. No additional C source was provided.

*3.3 Measurement of pH-dependence of growth rate*
For **Extended Data Fig. 2c**, we precultured 1A01 and 3B05 each in 0.4% v/v glycerol HEPES minimal medium. We prepared a 96 well plate (Falcon, Product number 353072) containing 250 μL of 0.4% v/v glycerol minimal medium buffered by 10 mM MES. To vary the pH of the medium, we varied the ratio of the base and acid forms of MES and measured the pH using a Thermo Scientific Orion Star A221 pH meter. We allowed the plate to warm at 27 °C for 10 min in the plate reader before adding 1A01 or 3B05 to the wells.

To initiate growth in the plate, 2 mL of the preculture grown in HEPES medium with 0.4% glycerol was added to a well containing medium in the 96 well plate described above such that the $OD_{600}$ in the well (as measured by the plate reader) was ~0.0002. To allow for aeration of the cultures while avoiding evaporation of water and condensation on the lid, we attached the plate to its lid by lining the inside of the edges of the lid with high vacuum grease (Dow Corning). Note that we did not use grease on the corners of the lid to allow for aeration. The plate reader (Tecan Spark) was set to Orbital shaking with an amplitude of 2 mm and a frequency of 240 rpm. The $OD_{600}$ was measured every 10 min. We measured the growth rate starting at $OD_{600}$ ~ 0.03, so that the cells had ~7 doublings to acclimate to the pH. Following the cessation of growth, the volume of culture in the wells were spot checked for evaporation of water.

*3.4 Measurement of 1A01 death rate in the absence of a nutrient*
For **Extended Data Fig. 3g**, we first grew a 1A01 preculture in GlcNAc (-N) HEPES medium to $OD_{600}$ of 1. We sampled this preculture for plating as it grew exponentially. We then washed the cells twice and resuspended these cells in 18 mL of C-N- naturally buffered medium to an OD of ~1. We split this resuspension into two 25 mm tubes each with 9 mL of culture; in one of the tubes nothing was added so the pH remained at 7.5 and in the other tube we added 4 μL of 1 M acetic acid to a final concentration of 3.6 mM to lower the pH to 5.25. We then put both tubes back into the 27 °C water bath shaker and periodically sampled the cultures for $OD_{600}$ and plating (Section 5.2). Five hours after starting the cultures, we measured the pH of the two tubes again to confirm that the pH had not altered during the experiment.

*3.5 1A01 growth in manual 'pH stat'*
For **Fig. 3d-f**, we first grew 1A01 preculture in 10 mM GlcNAc (-N) HEPES medium. We washed the cells twice and resuspended the cells in 5 mM GlcNAc bicarbonate medium to an $OD_{600}$ of 0.25 in a culture volume of 20 mL in a 250 mL flask. We allowed the culture to grow and acidify

the medium through the excretion of acetate. When the pH reached ~5, we added small quantities of 0.1 M sodium bicarbonate (10-100 µL) to maintain the pH above 5. We sampled this culture for $OD_{600}$, plating, and spent medium (Section 5.2).

*3.6 3B05 in acid stress*

For **Fig. 3a-c**, we first grew a 25 mL 3B05 preculture in 60 mM sodium acetate and 10 mM $NH_4Cl$ at pH 7.3 using 40 mM MOPS buffered growth medium. At this pH and concentration of total acetate, the concentration of acetic acid in the medium is ~0.2 mM; thus, 3B05 in this preculture had some exposure to acetic acid while still growing. We washed the cells twice and resuspended into 1 mL of 2 mM HAc N- bicarbonate buffered medium to an OD of 3.7. We then diluted these cells to an $OD_{600}$ of ~0.1 to 2 x 25 mm tubes containing 8 mL of bicarbonate buffered media containing 4.5 mM of acetic acid and 1 mM of ammonium chloride. In one tube we added 1 M lactate, 0.5 M pyruvate, and 1 M glutamate to a final concentration of 1.5 mM, 1 mM, and 1 mM, respectively. We subsequently returned both tubes to the water bath shaker and periodically sampled the tubes for $OD_{600}$ and spent medium (Section 5.2).

We used a similar protocol to measure the metabolic response of 3B05 to acid stress (**Extended Data Fig. 4l**). In brief, 3B05 acetate precultures in strongly buffered medium were washed and resuspended in C-N- bicarbonate buffered medium. The resuspended cells were added to an $OD_{600}$ of ~0.1 to a 25 mm tube containing 8 mL of bicarbonate buffered media to which 4.5 mM of acetic acid and 1 mM of ammonium chloride were added. Following the addition of 3B05 cells to the acidic growth medium, the cells and spent medium were sampled periodically.

*3.7 1A01 during and after exposure to acid stress*

For **Extended Data Fig. 4** and **Extended Data Fig. 6a**, we first grew a 1A01 preculture in 5 mM GlcNAc (-N) bicarbonate buffered medium. We monitored the $OD_{600}$ and pH of this culture as it acidified the medium during the course of growth due to acetate excretion. We sampled cells and spent medium for measuring internal amino acid concentrations and metabolomics (Sec. 7.2, 7.3). The same method was used to grow *Citrobacter freundii* monoculture in **Extended Data Fig. 7ab**.

When the culture reached pH 5 (or other pH values as indicated in **Extended Data Fig. 6**), we washed and resuspended the cells in fresh C-N- bicarbonate buffered medium. We added these cells to 5 mL of fresh 5 mM GlcNAc (-N) bicarbonate buffered medium such that the $OD_{600}$ was 0.02 and returned the culture to the 27 °C shaker. We subsequently sampled this culture for $OD_{600}$ and spent medium (Section 5.2).

*3.8 Monoculture growth of E. coli*

For **Extended Data Fig. 7f-h**, *E. coli* was grown exactly as the *E. coli-P. putida* co-culture (Sec. 4.4).

### 4. Growth of cocultures

*4.1 Coculture in HEPES buffer*

As with monocultures, co-cultures of 1A01 and 3B05 were grown in a water bath shaker at 27°C with shaking at 250 rpm. Whenever cells were washed with or added to growth medium to initiate a culture, the medium was prewarmed in the water bath shaker for at least 15 min.

For the growth-dilution experiments in **Fig. 1**, we first grew 1A01 and 3B05 in marine broth for 12 hr, then washed and resuspended cells in 1x seawater. We then added each strain to 6 mL of prewarmed 5 mM GlcNAc (-N) HEPES minimal medium in a 20 mm x 150 mm tube such that the $OD_{600}$ of each strain was 0.01 (total $OD_{600}$ of 0.02). After 24 hr, we added 150 µL of this culture to 5.85 mL of prewarmed 5 mM GlcNAc (-N) HEPES minimal medium in a 20 mm x 150 mm tube. We repeated this every 24 hr until the conclusion of the experiment.

For later experiments involving coculture in HEPES buffer, we prepared 1A01 and 3B05 precultures in 10 mM GlcNAc (-N) HEPES and 60 mM acetate, 10 mM $NH_4Cl$ HEPES minimal media, respectively. We washed and resuspended each strain in 1x seawater before adding each strain to 6 mL of prewarmed 5 mM GlcNAc (-N) HEPES minimal medium in a 20 mm x 150 mm tube such that the $OD_{600}$ of each strain was 0.01 (total $OD_{600}$ of 0.02).

*4.2 Coculture in weak (bicarbonate) buffer*

We prepared precultures of 1A01 growing on 10 mM GlcNAc (-N) HEPES minimal medium and 3B05 growing on 60 mM acetate HEPES minimal medium. We washed and resuspended each of the strains into C-N- naturally buffered (2 mM bicarbonate) minimal medium. Then we added each of the strains in a 1:1 ratio to a total $OD_{600}$ of 0.02 in 6 mL of prewarmed 5 mM GlcNAc (-N) naturally buffered minimal medium in a 20 mm x 150 mm tube. Both the C-N- and 5 mM GlcNAc (-N) naturally buffered minimal media were prepared ~30 min before use. Growth-dilution experiments were propagated into fresh 5 mM GlcNAc (-N) naturally buffered minimal medium as described above. Occasionally we observed aggregation/biofilm formation following the completion of a stable cycle. In those cases, we removed any spatial structure in the tube by vigorously pipetting up and down a 500 µL volume. The stable cycle for 1:1 and 3:1 3B05:1A01 starting ratios were measured following a cycle in which there was no aggregation by eye.

For the data indicated by the open diamonds in **Fig. 1b**, we repeated the protocol described above using 1A01 and 3B05 colonies from frozen glycerol stocks of cocultures collected at the end of a previous round of 5 growth-dilution cycles.

*4.3 Mimic of stable cycle in weak (bicarbonate) buffer*

As the additional cross-feeding of internal metabolites in the stable cycle in naturally buffered medium occurs >12 hr after the start of the cycle, we developed a way to observe this using a mimic. As both strains were growing exponentially for several hours before the growth arrest, we mimicked this exponential state of the stable cycle by starting a coculture using exponentially growing 1A01 and 3B05 monocultures (in 10 mM GlcNAc and 60 mM acetate/10 mM ammonium chloride, respectively), with the density of 3B05 adjusted to that found at ~8 hr before the onset of growth arrest and the density of 1A01 kept small enough such that the following morning the coculture would reach the point 4 hr before the onset of growth arrest.

*4.4 Growth of soil cocultures*

For the *C. freundii* (Cf)-*P. fluorescens* (Pf) coculture, we prepared precultures of Cf growing on 10 mM GlcNAc M9 medium and of Pf growing on 60 mM acetate M9 medium. We washed and resuspended each of the strains into C- M9 medium with no phosphate buffer aside from 1 mM $Na_2HPO_4$. Then we added each of the strains in a 1:1 ratio to a total $OD_{600}$ of 0.02 to prewarmed 10 mM GlcNAc M9 medium with 1/32x phosphate buffer to a total volume of 200 µL in a 96 well plate (Falcon, Product number 353072) with the lid greased as described in section 3.3. (The $OD_{600}$ measured on the Thermo Scientific GENESYS 30 Spectrophotometer was equivalent to absorption at 420 nm for 200 μL of culture in the Tecan Spark plate reader.) The plate reader was set to Orbital shaking with an amplitude of 2 mm and a frequency of 240 rpm. Every 15 min the absorption from 400-650 nm was measured, in 10 nm increments. After 24 hr, 2 μL of the culture was added to 198 μL of fresh 10 mM GlcNAc M9 medium with 1/32x phosphate buffer.

For the *E. coli* (Ec)-*P. putida* (Pp) coculture, we prepared precultures of Ec growing on 10 mM Glucose M9 medium and of Pp growing on 60 mM acetate M9 medium. We washed and resuspended each of the strains into C- M9 medium with no phosphate buffer aside from 1 mM $Na_2HPO_4$. Then we added each of the strains in a 1:1 ratio to a total $OD_{600}$ of 0.02 to prewarmed 10 mM GlcNAc M9 medium with 1/4x phosphate buffer to a total volume of 200 μL in a 96 well plate (Falcon, Product number 353072), with the lid greased as described in section 3.3. The plate reader (Tecan Spark) was set to Orbital shaking with an amplitude of 2 mm and a frequency of 240 rpm for 2 min to mix the culture. Every 15 min the absorption from 400-650 nm was measured, in 10 nm increments. Between absorption measurements, the Tecan was set to "Wait," i.e. no shaking. After 24 hr, the plate was shaken for 2 min as at the start of the cycle to mix the culture, then 2 μL of culture was added to 198 μL of fresh 10 mM Glucose M9 medium with 1/4x phosphate buffer.

To monitor the pH, we added 2 μL of 0.04% bromocresol purple dye (Sigma, Product No. 114375) in a duplicate culture in a separate well at the start of the cycle. To calculate the change in the absorption spectrum of the dye, we 1) calculated the difference in absorption between the well with dye and the well without dye to remove cell background, 2) calculated the difference between the background-corrected absorption measured at 590 nm and that measured at 650 nm, and 3) calculated the pH corresponding to this value using a standard curve made using buffer solutions with known pH values (**Extended Data Fig. 7**).

### 5. Sample collection

#### *5.1 $OD_{600}$ for growth curve*

For measuring the growth curves shown in **Extended Data Fig. 1**, the $OD_{600}$ was measured by taking the tube out of the shaker, taking out 200 μL, putting the tube back into the shaker, and then pipetting the 200 μL of culture into a quartz cuvette sitting in the spectrophotometer. The reading would stabilize within 5 s. The culture spent <10 s out of the shaker for these measurements.

#### *5.2 Sampling of culture for $OD_{600}$, pH, spent medium, and plating.*

When the culture was sampled for $OD_{600}$, spent medium, and plating, the culture was taken out of the shaker, and ~700 μL was pipetted into an Eppendorf tube. The culture was then put back into the shaker within 30 s.

First, 10 mL of the culture in the Eppendorf tube was added to 990 μL of marine broth to start the dilutions for plating. Second, 200 μL of culture was used to measure the $OD_{600}$. Third, the remaining culture in the tube was centrifuged to pellet the cells for 2 min at 7.5k rpm. The supernatant (~500 μL) was then added to a Spin-X centrifuge tube containing a 0.22 μm filter (Corning Life Sciences, Costar 8169) and centrifuged for 1 min x 10k rpm. The pH was measured on either the culture used for measuring $OD_{600}$ or the filtered medium using a Thermo Scientific

Orion Star A221 pH meter. During the two centrifuge steps for collecting spent medium, subsequent dilutions into marine broth were carried out for plating, as well as the spreading of cells on the plates (see Plating section for more details). The $OD_{600}$ was measured within 1 min of collecting the culture, and the spent medium collection and pH measurement took ~6 min. Dilutions for plating were initiated ~5 min after culture collection and completed 10 min after that.

The spent medium was stored at -20°C.

*5.3 Intracellular metabolites*

We followed the protocol described by Ikeda *et al.*[5] with some modifications.

To extract intracellular metabolites from a culture, 150 μL of culture was immediately (within 10 s) added to an Eppendorf tube containing a mixture of ice cold 600 μL methanol and 30 μL of 50 μM a-amino-adipate (AAA, internal standard); the tube was vortexed for 5 s and placed into dry ice. Immediately (within 10 s), 200 μL of spent media was collected using centrifuge filtration (as described in Section 5.2). Ten μL of AAA was added to this spent medium, the mixture vortexed for 5 s, and placed into dry ice. The culture was not used following these sample collections. Both the culture and spent medium samples were stored at -80°C.

The lid of the Eppendorf containing the cell sample was opened, and the opening was covered with Parafilm, with 10-15 holes in the Parafilm made using a needle. The tube was then placed SpeedVac vacuum concentrator. The samples dried after ~3 hrs under vacuum. The resulting pellet was resuspended in 150 μL of 0.22 μm-filtered dd$H_2O$; vortexed for 30 s; heated for 1 min at 37°C; then vortexed again for 30 s. Cell debris were removed by centrifuge filter as described in Section 5.2 for the isolation of spent medium. 40 μL of this sample was used for HPLC measurement.

*5.4 Cell pellets for 16S amplification and sequencing*

For determining the ratio of the two species at the end of a growth-dilution cycle using 16S sequence amplification followed by Sanger sequencing, we first centrifuged 0.95 mL of the culture for 10 min x 7.5k rpm, as recommended by the Qiagen DNeasy Blood and Tissue Kit protocol for Gram negative bacteria. Following centrifugation, the spend medium was removed using a 1 mL pipette, with significant care taken to not dislodge the pellet. Because the pellet loosened within 30 s following centrifugation, only 2 pellets could be isolated per centrifugation to ensure the pellet composition resembled the culture composition. We note that 3B05 inefficiently pelleted as its density in the culture increased; as a result, its proportion in cocultures where its $OD_{600}$ was >0.2 was underestimated by subsequent quantification by 16S amplification and sequencing. The pellets were stored at -80 °C.

*5.5 Sampling of spent medium from 96-well plate*

For measuring spent medium from the cultures grown in 96 well plate (Sections 3.8 and 4.4), we initiated several duplicate cultures in the same plate so that each duplicate underwent the same growth dynamics. To sample a single time point of these dynamics, the entire 200 μL volume of a duplicated culture was harvested for the spent medium (Section 5.2).

### 6. Plating

As described in Section 5.2 plating was initiated by adding 10 μL of the culture to 990 μL of marine broth (either at room temperature or 27°C). This dilution was mixed by pipetting 500 μL up and down. Further dilutions were done in marine broth and by mixing the same way. One-hundred µL of the diluted culture was added onto marine broth/agar plates prewarmed to 27°C. The culture was spread using autoclaved glass beads and dried in a PCR hood until no liquid was visible on the surface of the plate. We incubated the plate in an oven at 27°C for at least 24 h. Plates with >100 and <300 colonies were counted by hand.

**7. Assays for measuring spent medium**

*7.1.1 HPLC method for measuring carbohydrates and organic acids.*

We quantified the consumption of carbohydrates and excretion of organic acids in spent medium using HPLC. We added 120 µL of spent media to a vial. The chromatographic system was a Shimadzu LC-20AB connected to a Shimadzu RID-20A refractive index detector. The auto-injector delivered a 20 µL injection volume from the vials to a Rezex ROA-Organic acid $H^+$ (8%) column (Phenomenex) kept at 40°C. The solvent system was 0.01 M $H_2SO_4$ with a flow rate of 0.4 mL/min. We analyzed the data using home-made Python code. Absolute concentrations were obtained by comparing the peak areas obtained in a sample with those from standards with known concentration.

*7.1.2 HPLC method for measuring amino acids*

The same LC system described above was used with a Gemini 5 mm C18 110 angstrom column 150 x 4.6 mm (Phenomenex) and fluorescence detection (RF-10AXL, Shimadzu). Using the LC's auto-sampler, 10 µL of the sample was derivatized with a *o*-phthaldialdehyde (OPA) solution[5] for fluorescence detection. A gradient elution for separating the derivatized amino acids was used with two solvents: solvent A was 90% sodium acetate, 9.5% methanol, and 0.5% THF set at pH 7.2; solvent B was 100% methanol. The gradient sequence was as follows: 0 to 10% of B over 6 min; 10% of B from 6 to 21.75 min; 10% to 80% of B from 21.75 to 22.5 min; 80% of B from 22.5 to 34.5 min; 80% to 0% of B from 34.5 to 35.25 min; 0% of B until 36 min (end of the run). This gradient could clearly separate aspartate, glutamate, asparagine, serine, and glutamine OPA derivatives.

The concentration of an amino acid (e.g., glutamate) in a sample was calculated by (1) calculating the ratio of the peak areas of 5 µM glutamate standard to 5 µM AAA standard; (2) calculating the ratio of the peak areas of glutamate to AAA in the sample; (3) dividing ratio (2) by ratio (1); and (4) multiplying (3) by the concentration of AAA in the sample.
To calculate the cellular amino acid content, we subtracted the amino acid amount measured in spent media sample from that in the culture sample (which contains cells and spent media).

*7.2 Enzymatic assay for ammonium concentration*

We adapted the assay procedure for L-Glutamic Dehydrogenase from Sigma[6] for measuring ammonia (ammonium) concentrations in the spent medium. We mixed the following reagents (kept ice cold until mixing):

- 700 µL of 0.1 M Tris buffer, pH 8.3
- 66.7 µL of sample (e.g., spent medium or $NH_4Cl$ standard)
- 33.3 µL of 0.225 M a-ketoglutarate, pH 7-9
- 16.7 µL of 7.5 mM NADPH

and incubated at 30°C for 5 min. Then we added 16.7 µL of a 500x dilution of L-Glutamic Dehydrogenase (Millipore Sigma, G4387-1KU) diluted with the Enzyme Diluent indicated in the protocol, mixed by pipetting up and down, aliquoted 250 µL into three wells of a 96-well plate, and started measuring absorption at 365 nm in a plate reader. The change in absorption at 365 nm after 4 min varied linearly with known ammonium chloride concentrations between 0 and 1 mM. We averaged the results of three wells to get the results for a single spent media sample.

*7.3 High throughput mass spectrometry (FIA-TOF)*
Samples were prepared for untargeted metabolomics by diluting supernatants 1:20 in water. Samples were directly injected and measured using flow-injection time-of-flight mass spectrometry (FIA-QTOF-MS). Measurements were performed using a binary LC pump (Agilent Technologies) and an MPS2 autosampler (Gerstel) coupled to an Agilent 6520 time-of-flight mass spectrometer (Agilent Technologies). Measurements were performed in negative ionization mode, at 2 Ghz for extended dynamic range, with a m/z (mass over charge ratio) range of 20-450. To reduce the matrix effects induced by high salt concentrations[7], isocratic measurements were coupled to an Agilent Poroshell 120 EC-CN column (50x2.1mm, 2.7µm). Due to the poor retention of compounds on the column, the injection peak was treated as a flow-injection approach for downstream data analysis. The mobile phase consisted of 10 mM ammonium acetate pH 5.9, and the flow rate was 250 µL/minute. 2 µL of sample was injected every 2.5 minutes. After every 30 injections, the column was washed for 5 minutes with a buffer that contains water (40%), isopropanol (30%) and acetonitrile (30%). Raw data was processed and analyzed with preprocessing raw mass spectrometry data functions contained in the bioinformatics toolbox of Matlab (The Mathworks, Natick)[8]. Ions were annotated with a tolerance of 0.005 Da against a compound library that is curated from BioCyc databases[9], which contains metabolites predicted to be present in marine bacterial isolates. 957 ions were detected, of which 124 were annotated based on the curated compound library. If a single ion is matched with multiple isomeric or isobaric compounds in the compound library, the compound that participates in the largest number of enzymatic reactions based on the BioCyc database was chosen as the top annotations.

Metabolites were first filtered to contain only those that surpass the limit of detection for at least one timepoint over the course of measurement for each timepoint represented in **Fig. 2f**, and **Extended Data Fig. 4**. This is defined as having a mean intensity that is greater than the mean intensity of a blank sample plus 3 times the standard deviation of the blank sample.
The data in **Fig. 2f** was plotted as follows. The scaled intensity for a single metabolite time course was calculated by taking the difference between the intensity and the intensity at the first time point and then dividing by the maximum such value for that time course. Metabolites for which the scaled intensity of the last time point is less than 0.5 are plotted in purple. Other detected metabolites are plotted in grey.

The data in **Extended Data Fig. 4** was plotted by dividing the intensity by the maximum intensity for a single metabolite time course.

**8 Assays for measuring coexistence**
*8.1 16S PCR and Sanger sequencing*
We isolated the genomic DNA from a pellet obtained as described in Section 5.4 using the Qiagen DNeasy Blood and Tissue Kit, using the protocol suggested for Gram negative bacteria. We amplified the 16S region using the following PCR conditions:

Primers:

- 27F: AGAGTTTGATCMTGGCTCAG
- 1492R: TACGGYTACCTTGTTACGACTT

PCR reaction:

| | Volume for 1 rxn |
|---|---|
| dd$H_2O$ | 20.8 μL |
| 5x HF buffer | 8.0 μL |
| dNTPs | 0.8 μL |
| 27F (3 μM) | 4 μL |
| 1492R (3 μM) | 4 μL |
| Template (100 ng) | 2 μL |
| Phusion polymerase | 0.4 μL |
| Total | 40 μL |

Cycling conditions:

| Step | Temperature | Time |
|---|---|---|
| **Initial denaturation** | 98°C | 30 s |
| **Amplification** | 98°C | 30 s |
| **(25 cycles)** | 54°C | 30 s |
| | 72°C | 90 s |
| **Final extension** | 72°C | 10 min |

The result of this PCR reaction was a 1506 nt product. We chose 54°C as the annealing temperature because this was the lowest temperature at which neither species gave a non-specific second band at ~1 kb. A warning: The M residue in 27F is some mixture of A and C that varies between different oligo syntheses. For one such oligo, the non-specific band at ~1 kb would not go away even with higher annealing temperatures. Thus, either A or C results in more specific binding of the 27F primer for these two species.

For a single genomic sample, we pooled together two PCR reactions and purified the PCR product using a QIAquick PCR purification kit (Qiagen). We prepared a sample for Sanger sequencing (Genewiz) by mixing 30 ng of the purified PCR product with 6 pmol of the 27F primer in 15 mL. We fit the electropherograms using the CASEU package[10] to get the fraction of each species' 16S sequence in the mixture of amplicons. With high quality electropherograms of 1A01 and 3B05, we routinely got an $R^2 > 0.9$ on fits of mixtures of amplicons using the default settings in the package.

*8.2 CFU count*

We performed the plating procedure as described in Section 6. The two strains have a different colony size and density that could be detected by eye (**Extended Data Fig. 3c-e)**.

**9. Simulations**

Numerical simulations of the models described in **Supplementary Notes** and in **Extended Data Figs. 2d,e, 5b-e** were performed using Python or Matlab.

**Supplementary Note 1: Effect of acetate built-up on the growth of co-culture**

To determine whether the co-culture of 1A01 and 3B05 (abbreviated as A and B, respectively when needed) consumes all of the carbon in weakly buffered 2 mM bicarbonate medium with 5 mM GlcNAc, we constructed a kinetic model of growth and cross-feeding (**Fig. N1a**) parameterized using measurements on the single strains (**Extended Data Table 1**).

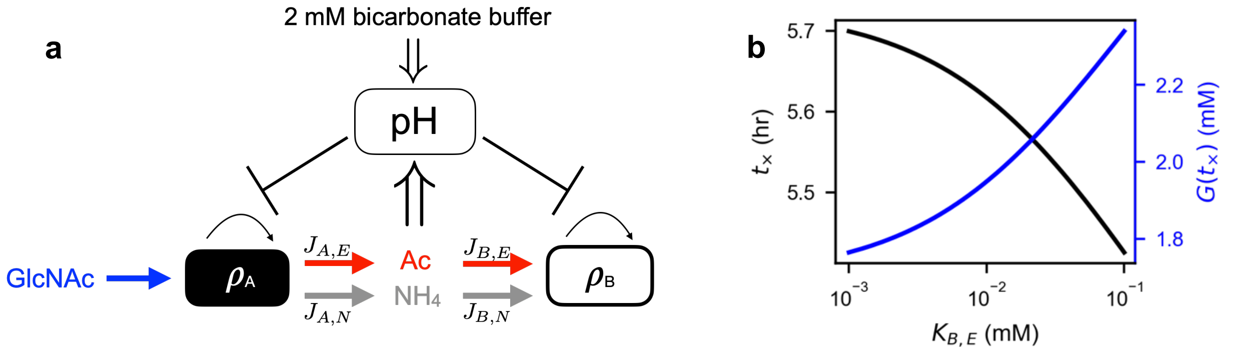


**Figure N1. Model of coculture growth with stopping due to acetate buildup. (a)** Schematic of model. The density of 1A01 ($\rho_A$) increases by taking up GlcNAc and turning it into biomass while excreting both acetate and ammonia with fluxes $J_{A,\mathrm{E}}$ and $J_{A,\mathrm{N}}$, respectively. The density of 3B05 ($\rho_B$) increases by taking up acetate and ammonia with fluxes $J_{B,\mathrm{E}}$ and $J_{B,\mathrm{N}}$ and turning it into biomass. The growth of both 1A01 and 3B05 are inhibited by acetate due to the weak buffering capacity of 2 mM bicarbonate. **(b)** Dependence of the co-culture stopping time ($t_\times$) and GlcNAc concentration at that time ($G(t_\times)$) on the Monod constant of acetate ($K_{B,E}$) for the growth of 3B05.

The growth rate of 1A01 depends on the GlcNAc concentration, $G(t)$, and the pH of the co-culture. The growth rate of 3B05 depends on the concentrations of acetate, $E(t)$, ammonium, $N(t)$, and the pH of the culture. As shown in **Extended Data Fig. 3a, 3b**, both 1A01 and 3B05 stop growing at acetate concentrations above 2.5~3 mM. Further, we note from **Extended Data Figure 2c**, that 3B05 is more sensitive to pH, and hence to increased acetate (since for our conditions, pH is a unique function of the acetate concentration, $E(t)$). For simplicity, we take the transition from growth to growth arrest to be a hard switch. Thus, for $E$ below some threshold concentrations, $E_{B1} = 2.5$ mM and $E_{A1} = 3$ mM for 3B05 and 1A01 respectively, we take the growth rates to be of the Monod form, and no growth if $E(t)$ is above the thresholds. This leads to Eqs. (1.1) & (1.2) for the growth of 1A01 and 3B05, whose densities are denoted as $\rho_A$ and $\rho_B$ respectively,

$$\frac{d\rho_A}{dt} = \begin{cases} r_A \cdot f\left(G/K_{A,G}\right) \cdot \rho_A, & E < E_{A1} \\ 0, & E \geq E_{A1} \end{cases} \tag{1.1}$$

$$\frac{d\rho_B}{dt} = \begin{cases} r_B \cdot f\left(E/K_{B,E}\right) \cdot f\left(N/K_{B,N}\right) \cdot \rho_B, & E < E_{B1} \\ 0, & E \geq E_{B1} \end{cases} \tag{1.2}$$

with $f(x) \equiv x/(1+x)$ describing the Michaelis dependence on the substrate concentration. For 3B05, we take growth to be simultaneously co-limited by the absence of either of the two substrates: acetate or ammonium. The multiplicative form for nutrient co-limitation can be thought of as a simple AND function requiring both nutrients to be present and is the predicted form for the limit of slow metabolic rates[9]. For the parameters of the Monod functions, the maximum growth rates $r_i$ are taken from the batch culture measurements reported ($r_A = 0.7/\text{h}$ and $r_B = 0.35/\text{h}$ for weak buffer; see **Extended Data Table 1)**. The Michaelis constants $K_{A,G}$, $K_{B,E}$, and $K_{B,N}$ are very small ($\ll 1$ mM) given how abruptly the cultures cease growth when each nutrient runs out. For our calculations, they are taken to be 10 μM, but their precise values do not drastically affect the results as we will discuss below.

The dynamics of the metabolite concentrations are given by Eqs. (1.3)-(1.5). The uptake and excretion fluxes of metabolite $m$ by species $i$, denoted as $J_{i,m}$ , are given by $\lambda_i \rho_i / Y_{i,m}$, where $\lambda_i \equiv r_i \cdot f(x_i/K_{i,m})$ are the actual growth rates, and $Y_{i,m}$ (in unit of OD/mM) are the yield factors or the excretion factors, whose inverse values are listed in **Extended Data Table 1**.

$$\frac{dG}{dt} = -\frac{\lambda_A \rho_A}{Y_{A,\text{G}}} \equiv -J_{A,\text{G}}, \tag{1.3}$$

$$\frac{dE}{dt} = \frac{1}{Y_{A,E}} \frac{d\rho_A}{dt} - \frac{1}{Y_{B,E}} \frac{d\rho_B}{dt} = J_{A,E} - J_{B,E}, \tag{1.4}$$

$$\frac{dN}{dt} = \frac{1}{Y_{A,N}} \frac{d\rho_A}{dt} - \frac{1}{Y_{B,N}} \frac{d\rho_B}{dt} = J_{A,\text{N}} - J_{B,\text{N}}. \tag{1.5}$$

We simulated the 1A01-3B05 co-culture using Eqs. (1.1) -(1.5), starting with a 1:1 ratio at OD = 0.02 as was done in the experiment. As shown in **Extended Data Fig. 2e**, the model predicts that after a short period, the acetate concentration increases exponentially and reaches the stopping concentration for 3B05 ($E_B$) after ~6 h. Although 1A01 has a slightly larger stopping concentration ($E_{A1}$), once 3B05 stops growing and stops taking up acetate, acetate accumulates even more rapidly and 1A01 stops growing very shortly after. So, the two strains essentially stop growing at the same acetate concentration, $E \approx E_{B1}$. At this point, ~2 mM GlcNAc remains though the co-culture has stopped growing. During the 6-h growth period, the densities of 1A01 and 3B05 are expected to increase by 64-fold and 8-fold respectively, based on their maximum growth rates, such that the ratio of 3B05 to 1A01 cells has dropped from 1:1 at the start to 1:8 when growth ceases.

As mentioned above, the values of the Monod constants $K_{A,G}$, $K_{B,N}$, and $K_{A,E}$ are not known, other than that they are well below 1 mM. To assess the sensitivity of the co-culture dynamics shown in **Extended Data Fig. 2e** to the values of these parameters, we note first that the dependence on $K_{A,G}$ and $K_{A,N}$ are completely negligible: $K_{A,G}$ is negligible because throughout the simulation the concentration of GlcNAc stayed above 1 mM which much exceeds $K_{A,G}$. $K_{B,N}$ is negligible compared to the ammonium concentrations as ammonium accumulates, indicating that it does not limit 3B05 growth in our experiments. However, $K_{B,E}$ does affect the outcome moderately. We varied $K_{B,E}$ from 1 to 100 $\mu M$ while keeping $K_{A,G}$ and $K_{B,N}$ constant. The two key outputs of the model are plotted against $K_{B,E}$, the time at which the co-culture stops due to acetate build-up, $t_\times$, and the amount of GlcNAc remaining in the medium, $G(t_\times)$ (**Fig. N1b**). The plots show that $t_\times$ changed by less than 10% (black line), with the remaining GlcNAc concentration close to ~2 mM (blue line). Thus, the dynamics of the coculture are not significantly affected by the precise values of the Monod constants.

### Supplementary Note 2. Model of growth-dilution cycles including cell death

In **Extended Data Fig. 3f-g**, we showed that 1A01 cells died rapidly when exposed to low pH in the absence of GlcNAc. Such a death would decrease the average growth rate of 1A01 over a cycle and possibly allow 3B05 to recover and de-acidify the environment. Can the death of 1A01 explain the stable coexistence the 1A01-3B05 co-culture in weak buffer (**Fig. 1e**)? In this note, we examine this possibility based on the single strain characteristics measured in **Extended Data Fig. 1** and **2c**, together with the death characteristics. Our model for the co-culture is identical to that presented **in Supplementary Note 1** except that when the acetate concentration $E$ reaches a threshold of $E_{A2} \approx 4$ mM, 1A01 starts dying with a death rate $\delta_A = 0.5/\text{h}$ (**Extended Data Fig. 3g**). This is implemented with the modification of Eq. (1.1) in **Supplementary Note 1** to the following:

$$\frac{d\rho_A}{dt} = \begin{cases} r_A \cdot f\left(G/K_{A,G}\right)\rho_A, & E < E_{A2} \\ -\delta_A \cdot \rho_A, & E \geq E_{A2}. \end{cases} \tag{2.1}$$

To simulate the growth-dilution cycles, the model now keeps track of the cycles. At the start of new cycle, all the metabolite concentrations and cell densities obtained at the end of period (24 hours) of the previous cycle were divided by the dilution factor (40-fold). In addition, 5 mM of GlcNAc was added upon dilution. The simulation results are shown in **Fig. N2a-d** and discussed below.

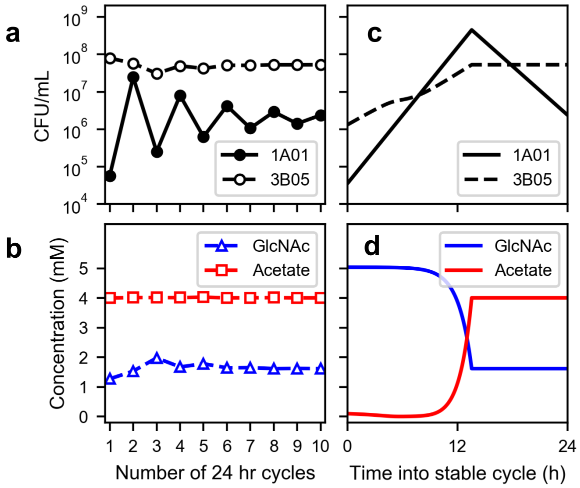


**Figure N2.** Simulation results using the model of 1A01-3B05 cross-feeding including cell death as described above, applied to 24-h growth-dilution cycles with 5mM GlcNAc at the start of the cycle and 40x dilution. The density of live 1A01 cells is indicated by a solid black line and 3B05 by a dashed black line. The concentrations of acetate and GlcNAc are indicated by the red and blue lines, respectively. **(a)** and **(b)** show the simulation results at the end of each 24-h cycle. **(c)** and **(d)** show the simulation results during Cycle 10, after the co-culture has stabilized.

The simulation results show that the model with cell death is able to produce the co-existence of 1A01 and 3B05. The effect of cell death can be understood as follows: since the growth rate for 3B05 on acetate in the weakly-buffered medium is 0.35 $h^{-1}$, it needs at least 11.8 h to grow 40x to make up for the dilution at the end of a cycle. The rapid death of 1A01 means that, following dilution, it needs to grow $\gg$ 40x to maintain its population. Thus, 1A01 must undergo additional doublings until it reaches densities that can acidify the culture. This delays acetate buildup and the ensuing stoppage of coculture growth, thereby giving 3B05 more time to replicate and catch up for the 40x density increase required for coexistence. Quantitatively, our model shows that coexistence can be achieved in a stable cycle with growth occurring for the first ~13 h (panels c and d), as opposed to the first cycle where growth was limited to the first 6 hours (**Extended Data Fig. 3f**). This results in a reduced period of growth arrest and hence a reduced drop in the viable 1A01 cells, such that in one cycle 1A01 and 3B05 each has just enough time to increase its density 40x. Thus, the death of a faster species can effectively delay the onset of toxicity, thereby enabling a slower cross-feeder to avoid extinction.

In this model, the key to the co-culture finding a stable cycle is that it stops growing prematurely, i.e., before the full consumption of GlcNAc. This point can be adjusted dynamically until a stable cycle is reached where the growth period for 1A01 is just enough to compensate for its death and the dilution factor. However, our data on GlcNAc and acetate at the end of each cycle of the 24-h growth-dilution experiment in weak buffer (**Fig. 1f**) showed major differences from the simulation result here: More acetate and less GlcNAc than predicted were found at the end of Cycle 1, and, once the co-culture stabilized, GlcNAc and acetate were entirely consumed whereas significant amount remained in the simulation (**Fig. N2b**). Also, the ratio of viable counts of 1A01 is comparable to 3B05 (right panel, **Fig. 1c**) whereas it is more than 10x lower than 3B05 in this model (**Fig. N2a**). Therefore, death of 1A01 modeled in this simulation cannot be the cause of stable coexistence in weak buffer that was observed experimentally.

**Supplementary Note 3. Dynamical Model of acid-induced cross-feeding**

The results of the simulations of the models in **Supplementary Notes 1 and 2** demonstrate that growth arrest, and even death, are not sufficient to account for the coexistence of 1A01 and 3B05, nor the complete utilization of the carbon source. The other experimental observation that allows 3B05 to recover after acidification of the environment is acid-induced metabolic cross-feeding between 1A01 and 3B05. Is acid-induced metabolic cross-feeding as has been characterized at the single-strain level in the main text (**Fig. 3**), together with the growth characteristic established in **Extended Data Figures 1, 3** and used in **Supplementary Note 1, 2**, sufficient to explain the coexistence of 1A01 and 3B05 and the complete utilization of the carbon source in the stable cycle (**Fig. 1e, 1f**)? Can we also account for the dynamical observations, e.g., the crash of the co-culture during the first 24 hours (**Extended Data Fig. 3f, 3g**), the approach to the stable cycle shown in **Figs. 1e, 1f**, and the stable cycle dynamics itself (**Fig. 2a, 2b**)? We explore these possibilities by constructing in this Note a dynamical model of acid-induced metabolic cross-feeding between 1A01 and 3B05 based on the experimentally observed phenomena and parameters reported in the main text.

In addition to the densities of 1A01 and 3B05 cells, $\rho_A(t)$ and $\rho_B(t)$, respectively, and the metabolite concentrations, GlcNAc $G(t)$ and acetate $E(t)$, as discussed above, we also track the concentration of the sum of the major acid-induced metabolites in the culture (pyruvate and lactate), denoted by $M(t)$, which are cross fed during stress (**Fig. 2e**). We do not include the ammonium concentration here for simplicity as it is not the limiting nutrient in the coculture growth of 3B05.

Our model is set up in the general form of a consumer-resource model: 1A01 takes up GlcNAc with rate $\mu_{A,G}$, excretes acetate and acid-induced metabolites with rate $\mu_{A,E}$, $\mu_{A,M}$ respectively, and grows (or dies) at rate $\lambda_A$; 3B05 takes up acetate and acid-induced metabolites at rates $\mu_{B,E}$ and $\mu_{B,M}$ respectively and grows with rate $\lambda_B$. The population dynamics are therefore described straightforwardly as:

$$\frac{d\rho_A}{dt} = \lambda_A(G; E, \sigma_A) \cdot \rho_A, \tag{3.1}$$

$$\frac{d\rho_B}{dt} = \lambda_B(E, M; E) \cdot \rho_B, \tag{3.2}$$

$$\frac{dG}{dt} = -\mu_{A,G}(G; E, \sigma_A) \cdot \rho_A, \tag{3.3}$$

$$\frac{dE}{dt} = \mu_{A,E}(G; E, \sigma_A) \cdot \rho_A - \mu_{B,E}(E, M; E) \cdot \rho_B, \tag{3.4}$$

$$\frac{dM}{dt} = \mu_{A,M}(G; E, \sigma_A) \cdot \rho_A - \mu_{B,M}(E, M; E) \cdot \rho_B. \tag{3.5}$$

However, the rate functions, i.e., $\lambda_i$ for species $i \in \{A, B\}$, and $\mu_{i,m}$ for the uptake or excretion of metabolite $m \in \{G, E, M\}$, cannot be taken to be independent of the environment, as the physiological states of the two species (which determine the growth characteristics and environmental interactions) depend on the environment. Thus, the forms of the rate functions need to account for such dependences. In Eqs. (3.1) –(3.5), we indicated the two types of dependences of the rates, $\lambda_i$ and $\mu_{i,m}$, separated by a semicolon: The value of each rate for species $i$ depends on the concentration of the metabolites it takes up: $G$ for 1A01 and $M, E$ for 3B05; these are the entries that appear before the semicolon. The entries after the semicolon indicate factors that affect the physiologies of the species. The latter factors include not only $E$, which reflects the degree of acetate stress, but also another variable $\sigma_A$, which is an internal variable reflecting the depletion of other acid-induced metabolites and which affects the growth recovery of 1A01.

The forms of the rate functions $\lambda_i$ and $\mu_{i.m}$ will be described in detail below, together with the dynamics of $\sigma_A$ and the values of the parameters used. To simulate the growth-dilution cycles, Eqs. (3.1)–(3.5) are integrated numerically using the Finite Difference method and a step size of 0.36 seconds, starting with the initial conditions for the metabolites ($G(0) = 5$ mM, $E(0) = 0, M(0) = 0$) and the specified initial ratio of the two species (with $\rho_A(0) + \rho_B(0) = 0.02$ OD). After integrating these equations for 24 h, all the metabolite concentrations and the species densities are divided by the dilution factor (40-fold) and 5 mM is added to the GlcNAc concentration. The integration for the 24 h and the dilution at the end of the cycle is then repeated for five cycles. The simulation results for the viable cell density (cell/mL) reported in **Extended Data Fig. 5** were calculated by multiplying the OD from the simulation with the conversion factors (CFU/OD/mL) in **Extended Data Table 1**.

**Forms of the rate functions for 3B05.** In our experiments, 3B05 exhibited three distinct forms of growth depending on the environmental acidity and nutrient supplement as we describe below.

1. Under normal pH, 3B05 can grow on either acetate alone or on acetate, pyruvate, and lactate, with faster growth when pyruvate and lactate are present (**Extended Data Fig. 1f**). We describe these dependences by a simple sum of Monod growth functions, i.e.,

$$\lambda_B = r_{B,E} f\left(E/K_{B,E}\right) + r_{B,M} f\left(M/K_{B,M}\right), \quad \text{at normal pH} \qquad (3.6a)$$

where $r_{B,E}$ and $r_{B,M}$ are the maximum growth rate of 3B05 on acetate and on the acid-induced metabolites alone respectively, $f(x) \equiv x/(1+x)$ describes the Michaelis dependence on the substrate concentration, and $K_{B,E}$, $K_{B,M}$ are the respective Monod constants. The value of $r_{B,E}$ is taken from measurements (black squares in **Extended Data Fig. 1f**; see also **Extended Data Table 1**) to be 0.35/h. $r_{B,M}$ is taken to be 0.2/h, such that the high growth rate of 0.55/h observed for growth on acetate, pyruvate, and lactate (purple squares in **Extended Data Fig. 1f**) can be explained by additive growth on both substrates.

As discussed in **Supplementary Note 1**, the Monod constants are very small given how abruptly the cultures cease growth when each nutrient runs out. For our calculations, they are taken to be 10 µM but the results of this model are again found to be insensitive to the values of the Monod constants as the key metabolic processes occur when the concentrations of the metabolites are well above the experimentally determined upper limits of these constants. In this regime, the nutrient uptake rates by 3B05 are simply

$$\mu_{B,E} = r_{B,E} f\big(E/K_{B,E}\big)/Y_{B,E}, \tag{3.6b}$$

$$\mu_{B,M} = r_{B,M} f\big(M/K_{B,M}\big)/Y_{B,M}, \tag{3.6c}$$

with the mass conservation condition $\mu_{B,E} \cdot Y_{B,E} + \mu_{B,M} \cdot Y_{B,M} = \lambda_B$. The value of $Y_{B,E}$ was determined experimentally (**Extended Data Table 1**). We take $Y_{B,M} = \frac{3}{2} Y_{B,E}$ as each molecule of acetate has 2 carbon atoms, while each molecule of pyruvate and lactate has 3 carbon atoms.

2. At low pH, 3B05 does not grow even when acetate or other metabolites are abundant; see e.g., **Fig. 3a**. Thus, we take

$$\lambda_B = \mu_{B,E} = \mu_{B,M} = 0 \quad \text{at low pH.} \tag{3.7}$$

3. Crucially for deacidification, there exists an intermediate regime where 3B05 can grow on acetate, but only with the supplement of pyruvate and lactate (**Fig. 3a-c**). We model this simultaneous colimitation on acetate and the cross-fed metabolites by a product of Monod functions, i.e.,

$$\lambda_B = r_B^+ f\big(E/K_{B,E}\big) \cdot f\big(M/K_{B,M}\big), \quad \text{at intermediate pH,} \tag{3.8a}$$

where $r_B^+$ is the growth rate in this phase. As with Eq. (1.2), The multiplicative form for nutrient colimitation is a simple continuous AND function requiring both nutrients to be present and is the predicted form for the limit of slow metabolic rates[11]. The range of possible values of $r_B^+$ is discussed below. In this intermediate regime, the nutrient uptake rates take on the form

$$\mu_{B,E} = b\, r_B^+ f\big(E/K_{B,E}\big) f\big(M/K_{B,M}\big)/Y_{B,E}, \tag{3.8b}$$

$$\mu_{B,M} = (1-b)\, r_B^+ f\big(E/K_{B,E}\big) f\big(M/K_{B,M}\big)/Y_{B,M}, \tag{3.8c}$$

such that the mass conservation condition $\mu_{B,E} \cdot Y_{B,E} + \mu_{B,M} \cdot Y_{B,M} = \lambda_B$ still holds. The factor $b$ reflects the fraction of carbon flux that 3B05 derives for its biomass from acetate. Plotting the OD and metabolite date shown in **Fig. 3a, b** and comparing to the carbon yield of 3B05 on acetate alone (**Extended Data Table 1**), we estimate $b \approx 0.75$.

A key challenge in completing the model is to specify quantitatively how the system transitions from one regime of pH to another. As this information is difficult to obtain experimentally (requiring the setting of acetate to different fixed levels while 3B05 consumes it), we used simplified forms for transitions as already described in **Supplementary Note 1**: The regime of normal pH applies for acetate concentration below a threshold value, $E_{B1}$. However, for $E > E_{B1}$, the transition between the intermediate and low pH regimes is much more complicated and possibly path-dependent, or even density-dependent. To recover the experimental phenomenological results, we implement this transition by a soft cutoff, $\theta(E_{B2} - E) \equiv \frac{1}{2}(1 + \tanh((E_{B2} - E)/\Delta E))$, around a second threshold, $E_{B2}$, such that for $E \gg E_{B2}$, the rates $\lambda_B$

and $\mu_{B,m}$ all vanish. A tanh function was chosen for functional simplicity, and the parameter values for $r_B^+$, $E_{B2}$, and $\Delta E$ were taken to be round numbers in the range determined by experimental constraints. For example, the dynamic nature of the data in **Fig. 3a-c** does not allow extraction of a steady-state growth rate, but help us to place a bound above ~0.1/h. We also see from the data in **Fig. 2c** that the growth of 3B05 slowed down from pre-stressed growth (0.35/h, dot-dashed line) when approaching the acetate peak before picking up past the peak. Thus, we constrain the parameter $r_B^+$ to the range between 0.1/h and 0.35/h and we take $r_B^+ = 0.25/\text{h}$. Similarly, in practice we found the transition width $\Delta E$ can be quite small and in our simulations, we use $\Delta E =$ 0.2 mM, which is only 20% of the difference between the threshold values $E_{B2} - E_{B1} = 1$ mM. Altogether, the rate functions in the different regimes are summarized in **Table N3.1**. We shall discuss the values of the thresholds $E_{B1}, E_{B2}$ in conjunction with similar thresholds for 1A01 after describing the rate functions for 1A01 below.

| Model for 3B05 | $E \le E_{B1}$ | $E \ge E_{B1}$ | $E \gg E_{B2}$ |
|---|---|---|---|
| $\lambda_B$ | $r_{B,E} f\left(\frac{E}{K_E}\right) + r_{B,M} f\left(\frac{M}{K_M}\right)$ | $r_B^+ f\left(\frac{E}{K_E}\right) f\left(\frac{M}{K_M}\right) \theta(E_{B2} - E)$ | $\rightarrow 0$ |
| $\mu_{B,E}$ | $r_{B,E} f\left(\frac{E}{K_E}\right) / Y_{B,E}$ | $b r_B^+ f\left(\frac{E}{K_E}\right) f\left(\frac{M}{K_M}\right) \theta(E_{B2} - E) / Y_{B,E}$ | $\rightarrow 0$ |
| $\mu_{B,M}$ | $r_{B,M} f\left(\frac{M}{K_M}\right) / Y_{B,M}$ | $(1-b) r_B^+ f\left(\frac{E}{K_E}\right) f\left(\frac{M}{K_M}\right) \theta(E_{B2} - E) / Y_{B,M}$ | $\rightarrow 0$ |

**Table N3.1**: Forms of the growth rate of 3B05, $\lambda_B$, and its uptake rate for acetate and pyruvate/lactate, $\mu_{B,E}$ and $\mu_{B,M}$, respectively.

**Forms of the rate functions for 1A01.** The growth of 1A01 is found to depend on both the acetate concentration ($E$) and the internal state of the cells, $\sigma_A$. Here we describe the forms of the rate functions $\lambda_A$ and $\mu_{A,m}$ for 1A01 in our model, which we take to be in four possible states.

1. At normal pH, 1A01 is in exponential phase. Thus, it grows on GlcNAc with a Michaelis dependence on the GlcNAc concentration and $K_{A,G}$ being the Monod constant; the latter is again taken to be 10 μM, with the results of the model insensitive to the exact value used (see above and **Supplementary Note 1**). Thus,

$$\lambda_{A,G} = r_A f(G/K_{A,G}), \quad \text{at normal pH.} \tag{3.9}$$

In this regime, the nutrient uptake rate by 1A01 ($\mu_{A,G}$) is simply given by mass conservation ($\lambda_{A,G} = Y_{A,G}\mu_{A,G}$) and acetate is excreted by 1A01 as a by-product with excretion rate given by $\mu_{A,E} = \lambda_{A,G}/Y_{A,\text{E}}$. No acid-induced metabolites are excreted in this state. We take $r_A = 0.7/\text{h}$, $Y_{A,G}^{-1} = 6$ mM/OD, and $Y_{A,E}^{-1} = 9.4$ mM/OD based on data in **Extended Data Table 1**. Thus,

$$\mu_{A,G} = -r_A f(G/K_G)/Y_{A,\mathrm{G}}, \tag{3.10a}$$

$$\mu_{A,E} = r_A f(G/K_G)/Y_{A,\mathrm{E}}, \tag{3.10b}$$

$$\mu_{A,M} = 0. \tag{3.10c}$$

2. At very low pH, we find experimentally that 1A01 dies at a constant rate, and there is no accompanying substrate consumption or production. We take $\delta_A = 0.5/\mathrm{h}$ based on the data in **Extended Data Fig. 3f,g**. Thus,

$$\lambda_{A,G} = -\delta_A, \text{ at very low pH} \tag{3.11a}$$

$$\mu_{A,G} = \mu_{A,E} = \mu_{A,M} = 0. \tag{3.11b}$$

3. At intermediate pH values, we find that there is no growth or death of 1A01 (**Fig. 3d**), i.e.,

$$\lambda_{A,G} = 0, \text{ at intermediate pH.} \tag{3.12}$$

However, in this pH range, 1A01 cells excrete acid-induced metabolites while consuming GlcNAc (and still producing acetate as a by-product); see **Fig. 3e,f**. We model this excretion process by the same Michaelis dependence on GlcNAc concentration:

$$\mu_{A,G} = \mu_{A,G}^{str} f(G/K_G), \tag{3.13a}$$

$$\mu_{A,E} = \mu_{A,\mathrm{E}}^{str} f(G/K_G), \tag{3.13b}$$

$$\mu_{A,M} = \mu_{A,\mathrm{M}}^{str} f(G/K_G). \tag{3.13c}$$

The values of $\mu_{A,G}^{str}$, $\mu_{A,E}^{str}$, and $\mu_{A,M}^{str}$ were determined from the **Fig. 3d-f**.

4. However, 1A01 cells do not always grow even in the normal pH regime (**Fig. 2a)**. For 1A01 cells that have experienced acetate shock, we find a substantial lag phase of at least 6 hours which likely results from the depletion of key metabolites not related to pyruvate and lactate (**Extended Data Fig. 6**). To model the effect of the latter, we introduce one additional variable that describes the internal state (of metabolite depletion), $\sigma_A$, whose value is normalized to be between 0 and 1. We can think of high values of $\sigma_A$ as corresponding the case of severe depletion of these additional metabolites, and low values of $\sigma_A$ to correspond to the normal state. Thus, 1A01 cells that have recently experienced acetate stress and are in the process of recovering from acetate stress (in which case $\sigma_A > \sigma_A^c$) do not grow even if the pH is normal:

$$\lambda_{A,G} = 0, \text{ for } \sigma_A > \sigma_A^c. \tag{3.14}$$

In this state, 1A01 cells still excrete pyruvate and lactate while consuming GlcNAc (and still excretes acetate as a by-product); see **Extended Data Fig. 6c**. We model the excretion process here again with a Michaelis dependence on GlcNAc concentration:

$$\mu_{A,G} = \mu_{A,G}^{lag} f(G/K_{A,G}), \tag{3.15a}$$

$$\mu_{A,E} = \mu_{A,\mathrm{E}}^{lag} f(G/K_{A,G}), \tag{3.15b}$$

$$\mu_{A,M} = \mu_{A,\mathrm{M}}^{lag} f(G/K_{A,G}). \tag{3.15c}$$

The values of $\mu_{A,G}^{lag}$, $\mu_{A,E}^{lag}$, and $\mu_{A,M}^{lag}$ were determined from **Extended Data Fig. 6a-c** during the growth lag. It must be noted that our results hold even if we take $\mu_{A,m}^{lag} = \mu_{A,m}^{str}$ for each metabolite $m$, and we distinguish between the two values to maintain fidelity with the reported experimental results.

All the rate functions pertaining to 1A01 are summarized in **Table N3.2**, with the transition between normal, intermediate, and low pH occurring at a threshold $E_{A1}$ and $E_{A2}$.

| Model for 1A01 | $E \le E_{A1}$ | | $E_{A1} \le E < E_{A2}$ | $E \ge E_{A2}$ |
|---|---|---|---|---|
| | $\sigma_A > \sigma_\mathrm{A}^c$ | $\sigma_A \le \sigma_\mathrm{A}^c$ | | |
| $\boldsymbol{\lambda_A}$ | $r_A f(G/K_{A,G})$ | 0 | 0 | $-\delta_A$ |
| $\boldsymbol{\mu_{A,G}}$ | $r_A f(G/K_{A,G})/Y_{A,G}$ | $\mu_{A,G}^{lag} f(G/K_{A,G})$ | $\mu_{A,G}^{str} f(G/K_{A,G})$ | 0 |
| $\boldsymbol{\mu_{A,E}}$ | $r_A f(G/K_{A,G})/Y_{A,E}$ | $\mu_{A,E}^{lag} f(G/K_{A,G})$ | $\mu_{A,E}^{str} f(G/K_{A,G})$ | 0 |
| $\boldsymbol{\mu_{A,M}}$ | 0 | $\mu_{A,M}^{lag} f(G/K_{A,G})$ | $\mu_{A,M}^{str} f(G/K_{A,G})$ | 0 |

**Table N3.2:** Forms of the growth rate of 1A01, $\lambda_A$, its uptake rate for GlcNAc ($\mu_{A,G}$), and its excretion rate for acetate and pyruvate/lactate, $\mu_{A,E}$ and $\mu_{A,M}$, respectively.

So what determines the transitions between the different physiological states enumerated above? As discussed in **Supplementary Note 1**, both 1A01 and 3B05 stop growing (with 3B05 being more sensitive to acetate concentrations) above 2.5-3 mM, (based on data shown in **Extended Data Fig. 2c** and **Extended Data Fig. 3f**). Accordingly, $E_{A1}$was taken to be 3mM and $E_{B2}$ was taken to be 2.5 mM. For $E_{A2}$, we note that as shown in **Extended Data Fig. 3f**, 1A01 starts dying around 9-10 hours, which corresponds to around 4 mM of acetate as shown in **Extended Data Fig. 3b**. Thus, we take $E_{A2}$ to be 4 mM. As discussed above, the cessation of growth of 3B05 in the presence of pyruvate and lactate is much more complicated, and we only attempt to recover the phenomenological features in our model with a constant rate and a simple tanh-like switch. However, from **Fig. 2c**, we know that 3B05 can still grow and clear acetate even after the growth arrest of 1A01; thus $E_{B2} > E_{A1}$. But from **Extended Data Fig. 3b**, we note that if acetate concentrations are around the concentration where 1A01 starts dying ($E_{A2}$), 3B05 stops growing and is ineffective in clearing acetate; thus $E_{B2} < E_{A2}$. In our model, we simply take $E_{B2}$ to be at the mid-point of $E_{A1}$ and $E_{A2}$, i.e., 3.5 mM. We note that all of the hard transitions of our model can be softened without any qualitative differences in the results, and we attempted to use hard transitions as much as possible for parameter frugality and model simplicity.

**Dynamics of the internal variable $\boldsymbol{\sigma_A}$.** The last factor determining transitions in physiological states for 1A01 is the internal state variable, $\sigma_A$. There is not much experimental data to base on here since the cause of the lag in growth arrest is not fully worked out. We do know that the cause is acetate stress, which occurs for $E > E_{A1}$, and the duration of the lag is proportional to the exposure to acetate. As a first approximation, we take the increase of $\sigma_A$ to be proportional to the acetate concentration integrated over time, with a proportionality constant $\sigma_A^{str}$, such that higher stress over longer times lead to a higher value of $\sigma_A$. Further, once the acetate concentration falls, i.e., $E \leq E_{A1}$, 1A01 slowly recovers if GlcNAc is present. We take the recovery dynamics of $\sigma_A$ to have a Michaelis-Menten dependence on GlcNAc concentration with a proportionality constant $\sigma_A^{lag}$.

The value of $\sigma_A^{str}$ was chosen to ensure that $\sigma_A$ reached a high enough value during acetate stress, and subsequently the values of $\sigma_A^{lag}$ and $\sigma_A^{c}$ were chosen such that the total lag time was between 6 and 10 hours as found in **Fig. 2a**. In the simulation, we used rounded parameter values that correspond to a lag time of 8 hours. The results of the simulations did not depend on the exact values of the parameters chosen as long as the resulting lag time was in the range of 4 to 12 hours.

$$\frac{d\sigma_A}{dt} = \begin{cases} \sigma_A^{str}\, E, & E_{A1} < E \text{ and } \sigma_A < 1 \\ -\sigma_A^{lag} f\left(G/K_{A,G}\right), & E \leq E_{A1} \text{ and } \sigma_A > 0 \\ 0, & \sigma_A \geq 1 \text{ or } \sigma_A \leq 0. \end{cases} \quad (3.16)$$

Our complete model is described by Eqs. (3.1)-(3.5) and Eq. (3.16), with the rate functions defined by the entries in **Table N3.1** for 1A01 and **Table N3.2** for 3B05, and with the parameters used for the simulations specified in **Table N3.3**. As discussed above, most of the parameters are either fixed by our experimental results (as is the case for most of the maximal growth rates, the yield parameters, the threshold acetate values, etc.), or do not affect the results significantly (as is the case for the Monod constants, $\sigma_A^{str}$, etc.) However, as we described following Eqs. (3.8a-c), the parameters associated with the important metabolite-assisted growth of 3B05 at intermediate pH, $r_B^+$, $E_{B2}$, and $\Delta E$, were only loosely constrained by experiments. In our simulation, we chose these parameters to take on rounded values that lie within the experimental constraints and recover the observed behaviors, such that 3B05 is able to recover 1A01 in the stable cycle, but is unable to do so in the transient first cycle of a 1:1 initial ratio coculture. The ability of our model to recover the salient features of both the stable cycle and the transient dynamics with a rough approximation for a complex process (metabolite-induced growth under stress) indicates that the emergent qualitative and quantitative features of the model are not very sensitive to model details. However, further experimental studies will be required to elucidate how internal metabolite-assisted growth takes place under acetate stress, to establish the form of the growth function and its associated parameters (Eqs. (3.8a-c)) which are currently proposed phenomenologically.

| Parameter | Value used | Description | Source |
|---|---|---|---|
| $\boldsymbol{r_A}$ | 0.7/h | Growth rate of 1A01 on GlcNAc | ED Table 1 |
| $\boldsymbol{\mu_{A,M}^{str}}$ | 1.7 mM/OD/h | Rate of excretion of pyruvate by 1A01 during acetate stress | Fig. 3 |
| $\boldsymbol{\mu_{A,G}^{str}}$ | 1.0 mM/OD/h | Rate of GlcNAc consumption by 1A01 during acetate stress | Fig. 3 |
| $\boldsymbol{\mu_{A,E}^{str}}$ | 1.1 mM/OD/h | Rate of acetate excretion by 1A01 during acetate stress | Fig. 3 |
| $\boldsymbol{\mu_{A,M}^{lag}}$ | 0.81 mM/OD/h | Rate of pyruvate excretion by 1A01 during growth lag | ED Fig. 6a-c |
| $\boldsymbol{\mu_{A,G}^{lag}}$ | 0.81 mM/OD/h | Rate of GlcNAc consumption by 1A01 during growth lag | ED Fig. 6a-c |
| $\boldsymbol{\mu_{A,E}^{lag}}$ | 1.2 mM/OD/h | Rate of acetate excretion by 1A01 during growth lag | ED Fig. 6a-c |
| $\boldsymbol{\delta_A}$ | 0.5/h | Death rate of 1A01 at high acetate stress | ED Fig. 3g |
| $\boldsymbol{r_{B,M}}$ | 0.2/h | Maximal growth rate of 3B05 on acid-induced metabolites | difference of the two rates shown in ED Fig. 1f |
| $\boldsymbol{r_{B,E}}$ | 0.35/h | Maximal growth rate of 3B05 on acetate | ED Table 1 |
| $\boldsymbol{r_B^+}$ | 0.25/h | Maximal growth rate of 3B05 on acetate and acid-induced metabolites during acetate stress | Fig. 2c |
| $\boldsymbol{\sigma_A^{str}}$ | 0.5/mM/h | Proportionality constant for increase of internal variable of 1A01 during acetate stress | chosen to account for observed lag time in Fig. 2a |
| $\boldsymbol{\sigma_A^{lag}}$ | 0.1/h | Proportionality constant for decrease of internal variable of 1A01 during recovery from acetate stress | |
| $\boldsymbol{\sigma_c}$ | 0.1 | Threshold value of internal variable above which growth does not take place for 1A01 | |
| $\boldsymbol{E_{A1}}$ | 3 mM | Threshold acetate concentration for 1A01 to experience acetate stress and excrete acid-induced metabolites rapidly | ED Fig. 2c |
| $\boldsymbol{E_{A2}}$ | 4 mM | Threshold acetate concentration above which 1A01 dies | ED Fig. 3f, 3b |
| $\boldsymbol{E_{B1}}$ | 2.5 mM | Threshold acetate concentration for 3B05 to be unable to use acetate and acid-induced metabolites independently | ED Fig. 2c |
| $\boldsymbol{E_{B2}}$ | 3.5 mM | Soft threshold acetate concentration for 3B05 growth, even with both acetate and acid-induced metabolites | Model Parameter |
| $\boldsymbol{\Delta E}$ | 0.2 mM | Slope of the regulatory function for switch of 3B05 around $E_{B2}$ from growth to lack of growth | Model Parameter |
| $\boldsymbol{K_{A,G}}$ | 10 µM | Monod constant for growth of 1A01 on GlcNAc | See Supp. Note 1 |
| $\boldsymbol{K_{B,E}}$ | 10 µM | Monod constant for growth of 3B05 on Acetate | See Supp. Note 1 |
| $\boldsymbol{K_{B,M}}$ | 10 µM | Monod constant for growth of 3B05 on acid-induced metabolites | See Supp. Note 1 |
| $\boldsymbol{Y_{B,E}^{-1}}$ | 32 mM/OD | Amount of acetate consumed by 1 OD of growth by 3B05 | ED Table 1 |
| $\boldsymbol{Y_{A,E}^{-1}}$ | 9.4 mM/OD | Amount of acetate secreted during 1 OD of growth by 1A01 | ED Table 1 |
| $\boldsymbol{Y_{A,G}^{-1}}$ | 6 mM/OD | Amount of GlcNAc consumed by 1 OD of growth by 1A01 | ED Table 1 |
| $\boldsymbol{Y_{B,M}^{-1}}$ | 21 mM/OD | Amount of acid-induced metabolites consumed by 1 OD of growth by 3B05 | ED Table 1 & stoichiometric constraints |
| $\boldsymbol{b}$ | 0.75 | Fraction of carbon flux that 3B05 derives for its biomass from acetate | Fig. 3a, b and ED Table 1 |

**Table N3.3.** Cross-feeding simulation parameters for *V. splendidus* sp. 1A01, *N. phycotrophica* sp. 3B05 as detailed in **Supp. Note 3**.

**Supplementary Note 4. Dynamics of the stable cycle and its approach from initial conditions.**

We now describe the resulting dynamics of our model of environment-dependent cross-feeding, as defined by Eqs. (3.1)-(3.5) and (3.16) in **Supplementary Note 3** (with the rate functions given by **Table N3.1** and **N3.2**, and the parameters given by **Table N3.3**), from numerical simulations. Outputs of the model are shown in **Extended Data Fig. 5** for two initial strain density ratios 3B05:1A01=1:1 and 3:1. We describe here the dynamics of the stable cycle and the transient dynamics for each initial ratio leading to the stable cycle. We also illustrate the dynamics of the internal variable $\sigma_A$ used in the model to capture the effect of metabolite depletion on growth recovery by 1A01 (**Supplementary Note 3**).

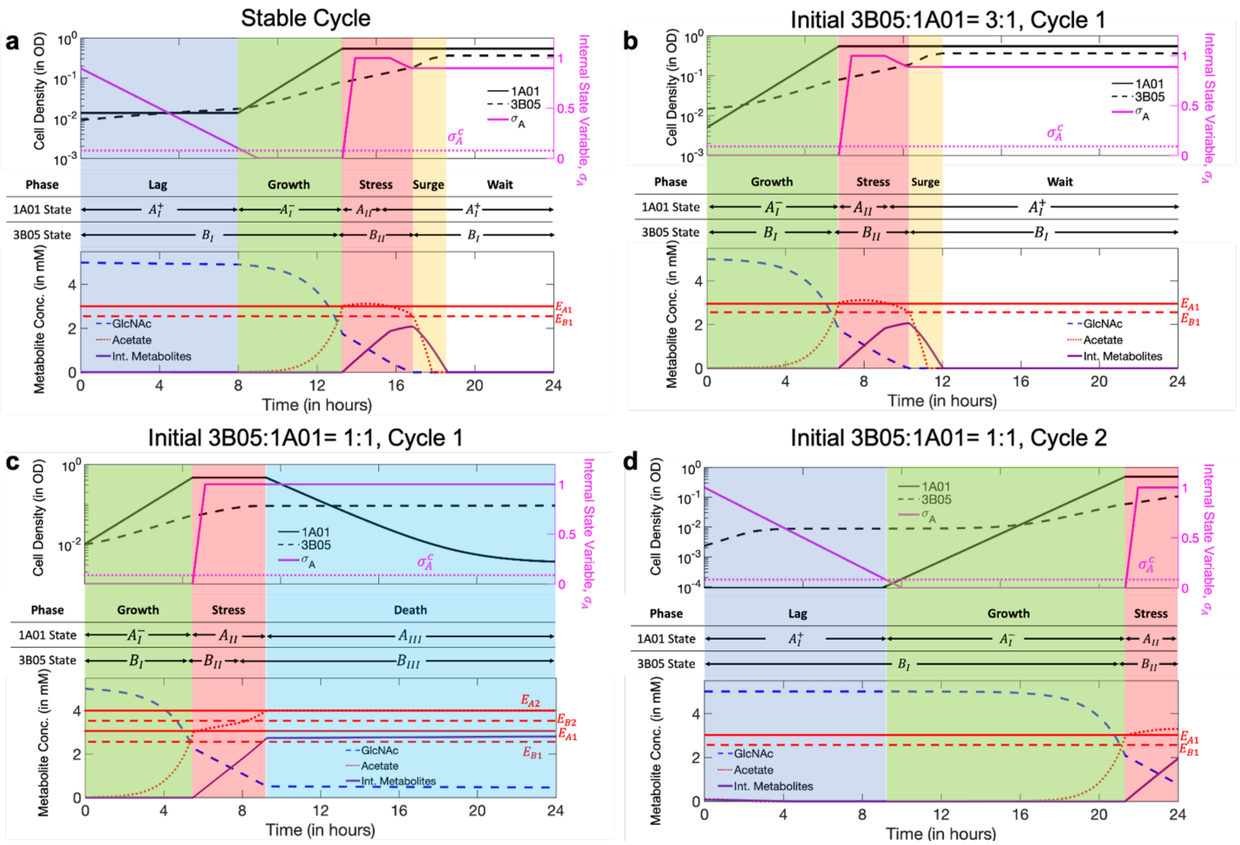


**Figure N4: Dynamics of cells and metabolites.** In each panel, densities of 1A01 and 3B05 are shown on the top, and the concentrations of GlcNAc, acetate, and acid-induced metabolites (taken to be the sum of pyruvate and lactate) are shown in the bottom. Different colors indicate the different phases of the co-culture: lag (purple), growth (green), stress (pink), surge (yellow), wait (white). **(a)** the stable cycle; **(b)** first cycle with initial inoculant of 3B05:1A01 = 3:1; **(c), (d)** first two cycles with initial inoculant of 3B05:1A01 = 1:1.

The final cycle attained for both 1:1 and 3:1 initial densities is the stable cycle depicted in **Fig. N4a**. This cycle illustrates the dynamic nature of the coexistence of 1A01 and 3B05. Starting from ~8 hours into the cycle, the system is in the "Growth Phase" (shaded green region) where both 1A01 and 3B05 grow at their normal rates (solid and dashed black lines, respectively, top plot), as there is abundant GlcNAc (dashed blue line, bottom plot); the pH is in the normal range (acetate, dotted red line bottom plot) below the threshold values indicated by the red horizontal lines above which growth stops; and the internal state of 1A01 ($\sigma_A$, solid magenta line in top plot) is below the threshold value ($\sigma_A^c$, dotted horizontal magenta line) above which growth stops. In this phase, 1A01 cells are in state $A_I^-$ and 3B05 cells are in state $B_I$ (as defined in **Extended Data Fig. 5a**).

As 1A01 grows rapidly in the "Growth Phase", acetate quickly accumulates in the coculture. This leads to the growth arrest of 1A01 (entering state $A_{II}$) when the acetate concentration exceeds the threshold $E_{A1}$. In this state, 1A01 continues to consume GlcNAc, turning it into metabolites which are then excreted. Additionally, the internal variable $\sigma_A$ increases rapidly, reflecting the loss of other metabolites due to acetate accumulation (**Fig. 3d-f**). This phase is labelled as the "Stress Phase" and shaded pink. Here 3B05 is in state $B_{II}$ and continues to grow, albeit slowly, assisted by the acetate-induced metabolites excreted from 1A01. The growth of 3B05 slowly removes acetate from the medium.

When the acetate concentration falls below the threshold $E_{B1}$, 3B05 is back in state $B_I$ and can resume normal growth. It grows even faster than in the Growth Phase due to the presence of acetate-induced metabolites from 1A01 (purple squares, **Extended Data Fig. 1f**). This phase is thus called the "Surge Phase" and is shaded as yellow. 1A01 is no longer inhibited by acetate either in this phase. But it cannot grow for two reasons: First, GlcNAc is depleted, and second, its internal variable $\sigma_A$ is large. It is in state $A_I^+$. Since acetate and acetate-induced metabolites are no longer produced by 1A01 in this phase (as GlcNAc is depleted), while 3B05 continues to consume them, eventually, 3B05 clears the acetate and stops growing. The co-culture stagnates at this point and enters the "Wait Phase" (not shaded).

The coculture remains in the "Wait Phase" until dilution into fresh medium in the new cycle. With the infusion of fresh GlcNAc, 1A01 starts its slow recovery by reducing the internal variable $\sigma_A$ while consuming GlcNAc, mimicking the replenishment of depleted metabolites inside the cell. 1A01 would remain in state $A_I^+$ as long as $\sigma_A > \sigma_A^c$ ("Lag Phase", purple shade). During this time, 1A01 continues to convert GlcNAc into acetate and acetate-induced metabolites, which support the growth of 3B05 (in state $B_I$). When 1A01 is finally able to grow again (when $\sigma_A$ drops below the threshold $\sigma_A^c$ ~8 hours later), the system enters the Growth Phase again and the cycle repeats.

While the experimental observations of stable cycle are explained by the dynamical model, we find that the model can even describe the approach to the stable cycle. We first explore the case with 3:1 initial ratio of 3B05 to 1A01 (**Fig. N4b**): In the first cycle, both 1A01 and 3B05 grow at their normal rates from time 0, when there is abundant GlcNAc and normal pH ("Growth Phase", green shaded region). This leads to acetate accumulation and stoppage of growth of 1A01 ("Stress Phase", pink shaded region) as described for the stable cycle in Panel a. The internal variable $\sigma_A$ also increases rapidly, indicating a loss of metabolites. The high initial ratio of 3B05 to 1A01 ensures that there is enough 3B05 to clear the acetate excreted by 1A01, until the point when GlcNAc is almost depleted, after which the coculture stagnates (Wait Phase, unshaded). Because the first cycle does not have a Lag Phase (as in the stable cycle shown in Panel a), the coculture has a prolonged Wait Phase. Starting from the second cycle, the Lag Phase occupies the first ~8 hours of the cycle, putting the coculture in the stable cycle.

We now explore the approach to the stable cycle of the 1:1 initial ratio of 1A01 and 3B05, which is more complex but still entirely described by the model. In the first cycle (**Fig. N4c**), 1A01 and 3B05 grow when there is abundant GlcNAc, and normal pH as described in panels a and b ("Growth Phase', green shaded region). However, the lower initial density of 3B05 makes it unable to clear acetate rapidly, so that 1A01 enters growth arrest with a lot of GlcNAc still remaining ("Stress Phase", pink shaded region). In the Stress Phase, 1A01 continues to consume GlcNAc and excrete acetate and internal metabolites. Because 3B05 grows slowly and hence consumes acetate slowly in the Stress Phase, acetate continues to accumulate, such that the system is pushed into very high acetate concentrations where 3B05 is in state $B_{III}$ and cannot grow, and 1A01 is in state $A_{III}$ and dies ("Death Phase", cyan shade). In this phase, the density of 1A01 drops rapidly. The cycle also ends with a very high value of the internal variable $\sigma_A$ indicating a severe loss of metabolites.

At the start of the second cycle for a 1:1 initial ratio of 1A01 and 3B05 (**Fig. N4d**), 1A01 stops dying as acetate is cleared out by dilution. However, due to the large value of $\sigma_A$ going into the cycle, 1A01 remains in state $A_I^+$ while its internal state recovers as long as $\sigma_A > \sigma_A^c$, while 3B05 grows on the lingering acetate from the previous cycle and the acetate-induced metabolites excreted by 1A01 as 1A01 is in state $A_I^+$ ("Lag Phase", purple shade). Once $\sigma_A < \sigma_A^c$, both 1A01 and 3B05 grow exponentially ("Growth Phase", green shade). This Growth Phase lasts much longer than the green phases described in Panels a-c as the starting density of 1A01 is very low, and thus it takes a long time for acetate to accumulate. ~21 hours after the start of the cycle, acetate accumulates enough to stop the growth of 1A01 and 3B05 ("Stress Phase", pink shade). Due to the prolonged period at very low densities, 1A01 is unable to consume all of the GlcNAc by the end of the cycle. Further, because of the short duration in Stress Phase in this cycle, 3B05 is unable to grow and clear the acetate accumulated during the Growth Phase by the end of the cycle. However, the stress at the end of 2nd cycle does lead to a high value of $\sigma_A$ and a significant Lag Phase in the subsequent (3rd) cycle, placing the coculture in the stable cycle.

From the detailed analysis of the above two examples, we see that if the initial ratio of 3B05 to 1A01 is high enough, it will enter a stable cycle provided that the duration of the cycle is long enough to contain the four crucial phases: Lag, Growth, Stress, and Surge. If the initial ratio is lower, then the coculture may be able to recover after 1A01 suffers death for a while.

| Crossfed Metabolites (Purple) | Remaining Metabolites (Grey) |
|---|---|
| Glycerate | 2-hydroxyglutarate |
| 1-amino-propan-2-one-3-phosphate | 2-aceto-lactate |
| 3-dehydroshikimate | Acetone |
| 4-amino-butyrate | Dyhydroxyphenylglycol |
| Coumaraldehyde | 2-methyl-maleate |
| 4-hydroxybenzoate | Glycine |
| 5-oxoproline | L-asparate-semialdehyde |
| Ribulose-5-phosphate | L-aspartate |
| Ascorbate | L-glutamate-semialdehyde |
| Glutamate | Acetyl-glutamate |
| Pyruvate | |

**Supplementary Table 1. Metabolites measured by untargeted metabolomics during the 'acetate peak' of the coculture. Fig. 2f** of the main text shows plots of the metabolite dynamics measured by untargeted metabolomics of the spent medium during the 'acetate peak' of the stable cycle. The cross-fed (purple lines in **Fig. 2f**) and the remaining metabolites (grey lines) are listed above. The determination of whether a metabolite was cross-fed or not is discussed in the Supplementary Methods.

**Supplementary Table 2. Result of FIA-TOF metabolomic measurement.** Metabolic features with their intensities for the samples described in **Fig. 2e,f** and **Extended Data Fig. 4c-j.**
